# Terpenes as Modulators of Nociceptive Signaling: Behavioral and Molecular Insights from *Caenorhabditis elegans*

**DOI:** 10.1101/2025.10.22.683951

**Authors:** Kaoutar Benkhraba, Fatma Boujenoui, Jesus D. Castaño, Jabin Sultana, Francis Beaudry

**Author notes:** Corresponding author: Francis Beaudry, Ph.D., Professor of Analytical Pharmacology, Canada Research Chair in metrology of bioactive molecules and target discovery, Département de Biomédecine Vétérinaire, Faculté de Médecine Vétérinaire, Université de Montréal, 3200 Sicotte, Saint-Hyacinthe, QC, Canada J2S 2M2.

## Abstract

Terpenes such as Limonene and β-Caryophyllene have demonstrated pain-modulating properties, potentially through interactions with TRPV1 receptors. This study examines the antinociceptive effects of four terpenes derived from *Cannabis sativa*: Limonene, β-Caryophyllene, α-Humulene, and α-Myrcene using *Caenorhabditis elegans* (*C. elegans*). The primary objective was to characterize terpene-induced modulation of nocifensive responses to noxious heat, and to elucidate their influence on molecular pathways via specific receptor targets. Thermotaxis assays quantified the antinociceptive activity of increasing terpene concentrations in wild-type nematodes. To assess receptor-specific mechanisms, assays were performed in mutant strains lacking functional OCR-2 and OSM-9 (TRPV-like vanilloid nociceptors), and NPR-19 and NPR-32 (encoding cannabinoid-like receptors). Proteomic profiling coupled with bioinformatics analysis identified terpene-induced alterations in signaling pathways and biological processes. All four terpenes exhibited significant antinociceptive activity in wild-type *C. elegans*, with impaired effects observed in vanilloid receptor mutants, implicating TRPV-like channels in their mechanism of action. Proteomic and pathway analyses revealed terpene-specific molecular signatures, highlighting differential modulation of neuronal and stress-responsive signaling cascades. By elucidating the molecular mechanisms underlying terpene-induced nociceptive modulation, this work strengthens the growing body of evidence supporting the therapeutic promise of terpenes in pain management outside the effect referred to as the “entourage effect.”

## 1. Introduction

Chronic pain is a prevalent, multifaceted, and burdensome condition that profoundly impacts individuals and society at large [1]. In contrast to acute pain, which functions as a vital warning signal of actual or potential tissue damage, chronic pain persists beyond the normal healing process and is increasingly recognized as a critical medical priority [2]. Its complexity spans physical symptoms as well as psychological and social dimensions, necessitating comprehensive, multidisciplinary treatment strategies [3]. Kosek et al. (2016) classified chronic pain into three primary categories: nociplastic pain, involving altered nociception without evident tissue or nerve damage; neuropathic pain, resulting from injury or disease of the somatosensory nervous system; and nociceptive pain, caused by ongoing tissue injury or inflammation [4]. Beyond its toll on individuals’ quality of life, chronic pain is a leading contributor to disability and poses a significant economic burden on global healthcare systems [5]. Treatment approaches typically include pharmacological agents such as nonsteroidal anti-inflammatory drugs (NSAIDs), anticonvulsants, antidepressants, and opioids, alongside psychological therapies and invasive interventions [6, 37–40]. However, many patients experience inadequate relief, and the escalating reliance on opioids has fueled a public health crisis marked by addiction and overdose fatalities [7, 41–43]. In light of these challenges, medical cannabis has emerged as a potential alternative for managing chronic pain, particularly in neuropathic and cancer-related cases. Research into the analgesic properties of cannabinoids, particularly tetrahydrocannabinol (THC) and cannabidiol (CBD), has expanded in response to the complex and heterogeneous nature of chronic pain, as well as the pressing need for safer alternatives to opioids [9,10]. Although phytocannabinoids such as THC and CBD show promising therapeutic potential, concerns persist regarding their long-term efficacy and associated psychotropic effects [11]. The so-called entourage effect, in which terpenes, aromatic and physiologically active components of cannabis, may modify or increase the pharmacological activities of cannabinoids, has drawn more attention as a result. [12]. Preclinical studies suggest that several terpenes, including myrcene, linalool, α-humulene, and β-pinene, may independently exert analgesic effects comparable to those of cannabinoids when administered in isolation [12, 44–46]. Although these compounds are widely present in cannabis and are thought to contribute to its synergistic effects, several studies have sought to differentiate them to better characterize their unique pharmacological properties. However, since most studies focus on terpene–cannabinoid combinations, isolating and characterizing the specific contributions of individual terpenes remains a major challenge [13, 30, 47]. Terpenes may have direct pharmacodynamic effects outside of cannabinoid receptor interaction, according to more recent research, especially through transient receptor potential (TRP) channels. A promising pharmaceutical target among these is the TRPV1 receptor, a non-selective cation channel implicated in pain perception and neurogenic inflammation [18, 68, 69]. Numerous terpenes produced from cannabis have been demonstrated to interact with TRPV1 either as allosteric modulators, such myrcene, α-pinene, limonene, and linalool, or as direct agonists, like β-caryophyllene and camphor [50, 51, 52]. Specifically, it has been demonstrated that myrcene, one of the most prevalent monoterpenes in *Cannabis sativa*, both directly and allosterically activates TRPV1, causing calcium influx in neuronal cells [49]. According to this dual mechanism, myrcene may have analgesic effects independently of cannabinoid receptor pathways by modulating nociceptive transmission through TRPV1. However, because most research focuses on intricate mixes or co-administration techniques, it is still difficult to distinguish the distinct contributions of particular terpenes, even with the increased interest in terpene– cannabinoid synergy. A thorough assessment of the effects of isolated terpenes in preclinical and clinical models is essential to completely clarify the medicinal potential of cannabis-derived terpenes beyond THC and CBD [49].

Recent studies have increasingly adopted *Caenorhabditis elegans* (*C. elegans*) as a model organism to investigate the molecular mechanisms underlying TRPV-mediated pain, due to its compact nervous system comprising 302 neurons and well-mapped neural circuitry. This model is particularly valuable because it expresses homologous TRPV channels, OSM-9 and OCR-2, which share structural and functional characteristics with mammalian TRPVs, including TRPV1. For example, OSM-9 and OCR-2 are essential for thermal nociception in *C. elegans*, as demonstrated by the reduced avoidance of noxious heat in mutants lacking these genes [15,16]. Functional conservation is further highlighted by the restoration of capsaicin sensitivity when TRPV1 is ectopically expressed in ASH sensory neurons of *C. elegans*, an effect absent in *osm-9* or *ocr-2* mutants [15]. Analogous to the TRPV1 mainly located on nociceptor terminals in vertebrates, OSM-9 and OCR-2 assemble into heteromeric channels that localize specifically to sensory cilia [15,16]. Moreover, recent findings reveal that the nocifensive response of wild-type *C. elegans* to thermal stimuli (32–35 °C) is significantly hindered following prolonged exposure to capsaicin or other vanilloids, a phenotype that reverses approximately six hours post-exposure [17]. This transient desensitization parallels that of TRPV1 in mammals following sustained agonist stimulation [30, 61]. Importantly, OCR-2 appears to mediate this response, as *ocr-2* mutants fail to exhibit heat avoidance attenuation, highlighting the evolutionary conservation and functional importance of this channel in nociceptive processing [17]. Leveraging *C. elegans* enables researchers to evaluate desensitization kinetics, dissect downstream signaling pathways, and elucidate terpene–TRPV interactions with molecular precision, all within an *in vivo* system that complements mammalian models. This approach not only deepens our understanding of terpene-mediated modulation of chronic pain but also provides a powerful platform for the objective screening of therapeutics targeting TRPV1-like channels.

We hypothesized that terpenes exert antinociceptive effects in *C. elegans* through interactions with vanilloid receptor homologs OSM-9 and OCR-2. To investigate this, the objectives of the current study were: (1) to characterize terpene exposure–response relationships using thermal avoidance assays; (2) to identify the specific receptor targets mediating these effects; and (3) to elucidate the proteins and signaling pathways underlying terpene-induced antinociception via mass spectrometry-based proteomic analysis.

## 2. Materials and methods

### 2.1 Chemicals and reagents

All chemicals and reagents were sourced from Fisher Scientific (Fair Lawn, NJ, USA) or Millipore Sigma (St. Louis, MO, USA). Capsaicin was obtained from Toronto Research Chemicals (North York, ON, Canada), while Limonene, α-Myrcene, α-Humulene and β-Caryophyllene were procured from Toronto Research Chemicals (Vaughan, Ontario, CAN), now LGC Standards.

### 2.2 *C. elegans* strains

The N2 (Bristol) strain of *C. elegans* was used as the reference strain. Mutant strains included ocr-2 (123), and osm-9 (CX-10). The N2 (Bristol) strain and all mutants were obtained from the Caenorhabditis Genetics Center (CGC) at the University of Minnesota (Minneapolis, MN, USA). Strains were handled and maintained in accordance with the previously mentioned standard methods [18, 19]. Nematodes were cultured on nematode growth medium (NGM) agar and kept at 22°C in a Thermo Scientific Heratherm refrigerated incubator (Fair Lawn, NJ, USA). Unless otherwise specified, all experiments were conducted at room temperature (∼22°C).

### 2.3 *C. elegans* pharmacological manipulations

Limonene, α-myrcene, α-humulene, and β-caryophyllene stock solutions were made at a 50 µM concentration in 0.01 % Tween 20. Following homogeneity-ensuring vortexing, successive dilutions were carried out in the same Tween-20 vehicle to achieve final concentrations of 20, 10, 5, 1, and 0.2 µM. Using the method outlined by Margie et al. [22], synchronously age-matched *C. elegans* nematodes were kept on NGM agar plates (92 × 16 mm) for 72 hours until they reached maturity. *C. elegans* were taken off of food on the day of the experiment and submerged in 7 mL of the corresponding terpene solutions. Due to partial absorption by the agar, the liquid produced a thin layer (∼2–3 mm height), which allowed the nematodes to freely float. Before behavioral testing, nematodes were extracted and washed three times with S-Basal buffer after a 60-minute exposure. Before testing, a parallel cohort was cleaned and kept on terpene-free NGM agar for an extra six hours to evaluate any possible aftereffects.

### 2.4 Thermal avoidance assays

Our approach to evaluating heat avoidance in this paper was adapted from the four-quadrant system that was previously outlined [20] and utilized in earlier, successfully published articles [17, 21]. In short, 92 × 16 mm petri dishes that were partitioned into four quadrants were used for the research. A central circle, measuring one centimeter in diameter, delineated a region that was not taken into account by *C. elegans*. Four quadrants of petri dishes were created: two control regions (B and C) and two stimulation areas (A and D). The nematodes (usually 100 to 500 young adults per petri dish) were paralyzed in each quadrant using sodium-azide (0.5 M). An electrically heated steel edge (0.8 mm in diameter) was used to generate toxic heat, resulting in a radial temperature gradient (e.g., 32 35°C on NGM agar 2 mm from the tip detected using an infrared thermometer). Within 30 minutes, the plates were kept at 4°C for a minimum of one hour, and the number of *C. elegans* in each quadrant was counted. Nematodes that stayed inside the inner circle were not taken into account. The following equation (*TI* = [(*A* + *D*) − (*B* + *C*)] / (*A* + *B* + *C* + *D*)) displays the thermal avoidance index (TI). Additionally, Figure S1 provides details. Each evaluated *C. elegans* experimental group was phenotyped using both the TI and the animal avoidance percentage. Previous investigations served as the basis for the stimulus temperature that was employed [23].

### 2.5 Sample preparation for proteomics

Before being collected and thoroughly cleaned with S Basal, cultured nematodes (with or without terpenes exposure) were collected in liquid medium and centrifuged at 1,000 g for 10 minutes. The nematodes were reconstituted in a phosphate buffered saline solution (137 mM NaCl, 2.7 mM KCl, 10 mM Na_2_HPO_4_, and 1.8 mM KH_2_PO_4_) that contained 1% (v/v) of the protease inhibitor cocktails cOmpleteTM and Triton X-100 (Roche Diagnostic Canada, Laval, QC, Canada). In reinforced 1.5 mL homogenizer tubes with 25 mg of 500 μm glass beads, the suspension was poured. A Fisher Bead Mill Homogenizer was used to homogenize the samples in five 60-second bursts at a speed of 5 m/s. For ten minutes, the homogenates were centrifuged at 12,000 g. The Bradford test was used to measure the protein content of each homogenate. The ice-cold acetone precipitation method (1:5, v/v) was used to isolate proteins (∼100 μg). To enhance protein breakdown, the resultant protein pellet was dissolved in 100 μL of 50 mM TRIS-HCl buffer (pH 8), combined with a Disruptor Genie running at maximum speed (2,800 rpm) for 15 minutes, and then sonicated. Using a heated reaction block, we incubated the solution at 120°C for 10 minutes to denature the protein. For fifteen minutes, the solution was let to cool. Proteins were alkylated with 40 mM iodoacetamide (IAA) at room temperature for 30 minutes while being shielded from light and reduced with 20 mM dithiothreitol (DTT) for 15 minutes at 90°C. Proteomic-grade trypsin (5 μg) was then added, and the reaction was run for 24 hours at 37°C. The addition of 10 μL (∼ 10% v/v) of a 1% trifluoroacetic acid (TFA) solution halted the digestion. Following a 10-minute centrifugation at 12,000 g, 100 μL of the supernatant was poured into HPLC injection vials for examination.

### 2.6 Proteomic analysis

Online chromatographic separation was performed using a Thermo Scientific Vanquish Neo UHPLC system (San Jose, CA, USA) configured in trap-and-elute mode within a two-dimensional nano-LC system. The setup included a Thermo Scientific PepMap Neo 5 μm C18 300 μm × 5 mm trap cartridge and a Thermo Scientific PepMap Neo C18 2 μm × 75 μm × 500 mm nano column. A 2 μL sample (∼2 μg of digested proteins) was loaded onto the trap column and desalted with solvent A [0.1% formic acid in water] at a 20 μL/min flow rate for 0.5 minutes. Peptide elution was carried out using a linear gradient of 5% to 35% solvent B [80% acetonitrile, 20% water, 0.1% formic acid] over 135 minutes, followed by a 2-step column wash at 50% solvent B for 10 minutes and 80% for 10 minutes. The flow rate was maintained at 200 nL/min. Afterwards, the column was equilibrated to the initial solvent composition (5% solvent B) by running 10 column volumes.

Data acquisition was conducted on a Thermo Scientific Q Exactive Plus Orbitrap Mass Spectrometer (San Jose, CA, USA) equipped with a Nanospray Flex ion source. Data collection was performed in positive ion mode with a nanospray voltage of 2.2 kV, and the ion transfer tube was maintained at 200°C. The mass spectrometer was operated in data-dependent acquisition (DDA) mode, utilizing a TOP-12 acquisition strategy. A high-resolution MS^1^ survey scan (m/z 375–1200) was acquired at a resolution of 70,000 (FWHM), with an automatic gain control (AGC) target value of 1 × 10⁶ and a maximum injection time of 100 ms. The 12 most intense precursor ions exceeding an intensity threshold of 1 × 10⁴ were selected for fragmentation via higher-energy collisional dissociation (HCD) at a normalized collision energy (NCE) of 27. MS^2^ spectra were acquired at a resolution of 17,500 (FWHM) using an AGC target value of 2 × 10⁵ and a maximum injection time of 100 ms.

### 2.7 Protein Quantification

Protein identification and quantification were performed using FragPipe (version 22.0) with the MSFragger search engine. Data processing was conducted using default settings. The raw MS^1^ and MS^2^ spectra were analyzed using *C. elegans* reference proteome from UniProt (taxon identifier 6239). Database search parameters included a precursor mass and fragment mass tolerance of 20 ppm. Trypsin was selected as the digestion enzyme, allowing a maximum of two missed cleavages. Fixed modifications included carbamidomethylation (Cys), while variable modifications accounted for oxidation (Met), N-terminal acetylation, and phosphorylation (Ser, Thr, and Tyr). Minimum peptide length was set at 7 during the analysis. False discovery rate (FDR) filtering was performed using Percolator, with an FDR threshold of 1%. Relative quantification was conducted using a label-free approach, integrating peak intensities with ionQuant default options within FragPipe, and analyzing MaxLFQ intensity with FragPipe Analyst.

### 2.8 Bioinformatics

Differential protein expression analysis was conducted using datasets generated by FragPipe (version 22.0) and its online tool FragPipe-Analyst Extracted parameters included the log₂ abundance ratio (experimental group vs. N2 control), corresponding p-values, p-adjusted values, and protein accession identifiers. Functional enrichment analysis was performed using ClusterProfiler, incorporating Gene Ontology (GO) biological process (BP) annotation (Figure 8-A,B) and Reactome pathways (Figure 8-C,D) [51]. UMAP plots were produced using the uwot package (uwot:, 2025) in the software R (version 4.4.1).

### 2.9 Statistical analysis

Behavioral data were analyzed using the non-parametric Kruskal–Wallis test, followed by Dunn’s post hoc test for multiple comparisons. A significance threshold was set at p ≤ 0.05. All statistical analyses were performed using GraphPad Prism (version 10.6.0).

## 3. Results and discussion

In this study, we began by using our established experimental protocol to assess potential behavioral bias in *C. elegans*. Specifically, we evaluated the motility and quadrant preference of wild-type (N2) and select mutant strains, both in the presence and absence of α-Myrcene, α-Humulene, β-Caryophyllene, or Limonene. For all experiments, the petri quadrants were all maintained at room temperature (approximately 22 °C). None of the experimental groups exhibited significant quadrant selection bias following terpene exposure (e.g., 50 µM) (Figure 2). Thirty minutes after initial placement at the center of the Petri dish, nematodes remained evenly distributed, showing no preference for any specific quadrant. So, 60-minute exposures to Limonene, α-Myrcene, α-Humulene, or β-Caryophyllene did not produce observable behavioral biases or significantly affect overall motility.

**Figure 1:**
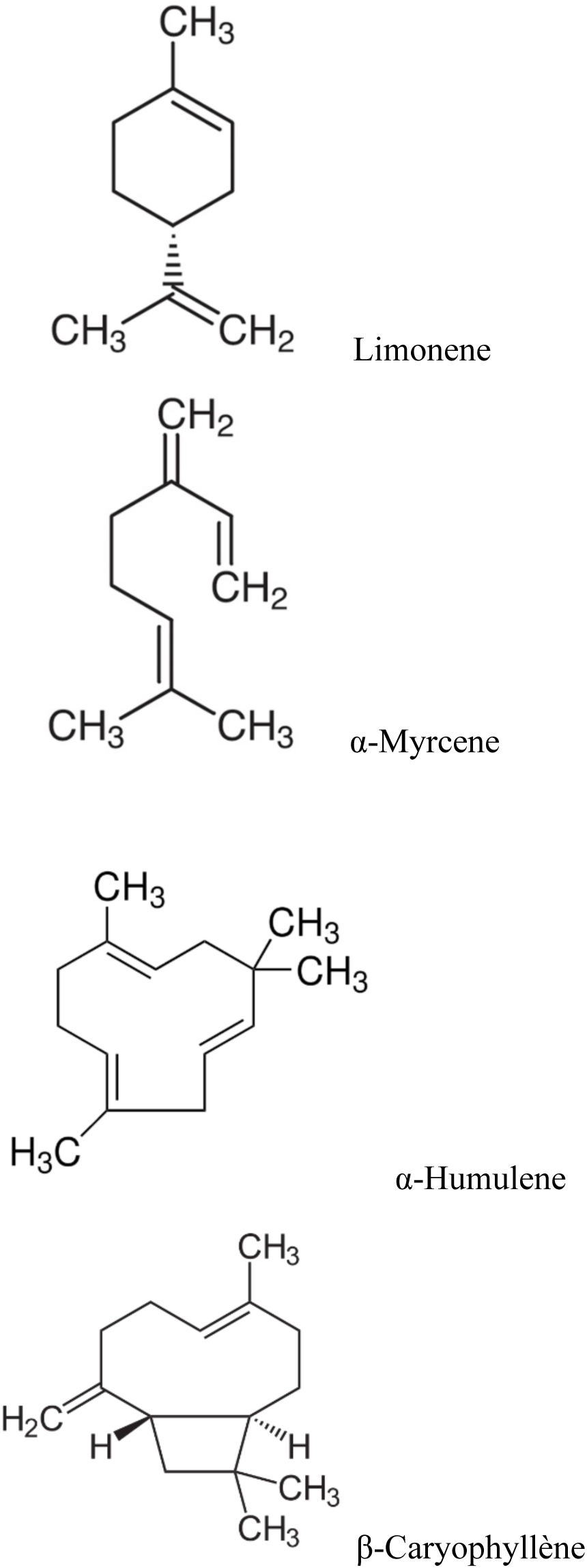
Molecular structure of Limonene, α-Myrcene, α-Humulene and β-Caryophyllene. Chemical structures of the four selected terpenes commonly found in cannabis and other botanical sources. Limonene is a cyclic monoterpene, α-myrcene is an acyclic monoterpene, while α-humulene and β-caryophyllene are sesquiterpenes characterized by bicyclic and tricyclic frameworks, respectively [35, 36].

**Figure 2:**
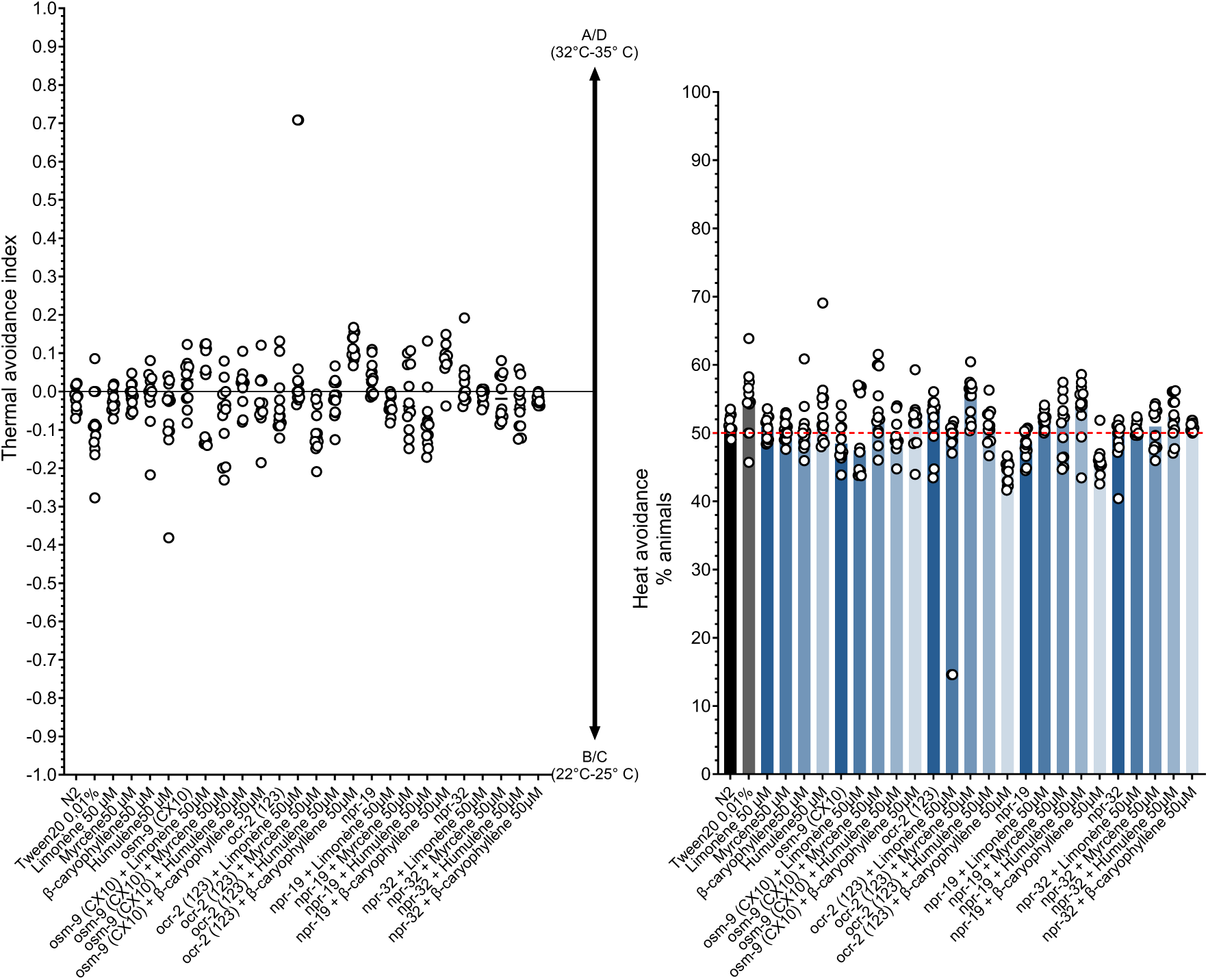
Assessment of Baseline Quadrant Bias in WT and Mutant *C. elegans* Exposed to Terpenes Under Non-Stimulating Conditions (22°C). WT (N2) and specific mutant nematodes’ motility and bias are compared on plates that are separated into four quadrants. Without a stimulation (negative control), the plates were maintained at a steady temperature of 22°C. All *C. elegans* genotypes evaluated with or without Limonene, α-Myrcene, α-Humulene or β-Caryophyllene showed no signs of quadrant selection bias.

### 3.1 Evaluation of antinociceptive activity of the terpenes

It is important to note that given the lipophilic nature of terpenes, their solubility in aqueous media is inherently limited. A non-ionic surfactant, Tween 20, was added at a low concentration (0.01% w/v) to guarantee adequate dispersion and provide homogenous working solutions appropriate for *C. elegans* bioassays. This concentration was determined to provide for adequate terpene emulsification while minimizing any possible harmful or behavioral impacts on the nematodes. N2 wild-type nematodes exposed to Tween 20 (0.01%) without terpenes were included as an additional control to account for any potential effects of the surfactant alone. This experimental design incorporated two essential control groups: (1) an untreated baseline group (no exposure) and (2) a vehicle control to isolate the influence of the Tween 20. This dual-control strategy strengthens the interpretability and reliability of the behavioral data by ensuring that any observed effects following terpene treatment can be attributed specifically to the bioactive compounds, rather than confounding solvent-related artifacts.

As previously noted, the therapeutic potential of terpenes in pain management remains insufficiently characterized. To address this gap, we evaluated the antinociceptive properties of Limonene, α-Myrcene, α-Humulene, and β-Caryophyllene. Following one hour of exposure, α-Myrcene and Limonene elicited concentration-dependent antinociceptive effects in *C. elegans*, as illustrated in Figures 3A and 3B. After treatment, nematodes were thoroughly rinsed and transferred to NGM agar plates maintained at 22 °C, then incubated for six hours to assess residual or latent effects. Subsequent thermal avoidance assays revealed that both monoterpenes continued to modulate nocifensive behavior, indicating sustained antinociceptive activity. Notably, α-Myrcene produced significant effects at higher concentrations (20 μM and 50 μM), while Limonene retained efficacy at concentrations as low as 1 μM. These observations suggest that the prolonged action of these compounds may be related to their strong lipophilic character. The high octanol–water partition coefficients (log P) of Limonene (4.233) [54] and α-Myrcene (4.22) [55], in particular, suggest a significant affinity for lipid environments and facilitate quick absorption into biological membranes. This physicochemical characteristic probably helps them maintain their pharmacologic effect by staying attached to intracellular compartments or tissue membranes beyond the exposure time. Figure 4A and 4B shows the concentration-dependent antinociceptive effects of the sesquiterpenes (α-Humulene [log P ∼5.16] and β-Caryophyllene[log P ∼6.11]) [56, 57] following a 1 h of exposure. Following the exposure, the nematodes were thoroughly washed as in the monoterpenes test. Comparable to α-Myrcene, exposure to α-Humulene and β-Caryophyllene (50, 20 μM for α-Humulene and up to 10 μM for β-Caryophyllene) still impaired *C. elegans*’ thermal avoidance response after six hours. These findings are consistent with the monoterpenes. Their pronounced lipophilicity may underlie the sustained effects observed following exposure [24]. In *C. elegans*, cannabinoid-like receptors (NPR-19 and NPR-32) are G-protein coupled receptors with limited neuronal expression, whereas vanilloid receptors (OSM-9 and OCR-2) function as nociceptors that are mostly found at the peripheral ends of sensory neurons [15,27]. Compounds must enter the nematode’s nervous system either by ingestion in the surrounding media or by absorption through the cuticle in order to interact pharmacologically with cannabinoid-like receptors. However, in the case of receptors like OSM-9 and OCR-2, which are expressed in environmentally exposed sensory neurons, direct interaction with external compounds is also possible. Interestingly, *C. elegans* primarily store lipids in the hypodermal layer and intestinal cells, tissues analogous to vertebrate skin [26, 28]. Given their high lipophilicity, terpenes are likely sequestered within these fat stores, which may act as depots enabling prolonged receptor engagement and sustained local release [26, 27].

**Figure 3:**
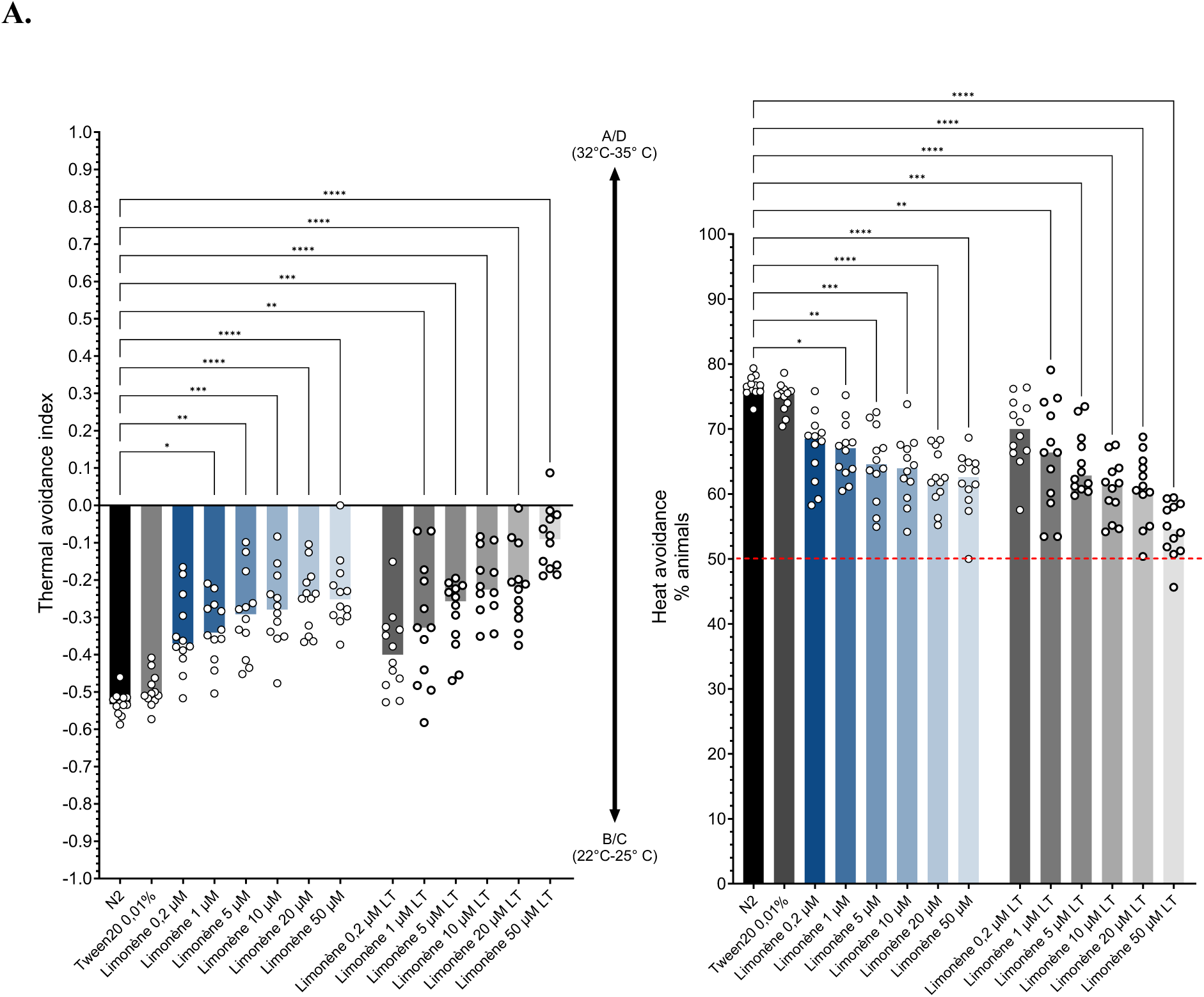
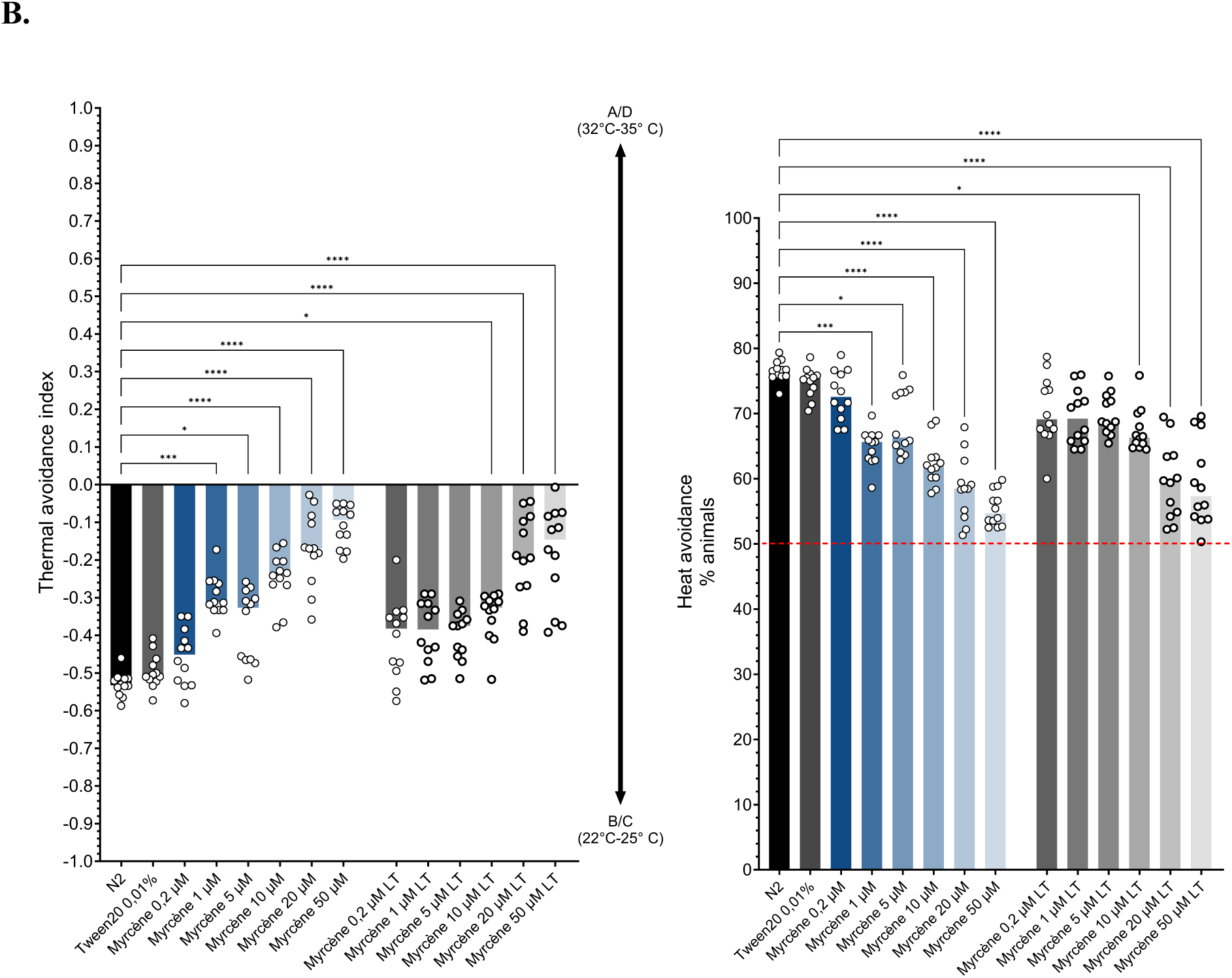
Evaluation of the pharmacological effects of monoterpenes (α-myrcene and limonene) on thermal avoidance in *C. elegans*. Nematodes were exposed to α-myrcene or limonene for 60 minutes prior to behavioral testing. Individual data points and group medians are shown, based on a minimum of 12 independent experiments per treatment group. Results indicate that both compounds significantly impaired thermal avoidance behavior in *C. elegans*. Statistical significance was assessed using the non-parametric Kruskal–Wallis test followed by Dunn’s post hoc test for multiple comparisons (n = 100 to 500 nematodes per Petri dish).

**Figure 4.**
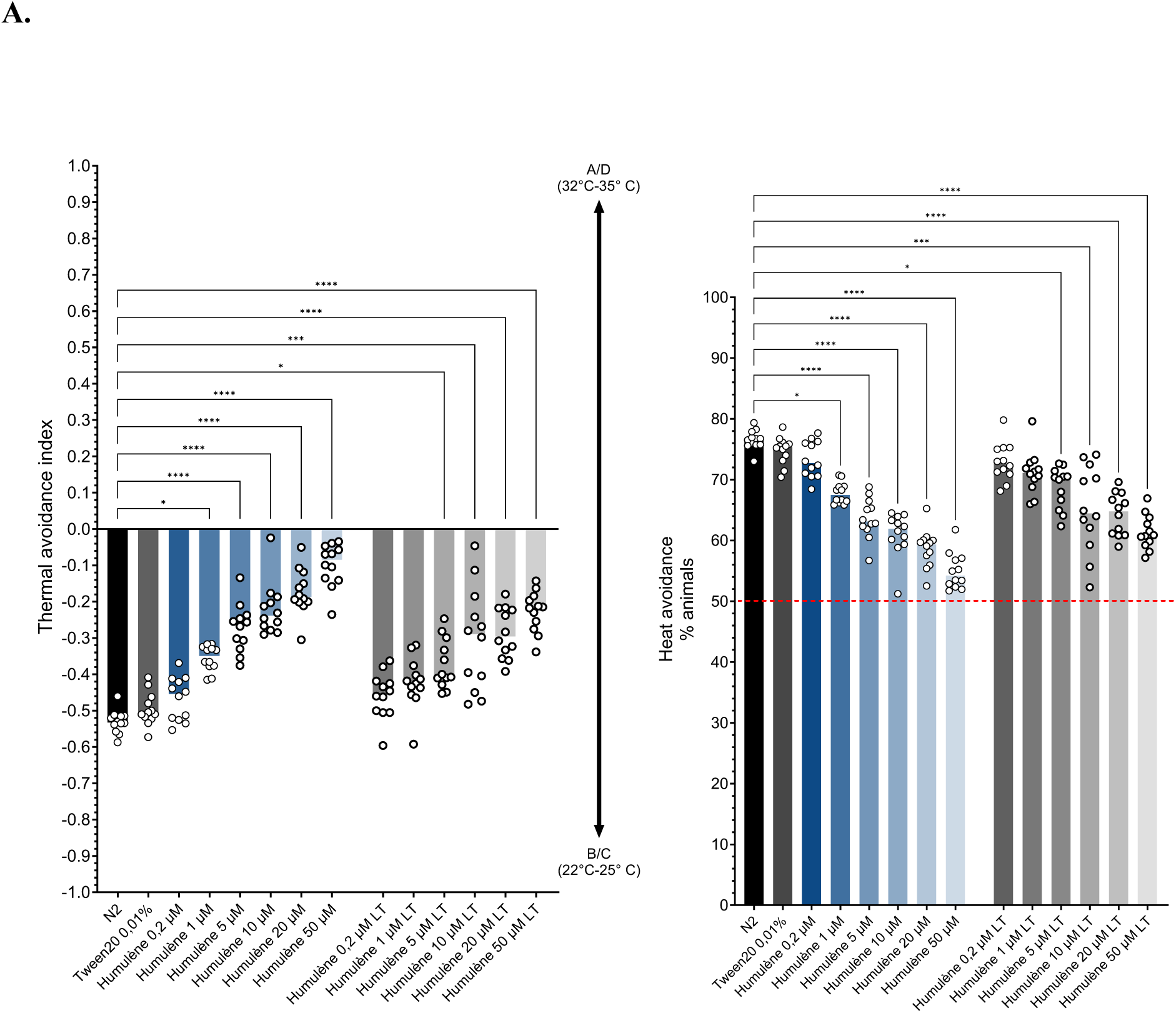
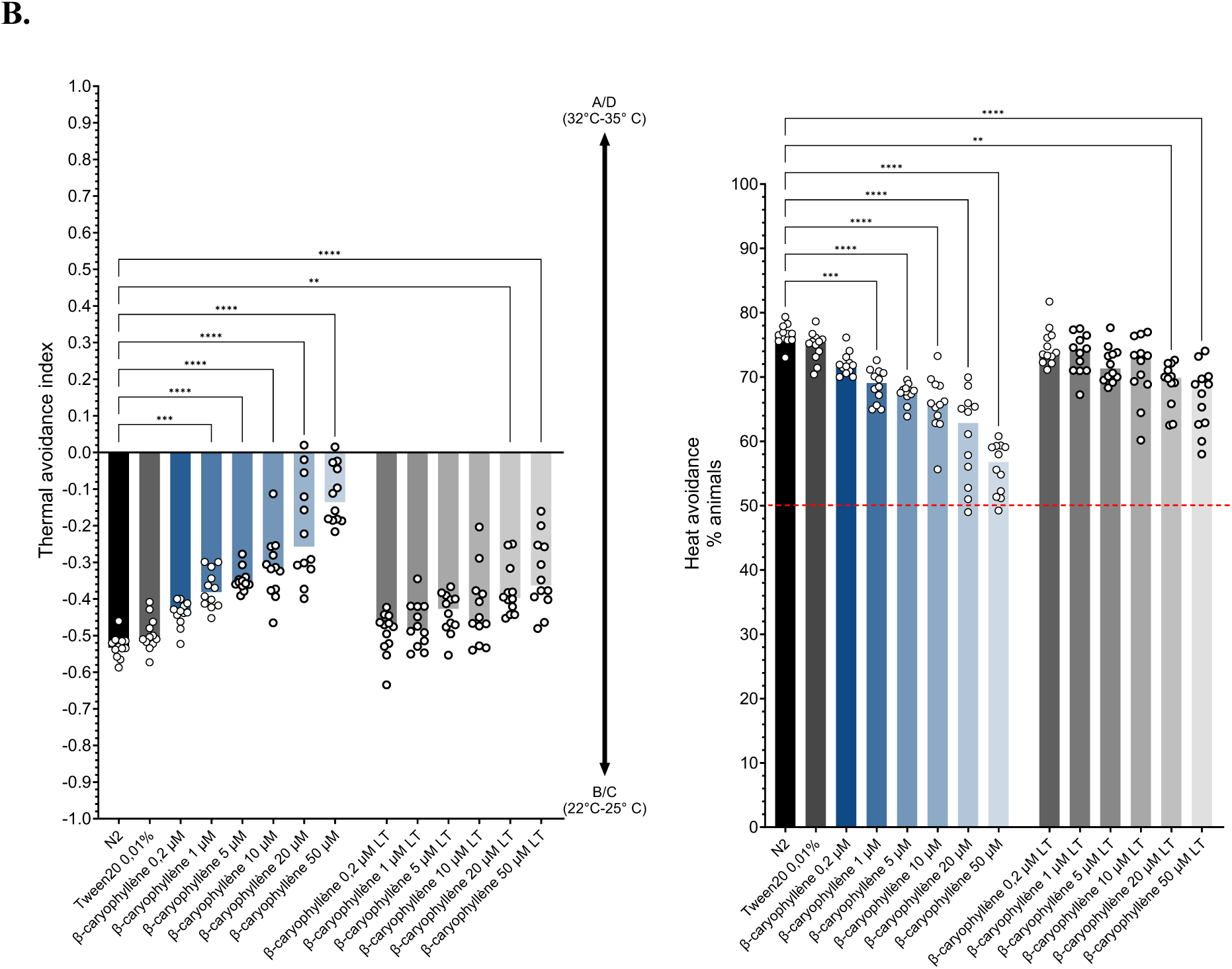
Pharmacological evaluation of α-Humulene and β-Caryophyllene on Thermal Avoidance Behavior in *C. elegans*. Nematodes were exposed to α-myrcene or limonene for 60 minutes prior to behavioral testing. Individual data points and group medians are shown, based on a minimum of 12 independent experiments per treatment group. Results indicate that both compounds significantly hindered thermal avoidance behavior in *C. elegans*. Statistical significance was assessed using the non-parametric Kruskal–Wallis test followed by Dunn’s post hoc test for multiple comparisons (n = 100 to 500 nematodes per Petri dish).

### 3.2 Identification of Limonene, α-Myrcene, α-Humulene and β-Caryophyllene’s targets

Identifying the molecular targets of compounds such as Limonene, α-Myrcene, α-Humulene, and β-Caryophyllene is essential for elucidating the mechanisms underlying the behavioral changes induced by terpene exposure. A substance typically exhibits pharmacological activity only when it binds to a significant fraction of its target receptors, producing a measurable physiological or phenotypic response. Thus, understanding receptor-ligand interactions is essential to grasp a drug’s potency and selectivity. In order to determine whether the observed terpene-induced effects are mediated through these conserved molecular pathways, we used *C. elegans* genetic models lacking functional receptor homologs, specifically the vanilloid-type TRP channel mutants (*ocr-2* and *osm-9*) and putative cannabinoid receptor mutants (*npr-19* and *npr-32*). Prior phenotyping analysis, *C. elegans* mutants were treated to each of the four terpenes separately for 60 minutes at a dosage of 50 μM. The *ocr-2* and *osm-9* mutants showed no discernible effects (p > 0.05), as shown in Figure 5A and B. These findings imply that the four terpenes are targeting homologs of the vanilloid receptor in *C. elegans*. The ability of *C. elegans* to detect and react to damaging heat stimuli is partially compromised when vanilloid receptor genes, including *ocr-2* and *osm-9*, are genetically ablated. In line with their function in modulating heat-induced avoidance responses, this loss of function in important TRPV homologs results in decreased nocifensive behavior and lowered thermal sensitivity [15, 29]. These phenotypic deficiencies provide credence to the crucial role that vanilloid-type TRP channels play in heat nociception pathways in both higher species and nematodes. Vanilloid receptor mutants (*ocr-2*, *osm-9*) in *C. elegans* have reduced sensitivity to harmful heat, as seen in Figure 5A and B.

**Figure 5.**
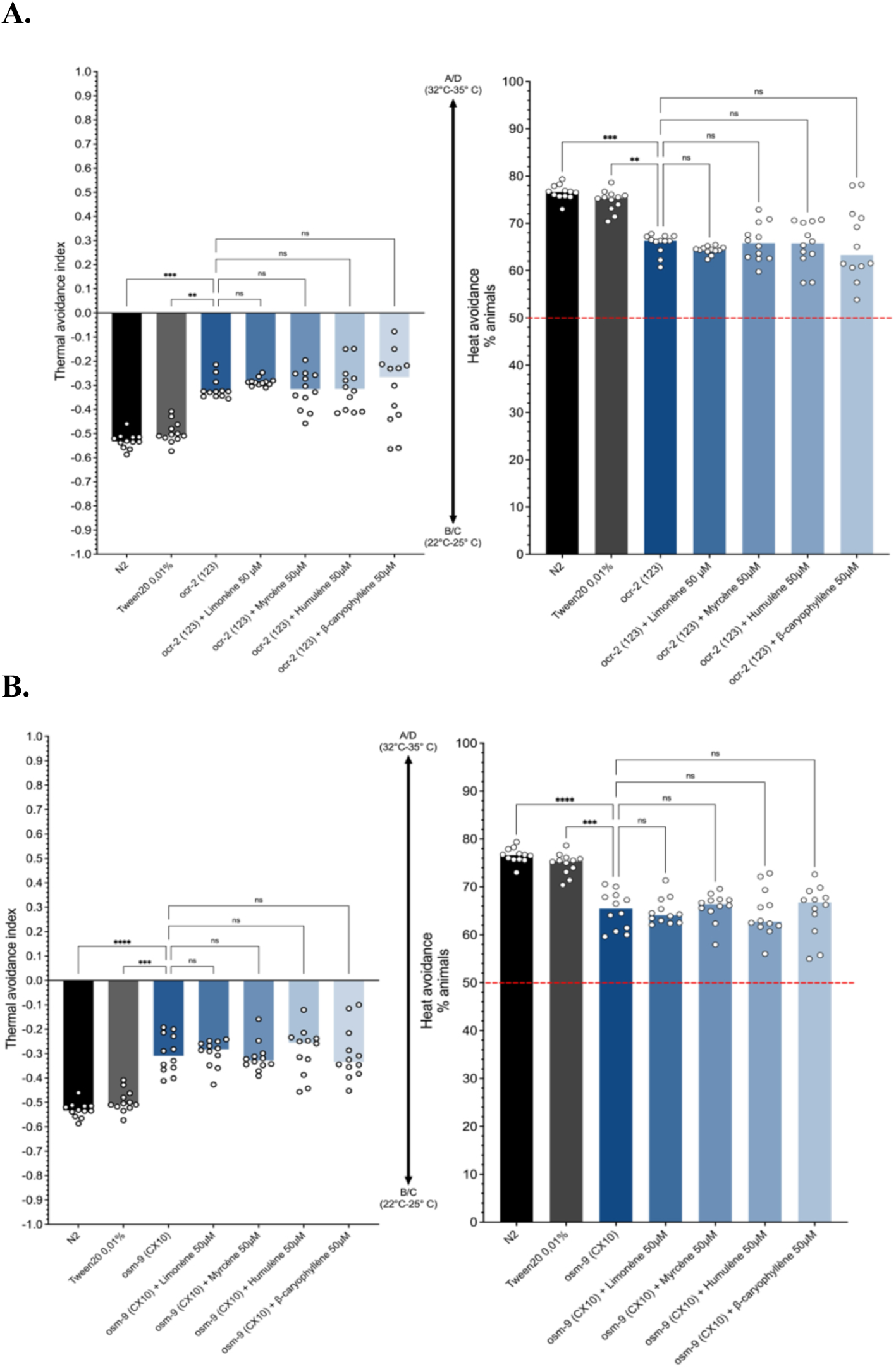
Identification of vanilloid receptor homologs mediating the antinociceptive effects of β-caryophyllene, α-myrcene, α-humulene, and limonene. Thermal avoidance behavior was assessed in C. elegans mutants lacking either ocr-2 (A) or osm-9 (B) to identify the vanilloid receptor homologs involved in terpene-induced antinociception. Data represent individual values and medians from a minimum of 12 independent experiments per treatment group. Results indicate that all four terpenes likely exert their effects through OCR-2 and OSM-9 channels. Statistical analysis was performed using the non-parametric Kruskal–Wallis test followed by Dunn’s post hoc test for multiple comparisons (n = 100 to 500 nematodes per Petri dish).

In *npr-19* and *npr-32* mutants, the antinociceptive effects of monoterpenes (Limonene and α-Myrcène) were considerable, as shown in Figure 6A and 6B. These findings imply that the cannabinoid receptor homologs *npr-19* and *npr-32* in *C. elegans* are not being targeted by these two terpenes. While both sesquiterpenes tested (β-caryophyllene and α-humulene) appear to exert part of their effects via vanilloid receptors (such as OSM-9 and OCR-2), our observations also suggest a potential involvement of cannabinoid receptor homologs NPR-19 and NPR-32 in *C. elegans*. Given that these terpenes are lipophilic and that β-Caryophyllene has a documented pharmacological interaction with CB2 receptors in animals [30], this dual receptor route seems especially likely. NPR-19 can be detected in a limited number of neurons that modulate behavioral outputs and has been suggested as a functional counterpart of mammalian cannabis receptors [27]. Similarly, endocannabinoid-like signaling and neuropeptidergic control have been linked to NPR-32 [31]. The notion that sesquiterpenes can modulate nociception and heat-avoidance behavior in nematodes not only via TRPV homologs but also through cannabinoid-like GPCR pathways is supported by the altered responses of *npr-19* and *npr-32* mutants to these terpenes. These findings are particularly compelling, as they suggest that terpenes may modulate multiple receptor systems, attenuating nocifensive responses to noxious heat through both cannabinoid and vanilloid receptor pathways. This dual-receptor targeting presents promising avenues for the development of innovative therapies for chronic pain, emphasizing the potential of compounds capable of simultaneously modulating vanilloid and cannabinoid signaling [32]. Furthermore, the human endocannabinoid, opioid, and vanilloid systems exhibit extensive crosstalk and complex regulatory interplay. Collectively, these interconnected networks govern key aspects of pain perception, emotional regulation, and physiological homeostasis [33,34]. Advancing our understanding of these multidimensional interactions may provide a critical framework for designing more effective multimodal analgesic strategies.

**Figure 6.**
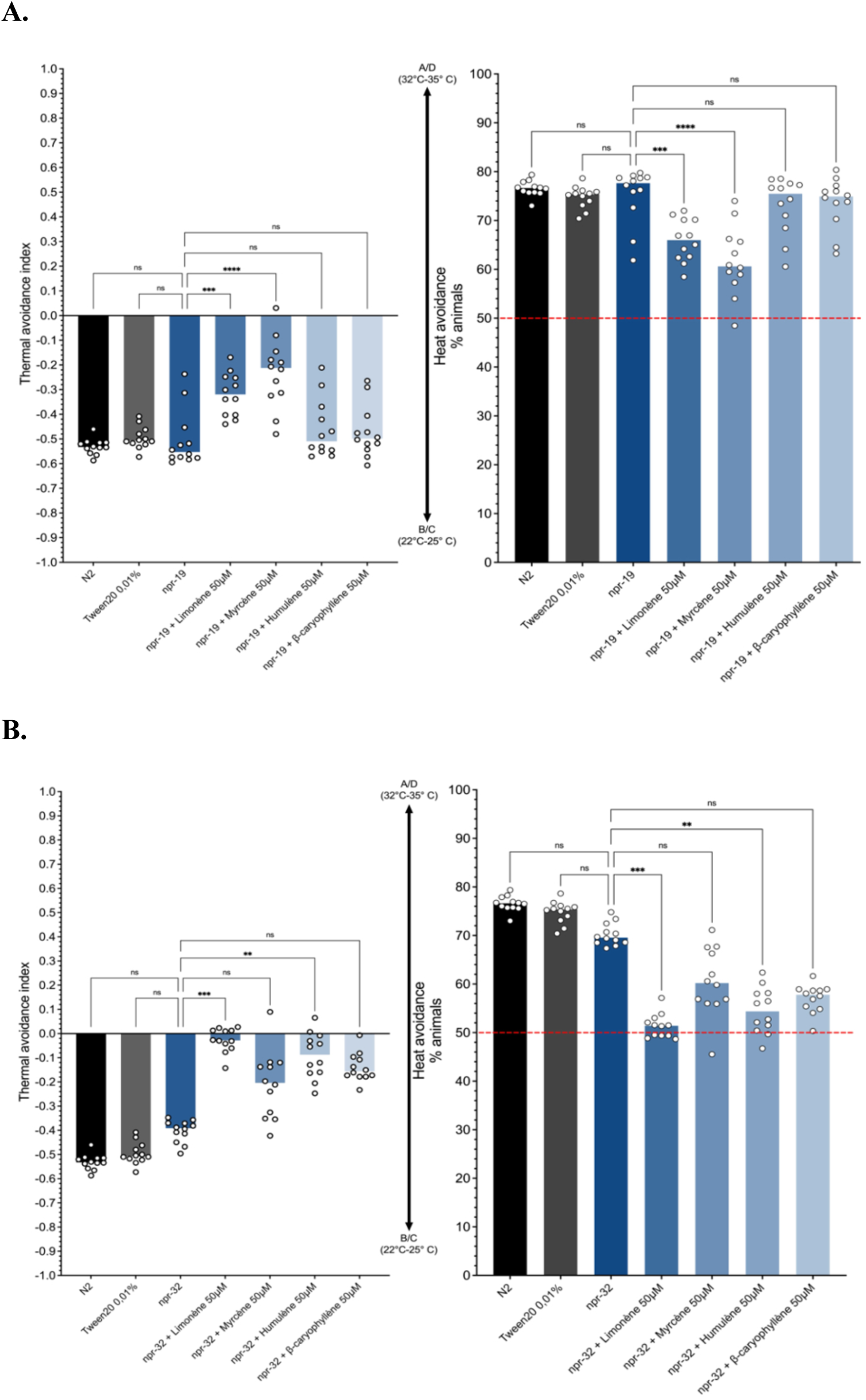
Identification of cannabinoid receptor homologs mediating the antinociceptive effects of β-caryophyllene, α-myrcene, α-humulene, and limonene. Thermal avoidance behavior was evaluated in *C. elegans* mutants deficient in either npr-19 (A) or npr-32 (B) to investigate the involvement of cannabinoid receptor homologs in terpene-induced antinociception. Data represent individual values and group medians from a minimum of 12 independent experiments per treatment condition. The results indicate that α-humulene and β-caryophyllene exert their antinociceptive effects, at least in part, through NPR-19 and NPR-32 signaling pathways. Statistical analysis was conducted using the non-parametric Kruskal–Wallis test followed by Dunn’s post hoc test for multiple comparisons (n = 100 to 500 nematodes per Petri dish).

### 3.3 Proteomic and bioinformatic investigations

Each terpene causes a different translational response in *C. elegans*, as evidenced by the global proteomic changes observed after exposure to four distinct terpenes (Limonene, α-Myrcene, α-Humulene and β-Caryophyllene) under heat-induced stress conditions, which are depicted in Figure 7 through combined dimensionality reduction and clustering analyses (PCA, hierarchical heatmap, and UMAP), reflecting differences in both the magnitude and spatial patterning of expression changes. With substantial intra-group consistency and inter-group divergence, PCA and hierarchical clustering demonstrated a clear segregation of conditions, with β-caryophyllene (BC) standing out as the most divergent chemical. UMAP visualizations provided additional evidence for this, showing that BC had the broadest log2 fold-change range and distinct geographic polarization, indicating strong and extensive proteome modulation. Moderately overlapping but distinct signals were produced by α-Myrcene (MYR) and α-Humulene (HUM), suggesting partially shared routes of differing strengths. Limonene (LIMO), on the other hand, had no effect and its expression patterns were quite similar to those of the control group. All of these findings support the idea that exposure to terpenes causes condition-specific molecular remodeling, with BC being the most bioactive of the substances examined.

**Figure 7:**
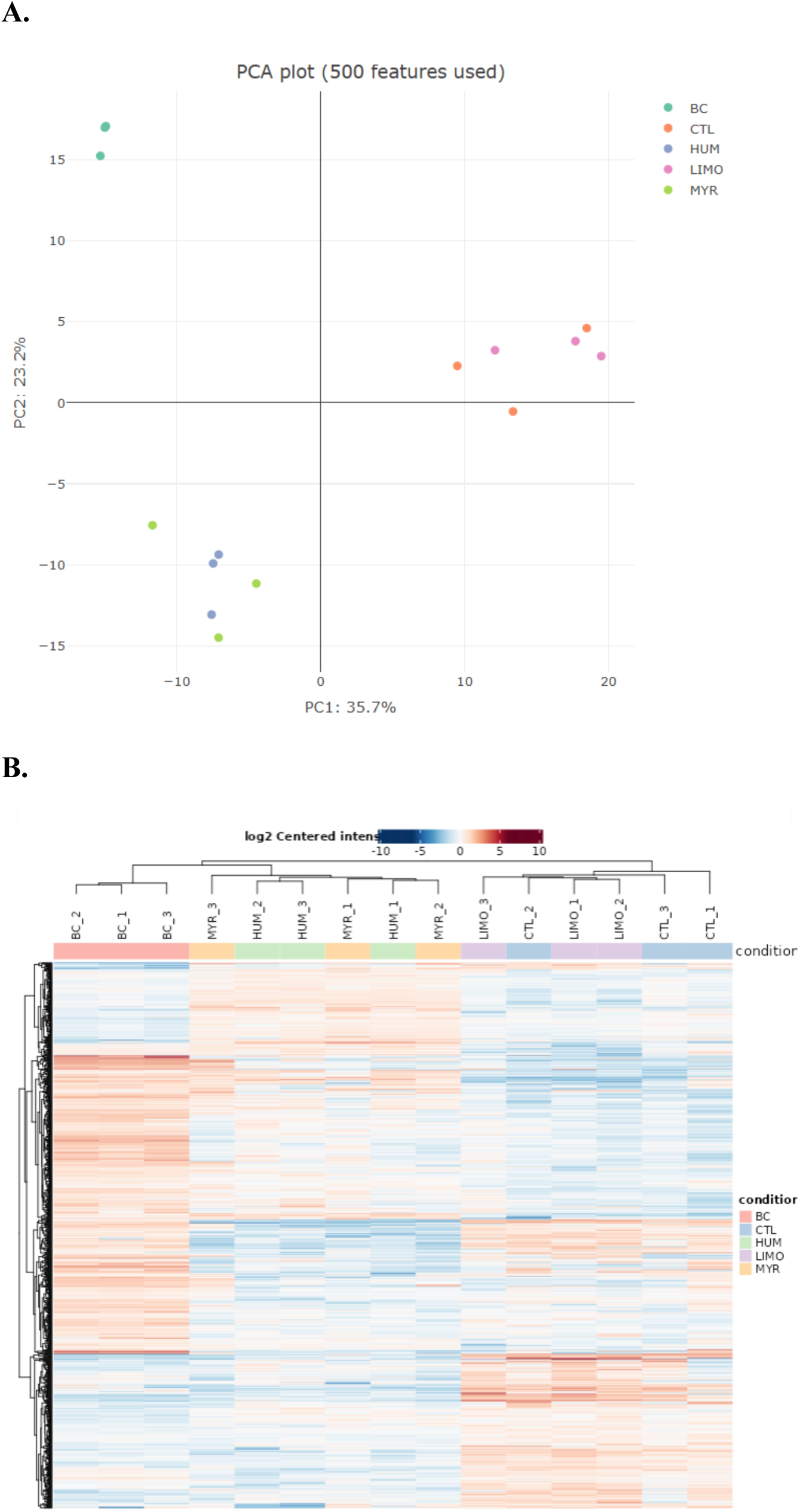
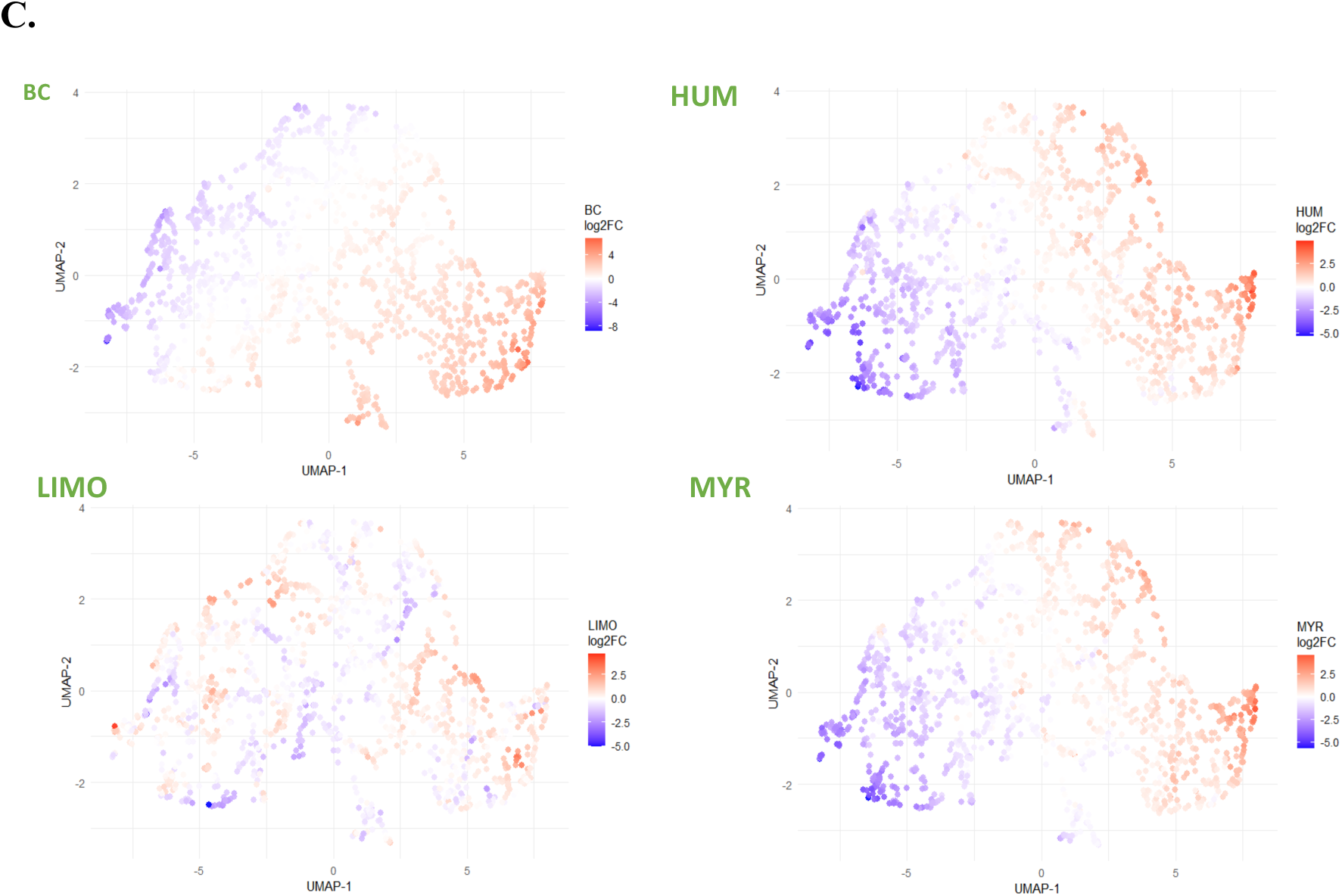
Terpene-specific proteomic signatures revealed by dimensionality reduction and clustering analyses. (A) A PCA map of the top 500 features reveals obvious sample grouping by terpene treatment, with MYR/HUM forming a distinct cluster from CTL/LIMO and β-caryophyllene (BC) clearly separated from other groups, demonstrating different and reproducible transcriptome profiles. (B) A heatmap of log2-centered expression intensities shows that BC has the most pronounced expression pattern, confirming consistent sample grouping by treatment. (C) The spatial and compound-specific translational effects of each terpene are illustrated by UMAP plots of log2 fold changes, where BC exhibits the highest and most extensive modulation and LIMO produces the least amount of change.

The PCA plot in Figure 7A supports these observations, showing that β-Caryophyllene (BC)– treated samples display the greatest global proteome divergence, clustering furthest from controls along the first principal component (PC1: 35.7%). Limonene (LIMO) exhibits minimal effects, overlapping largely with control samples, whereas α-Humulene (HUM) and α-Myrcene (MYR) form close but distinct clusters [62]. These relationships are further supported by the hierarchical clustering heatmap in Figure 7B, which reveals partial overlap between LIMO, MYR, and control groups, along with a consistent separation of BC-treated samples. Consistent with these findings, log₂ fold-change (log₂FC)–based UMAP projections in Figure 7C show distinct spatial patterns of protein modulation for each compound [63]. BC elicits the most pronounced response, marked by a left-right gradient of strong up- and down-regulation across cellular clusters. HUM and MYR display similar but less intense topologies, suggesting moderate regulatory effects, while LIMO induces only minimal transcriptional shifts, closely mirroring control distributions. Together, these clustering and dimensionality-reduction analyses demonstrate that terpene exposure triggers compound-specific translational responses, with BC driving the most extensive molecular reprogramming under heat stress.

The enrichment analyses presented in Figure 8 reveal compound-specific molecular responses to terpene exposure, highlighting distinct adaptive mechanisms triggered under heat stress. Both Gene Ontology (GO) and Reactome pathway analyses point to a coordinated modulation of metabolic, transcriptional, and neuronal processes, with differences in magnitude and directionality among β-Caryophyllene (BC), α-Humulene (HUM), α-Myrcene (MYR), and Limonene (LIMO). GO Biological Process enrichment (Figure 8A and B) identifies a strong activation of pathways associated with protein synthesis, RNA metabolism, and stress adaptation. Processes such as translation, ribonucleoprotein complex assembly, and vesicle-mediated transport are significantly upregulated in BC-treated worms, suggesting a robust compensatory translational response to thermal stress [64]. The concurrent suppression of categories linked to proteolysis and catabolic regulation supports a shift toward protein maintenance and repair rather than degradation [65]. HUM and MYR exhibit moderate enrichment across similar categories, indicating partial activation of stress-related proteostasis networks, whereas LIMO displays minimal changes, reflecting a weaker global response consistent with its limited transcriptional divergence observed in proteomic clustering analyses [65, 66]. Reactome pathway enrichment (Figure 8C and D) further supports these findings by uncovering terpene-dependent modulation of specific molecular systems. HUM and LIMO show significant enrichment in pathways related to neurotransmitter signaling, synaptic vesicle cycling, and mitochondrial energy metabolism. This suggests that these monoterpenes may influence neurophysiological activity and metabolic homeostasis, potentially explaining the subtle behavioral alterations previously observed following terpene exposure [38, 67]. BC, on the other hand, demonstrates broad and intense enrichment in pathways associated with translation initiation, ribosomal assembly, and stress granule formation, indicative of a more extensive reprogramming of cellular translation under stress conditions. Such responses are often associated with enhanced cytoprotective capacity and resilience to protein misfolding, aligning with BC’s role as a multifunctional sesquiterpene known to modulate stress and inflammatory pathways [58, 59, 60].

**Figure 8:**
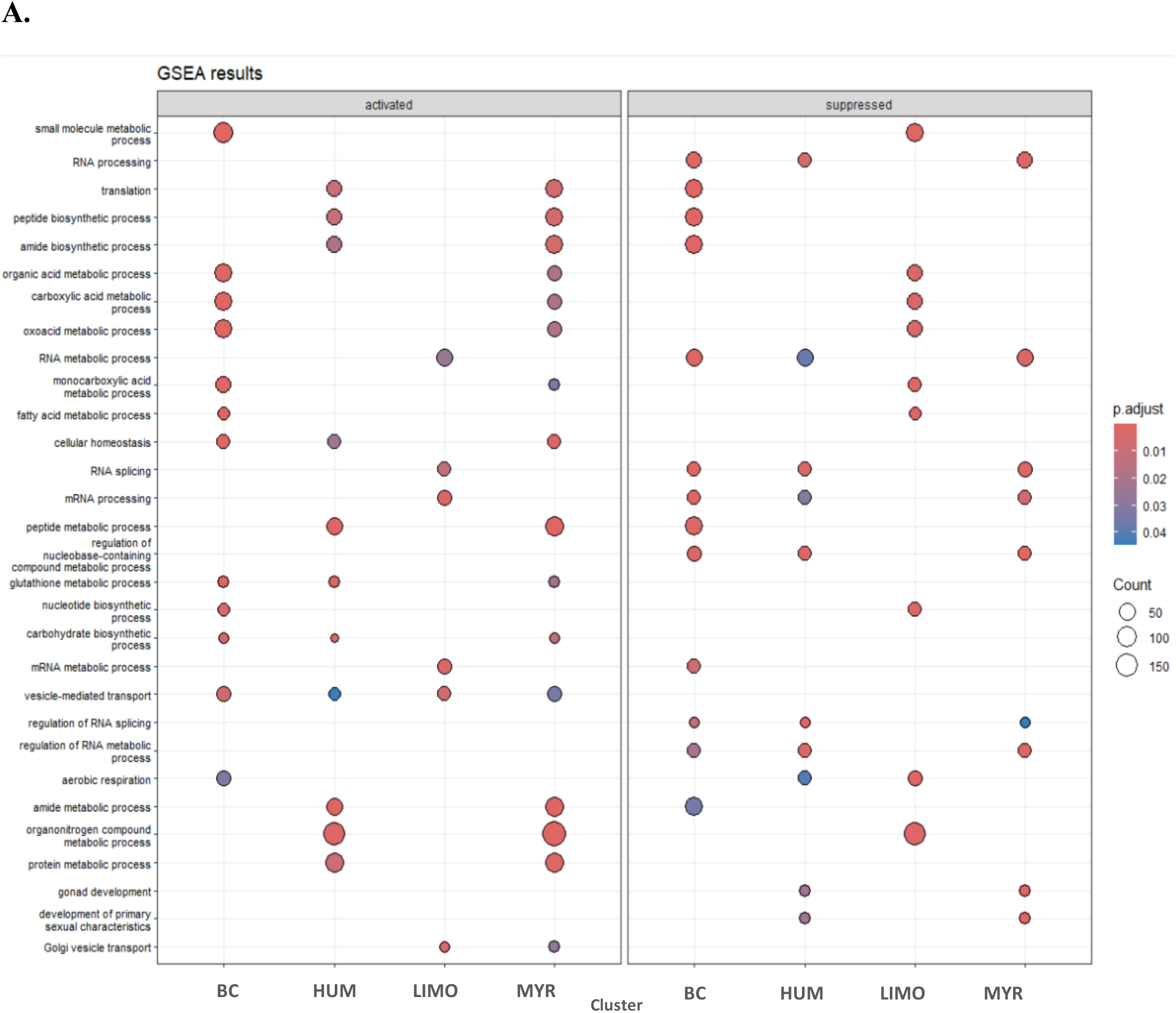
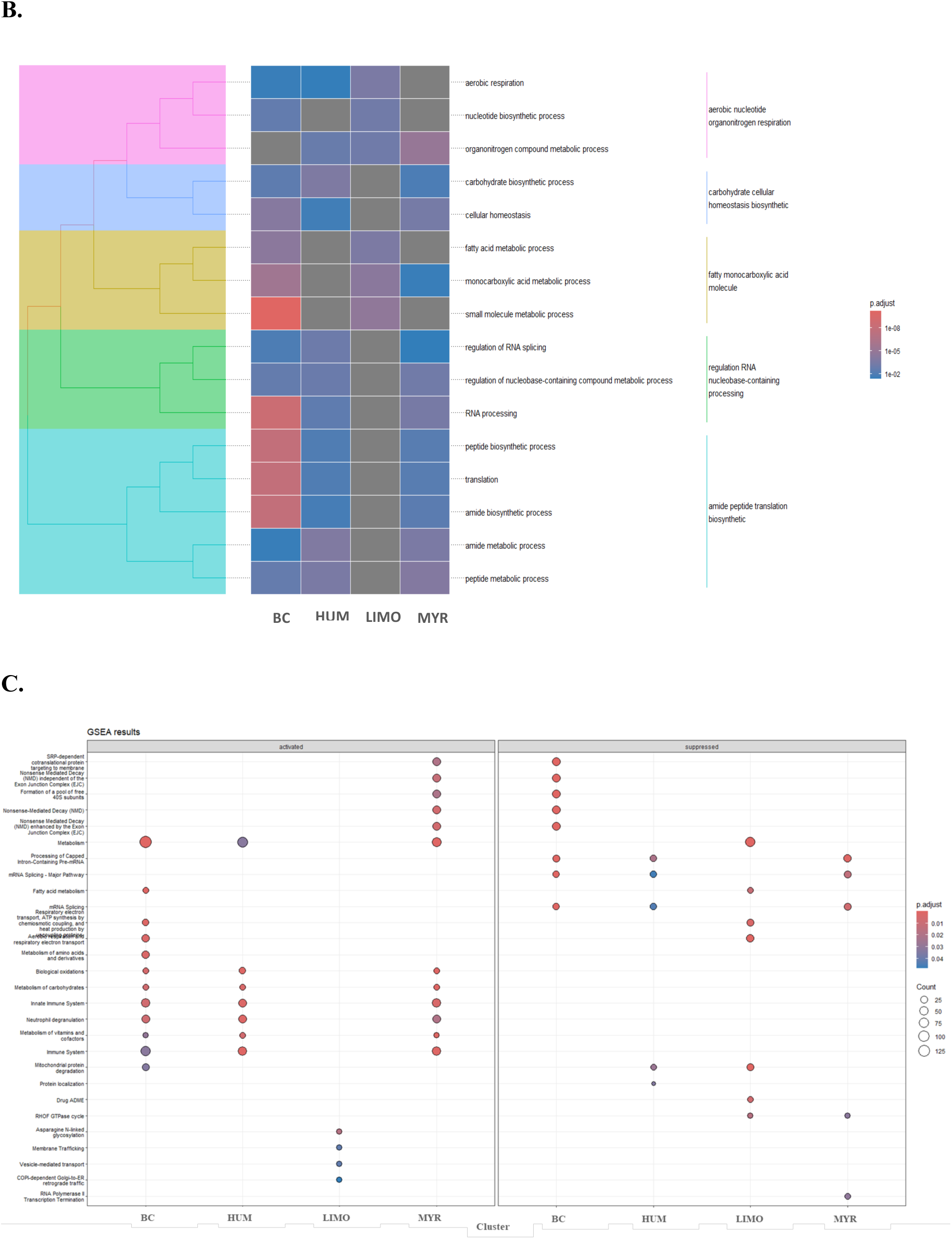
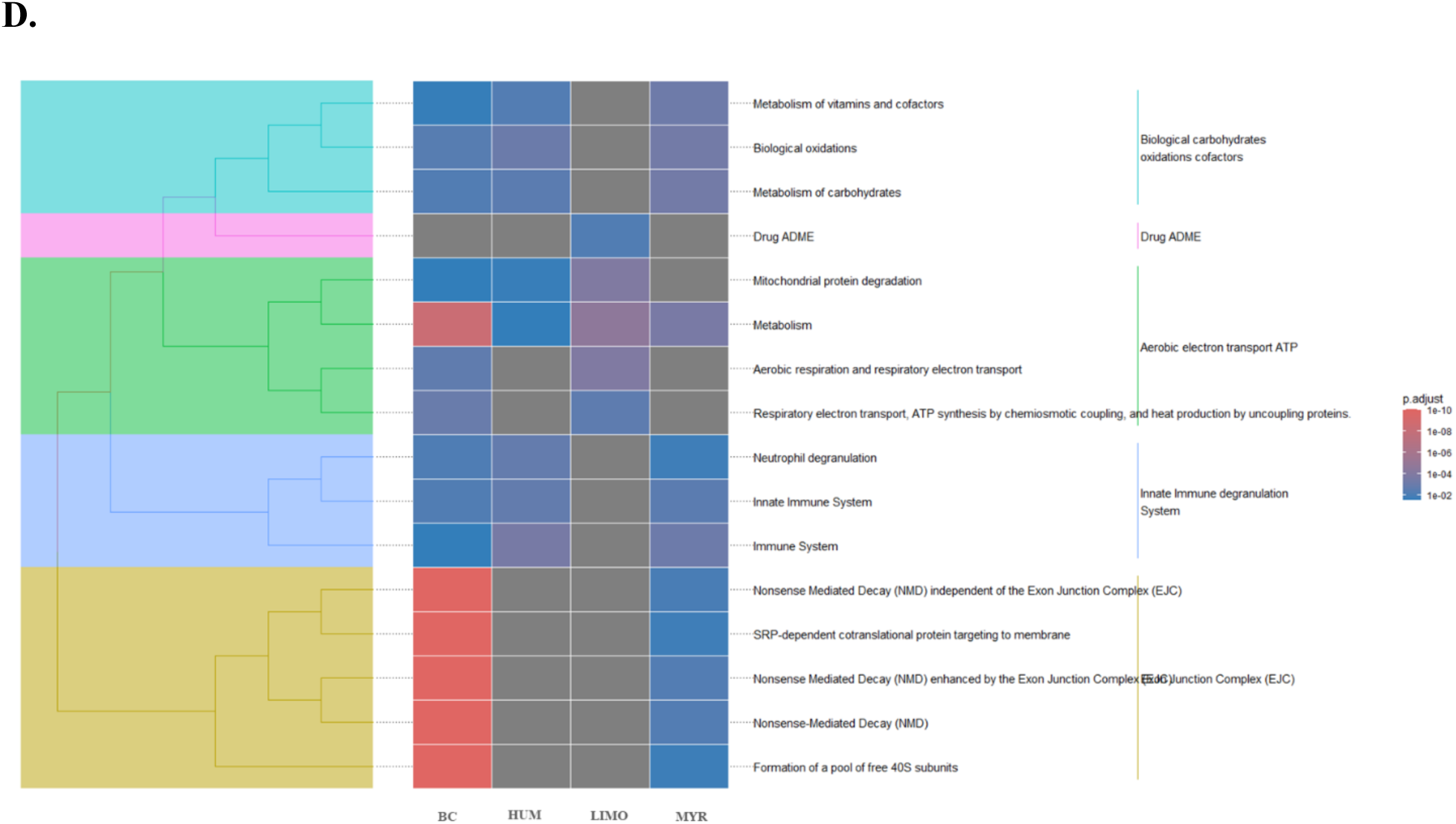
Dot plots and tree plots illustrating enrichment analysis using (A) GO terms (BP) and (B) Reactome pathways. The y-axis shows the different gene categories related to each enrichment analysis for each terpene exposed nematode group. The x-axis displays the relative abundance of protein in each term. Panel 1 displays enriched GO Biological Process terms, including categories related to metabolism, RNA processing, translation, cellular homeostasis, and neurotransmitter transport. Panel 3 shows Reactome pathway enrichments, highlighting terpene-dependent modulation of pathways involved in neuronal signaling, RNA metabolism, mitochondrial function, and innate immune responses. Notably, HUM and LIMO exhibited the strongest enrichment patterns, indicating distinct transcriptional responses under heat-induced stress.

When comparing GO and Reactome results across terpenes, a consistent theme emerges: terpenes appear to differentially shape the cellular response to heat stress by fine-tuning translational, metabolic, and neuronal signaling networks. The relative intensity of enrichment, strongest in HUM and BC, moderate in MYR, and minimal in LIMO, suggests that structural and physicochemical properties of terpenes likely influence their ability to interact with lipid membranes and cellular receptors. As lipophilic molecules, terpenes may partition into the cuticular and cellular lipid phases, altering membrane dynamics and indirectly affecting intracellular signaling cascades. Interestingly, HUM and LIMO display notable enrichment in pathways linked to neurotransmitter release and synaptic function, supporting the hypothesis that certain terpenes modulate neuronal activity via TRPV-like channels such as OSM-9 and OCR-2 [38, 64, 65]. This aligns with behavioral data showing modified nociceptive and chemotactic responses following terpene exposure. In contrast, BC’s extensive activation of translational and metabolic processes points toward a broader systemic response rather than a neuron-specific effect. Taken together, these enrichment analyses underscore the complexity of terpene action in *C. elegans*. The data suggest that distinct terpenes elicit specialized transcriptional and translational programs that balance metabolic adaptation, neuronal modulation, and proteome maintenance under stress. The finding that HUM and LIMO generate the strongest enrichment of neuronal and mitochondrial pathways, despite limited overall proteomic shifts, indicates that even subtle changes in gene regulation can lead to functionally meaningful adaptations in neuronal physiology.

These findings provide mechanistic insights relevant to pain treatment, as the terpene-induced modulation of neuronal signaling, translation, and stress-response pathways in *C. elegans* mirrors conserved processes involved in nociception across species [67, 68, 69]. The enrichment of neurotransmitter and TRP channel associated pathways suggests that terpenes such as BC, HUM, and LIMO may influence sensory neuron excitability and synaptic transmission. By fine-tuning cellular stress resilience and metabolic homeostasis, these compounds could reduce neuronal hyperactivity associated with pain sensitization in mammals. Understanding their molecular targets and translational effects thus offers promising avenues for developing non-opioid analgesics derived from natural terpenes.

## 4. Conclusion

*C. elegans* displayed clear antinociceptive effects following exposure to Limonene, α-Myrcene, α-Humulene, and β-Caryophyllene, reflected by a reduced nocifensive response to noxious heat. While α-Humulene and β-Caryophyllene may also engage the cannabinoid pathway, all four terpenes likely act, at least in part, through modulation of the vanilloid system, supporting their potential role as sensory circuit modulators. Enrichment Analysis further revealed that terpene exposure under heat stress alters key pathways involved in protein synthesis, energy metabolism, and neuronal signaling. Together, these findings provide molecular evidence that terpenes influence nociceptive-like responses by modulating neural activity, mitochondrial function, and cellular homeostasis, highlighting their promise as natural scaffolds for pain-modulating therapeutics.

## Acknowledgement

K. Benkhraba received partial tuition support from the Université de Montréal.

## Data availability

The data that support the findings from this study are provided in the supplementary file or available from the corresponding author upon reasonable request.

## Author contribution statement

KB, JS, JDC and FB conceived and designed research. KB and JDC B conducted experiments and analyzed data. KB, JS, JDC and FB wrote the manuscript. All Authors read, reviewed and approved the manuscript.

