## Supplementary material for "Terpenes as Modulators of Nociceptive Signaling: Behavioral and Molecular Insights from *Caenorhabditis elegans*": Supplental data

#### Supplementary Data

**Figure S1.** A schematic of the quadrants assay adapted from Margie *et al*. (2013). For head avoidance assay, plates were divided into quadrants two test (A and D) and two controls (B and C). Sodium azide was added to all four quadrants to paralyze nematodes. *C. elegans* were added at the center of the plate (typically, n = 100 to 1,000) and after 30 minutes, animals were counted on each quadrant. Only animals outside the inner circle were scored. The calculation of thermal avoidance index was performed has described.

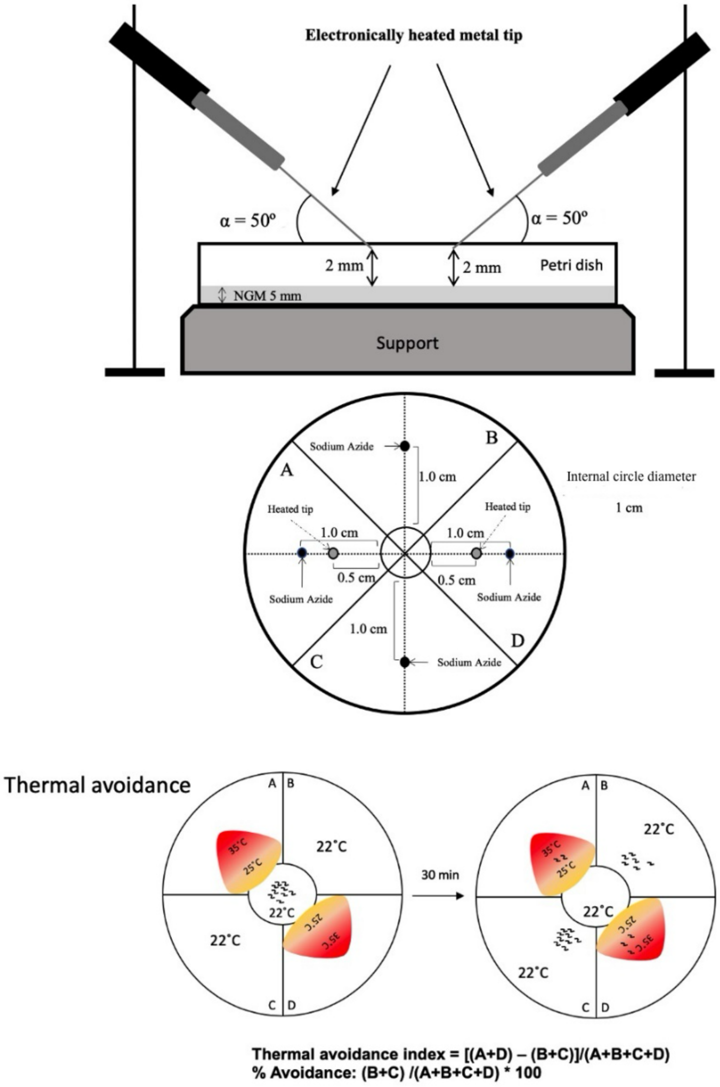

### Table S1: Protein detected and quantified for each experimental group

| Protein ID | BC_vs_CT L_log2 fold change | HUM_vs_CT L_log2 fold change | LIMO_vs_CT L_log2 fold change | MYR_vs_CT L_log2 fold change | BC_vs_CT L_p.val | HUM_vs_CT L_p.val | LIMO_vs_CT L_p.val | MYR_vs_CT L_p.val |
|---|---|---|---|---|---|---|---|---|
| Q86B36 | 6.77 | 0.441 | 3.26 | 2.61 | 9.83E-05 | 0.73 | 0.0211 | 0.0565 |
| Q21732 | 6.74 | 3.09 | 0.724 | 0.734 | 3.97E-06 | 0.00461 | 0.443 | 0.437 |
| Q23624 | 6.08 | 1.93 | 0.0696 | 1.45 | 2.43E-05 | 0.0684 | 0.944 | 0.161 |
| Q9GYI1 | 5.52 | 3.52 | 1.16 | 3.37 | 4.91E-05 | 0.00245 | 0.242 | 0.00335 |
| G5EF26 | 5.51 | 0.626 | 0.549 | 0.787 | 4.07E-07 | 0.326 | 0.387 | 0.221 |
| Q9U241 | 5.27 | 3.69 | 0.831 | 3.73 | 1.02E-07 | 6.55E-06 | 0.135 | 5.71E-06 |
| Q93725 | 5.14 | 2.09 | 1.05 | 0.475 | 5.42E-06 | 0.0117 | 0.168 | 0.52 |
| O61235 | 4.96 | 1.39 | 1.65 | 1.99 | 0.000528 | 0.228 | 0.158 | 0.0928 |
| Q20164 | 4.89 | -0.663 | -1.44 | -1.11 | 4.87E-08 | 0.17 | 0.00727 | 0.0298 |
| Q8T7Z3 | 4.87 | 3.63 | -0.206 | 2.89 | 3.52E-06 | 7.58E-05 | 0.757 | 0.000604 |
| Q8WSW2 | 4.71 | 1.01 | 2.9 | 0.639 | 1.54E-05 | 0.185 | 0.00137 | 0.394 |
| O45525 | 4.61 | 0.979 | 0.661 | 1.79 | 1.78E-05 | 0.196 | 0.374 | 0.0266 |
| O17641 | 4.49 | 0.483 | 2.13 | 0.221 | 0.000113 | 0.576 | 0.0247 | 0.797 |
| Q10453 | 4.47 | 0.565 | -0.268 | 0.382 | 1.53E-05 | 0.426 | 0.702 | 0.588 |
| Q9N4K0 | 4.44 | 0.576 | 0.0277 | 1.03 | 0.000134 | 0.509 | 0.974 | 0.245 |
| Q9NA74 | 4.36 | 1.18 | 0.551 | 1.18 | 1.17E-05 | 0.0932 | 0.414 | 0.0937 |
| G5EEE5 | 4.3 | 3.5 | 0.568 | 3.72 | 1.10E-08 | 1.42E-07 | 0.134 | 6.61E-08 |
| Q20676 | 4.28 | 2.15 | 0.76 | 2.45 | 1.42E-05 | 0.00557 | 0.265 | 0.00225 |
| Q9U228 | 4.27 | 1.39 | 0.021 | 1.38 | 6.61E-07 | 0.0141 | 0.967 | 0.0148 |
| Q18803 | 4.23 | 1.02 | -0.898 | 0.891 | 0.00232 | 0.386 | 0.443 | 0.446 |
| Q18787 | 4.21 | 0.19 | -0.0577 | 1.19 | 3.04E-05 | 0.787 | 0.935 | 0.107 |
| P34308 | 4.19 | 3.16 | 1.27 | 2.61 | 2.14E-05 | 0.00033 | 0.0785 | 0.00157 |
| G5ECV9 | 4.18 | 0.923 | 3.03 | 1.96 | 0.00281 | 0.436 | 0.0198 | 0.112 |
| Q17571 | 4.15 | 1.41 | 0.442 | 1.8 | 2.97E-06 | 0.0227 | 0.435 | 0.00566 |
| Q9XWJ5 | 4.12 | 0.412 | 0.152 | -0.091 | 6.38E-07 | 0.403 | 0.754 | 0.852 |
| Q11117 | 4.08 | 1.27 | 0.222 | 3.05 | 0.000225 | 0.147 | 0.792 | 0.00248 |
| O45496 | 3.98 | 1.19 | 0.349 | 0.878 | 3.96E-06 | 0.0448 | 0.529 | 0.127 |
| O61792 | 3.92 | 0.679 | 0.77 | 0.504 | 1.52E-05 | 0.279 | 0.223 | 0.417 |
| Q94307 | 3.9 | 0.529 | -0.0141 | 0.397 | 0.000103 | 0.478 | 0.985 | 0.593 |
| P17511 | 3.82 | -2.11 | -0.189 | -2.71 | 0.00399 | 0.0779 | 0.867 | 0.0285 |
| B7CED8 | 3.81 | 1.02 | 0.766 | 1.6 | 0.000122 | 0.179 | 0.306 | 0.044 |
| Q21154 | 3.78 | 0.743 | 0.647 | 1.19 | 5.20E-06 | 0.18 | 0.24 | 0.0399 |
| G5EF05 | 3.77 | 1.82 | 0.752 | 2.51 | 0.000266 | 0.0347 | 0.35 | 0.00622 |
| Q21966 | 3.76 | 1.25 | 0.716 | 1.83 | 1.36E-06 | 0.0178 | 0.146 | 0.00155 |
| P34461 | 3.74 | 1.29 | 0.336 | 1.2 | 1.31E-05 | 0.0395 | 0.562 | 0.0536 |
| P54812 | 3.73 | 1.13 | 3.26 | 0.379 | 4.42E-05 | 0.0977 | 0.000167 | 0.561 |
| Q18853 | 3.73 | 0.979 | -1.36 | 1.36 | 0.000693 | 0.273 | 0.136 | 0.137 |
| Q93545 | 3.72 | 0.322 | 0.603 | -0.252 | 5.56E-06 | 0.548 | 0.268 | 0.637 |
| Q95YD5 | 3.72 | 1.27 | 0.815 | 2.6 | 3.15E-05 | 0.0578 | 0.206 | 0.000868 |
| G5EBJ9 | 3.71 | 0.941 | 0.413 | 2.06 | 9.50E-05 | 0.191 | 0.557 | 0.00943 |
| G5EDZ5 | 3.58 | 1.88 | 0.354 | 3.64 | 0.000188 | 0.019 | 0.625 | 0.000162 |

| | | | | | | | | |
|---|---|---|---|---|---|---|---|---|
| Q7JLJ0 | 3.57 | 1.12 | 0.465 | 2.3 | 0.0127 | 0.384 | 0.715 | 0.0867 |
| Q9U2M4 | 3.55 | 0.843 | 0.759 | 0.506 | 7.94E-05 | 0.21 | 0.257 | 0.444 |
| H2L0K8 | 3.53 | 1.23 | 1.02 | 0.713 | 6.07E-06 | 0.0278 | 0.0595 | 0.175 |
| P91844 | 3.52 | 0.349 | 0.139 | 0.493 | 2.30E-06 | 0.457 | 0.764 | 0.298 |
| H2KYJ5 | 3.5 | 1.22 | -0.231 | 0.582 | 1.17E-05 | 0.0365 | 0.667 | 0.286 |
| H2L0Q5 | 3.5 | 0.672 | 3.38 | 0.742 | 1.17E-05 | 0.223 | 1.74E-05 | 0.181 |
| G5ECZ2 | 3.49 | 0.905 | -0.248 | 0.333 | 8.94E-06 | 0.0983 | 0.635 | 0.526 |
| P39055 | 3.48 | 2.13 | 0.722 | 1.69 | 0.00023 | 0.0093 | 0.324 | 0.0313 |
| O17606 | 3.47 | 0.761 | 0.694 | -0.118 | 1.61E-06 | 0.103 | 0.134 | 0.79 |
| O44566 | 3.47 | -0.316 | -0.87 | -0.852 | 0.000313 | 0.67 | 0.252 | 0.261 |
| O44555 | 3.46 | 0.787 | 0.721 | 0.151 | 1.71E-05 | 0.166 | 0.201 | 0.783 |
| P90780 | 3.45 | 0.793 | 0.173 | 1.43 | 1.46E-05 | 0.156 | 0.748 | 0.0172 |
| Q18655 | 3.45 | 0.338 | -0.455 | 0.336 | 8.81E-06 | 0.514 | 0.382 | 0.516 |
| Q8MXS8 | 3.45 | 0.911 | 0.694 | 0.746 | 2.15E-05 | 0.12 | 0.227 | 0.196 |
| Q9XWU2 | 3.44 | 2.15 | 0.184 | 2.34 | 0.00159 | 0.0281 | 0.837 | 0.0187 |
| P40614 | 3.42 | 1.53 | -1.79 | 1.57 | 6.95E-05 | 0.0255 | 0.0109 | 0.0219 |
| Q27487 | 3.41 | -2.17 | -0.509 | -1.75 | 0.0287 | 0.143 | 0.721 | 0.231 |
| D7SFI3 | 3.41 | 1.57 | 1.48 | 1.91 | 6.68E-07 | 0.00148 | 0.00231 | 0.000287 |
| Q20264 | 3.41 | 1.61 | -2.21 | 0.323 | 0.000677 | 0.0592 | 0.0136 | 0.686 |
| G5EGK8 | 3.39 | 3.61 | 3.05 | 3.64 | 9.48E-06 | 4.71E-06 | 2.87E-05 | 4.29E-06 |
| O45934 | 3.39 | 2.25 | 0.924 | 2.09 | 0.000176 | 0.00457 | 0.187 | 0.0073 |
| Q09364 | 3.38 | 1.96 | -0.0127 | 1.28 | 0.000885 | 0.0284 | 0.988 | 0.133 |
| O62244 | 3.38 | 1.23 | 1.05 | 0.95 | 5.76E-06 | 0.0219 | 0.0447 | 0.0656 |
| A0A0K3AVF2 | 3.37 | 3.8 | 0.285 | 4.56 | 1.73E-06 | 4.19E-07 | 0.513 | 4.56E-08 |
| P90994 | 3.36 | 1.11 | 0.706 | 2.11 | 9.80E-06 | 0.0423 | 0.177 | 0.000827 |
| G5EDK7 | 3.36 | 1.38 | 0.827 | 0.819 | 0.000406 | 0.0778 | 0.274 | 0.279 |
| Q9TZ51 | 3.35 | -0.629 | -1.47 | -0.793 | 0.000235 | 0.371 | 0.0484 | 0.264 |
| Q20584 | 3.33 | -0.097 | -0.719 | -0.715 | 6.96E-05 | 0.873 | 0.246 | 0.249 |
| L8E6W9 | 3.29 | 0.97 | 0.942 | 0.359 | 6.63E-06 | 0.0581 | 0.0647 | 0.457 |
| Q09517 | 3.27 | 1.23 | 0.858 | 0.65 | 4.11E-05 | 0.0436 | 0.144 | 0.261 |
| Q09545 | 3.26 | 4.97 | -0.885 | 3.4 | 2.85E-05 | 2.48E-07 | 0.119 | 1.83E-05 |
| O44906 | 3.26 | 2.09 | 0.628 | 1.94 | 0.000191 | 0.0061 | 0.348 | 0.00982 |
| O44443 | 3.25 | 1.11 | 1.15 | 1.64 | 1.67E-06 | 0.017 | 0.0144 | 0.00134 |
| Q4TT88 | 3.25 | 4.05 | 0.846 | 3.96 | 1.98E-05 | 1.76E-06 | 0.121 | 2.28E-06 |
| G5EBI0 | 3.22 | 1.26 | 2.36 | 1.69 | 0.000574 | 0.105 | 0.00591 | 0.0356 |
| O17694 | 3.22 | 0.712 | -0.706 | 0.583 | 0.00154 | 0.399 | 0.403 | 0.488 |
| Q9NAB4 | 3.2 | 2.83 | -0.693 | 3.03 | 0.000501 | 0.00135 | 0.345 | 0.000796 |
| Q9GZE9 | 3.19 | 1.11 | 1.18 | 0.919 | 1.63E-05 | 0.0423 | 0.0315 | 0.0843 |
| Q19663 | 3.18 | 1.14 | 0.59 | 0.851 | 0.00257 | 0.208 | 0.507 | 0.343 |
| H2KY72 | 3.17 | 1.36 | 1.58 | 0.534 | 0.000126 | 0.0406 | 0.0201 | 0.391 |
| O18239 | 3.17 | 1.51 | 0.137 | 2.66 | 0.000804 | 0.0619 | 0.856 | 0.00306 |
| Q20121 | 3.16 | 0.194 | 0.0741 | 0.679 | 0.000129 | 0.752 | 0.904 | 0.278 |
| Q19825 | 3.1 | 0.715 | 0.0815 | 0.832 | 1.15E-05 | 0.147 | 0.863 | 0.0956 |
| Q9U1Q2 | 3.06 | 2.29 | 0.0818 | 2.37 | 1.40E-05 | 0.000249 | 0.864 | 0.000179 |
| Q23571 | 3.04 | 1.16 | 0.0509 | 0.494 | 0.00019 | 0.076 | 0.934 | 0.426 |
| O44727 | 3.04 | -0.5 | 0.319 | 1.35 | 0.00548 | 0.597 | 0.735 | 0.167 |

| | | | | | | | | |
|---|---|---|---|---|---|---|---|---|
| Q27512 | 3.04 | 1.95 | 0.808 | 2.61 | 0.000463 | 0.0113 | 0.246 | 0.00162 |
| Q09596 | 3.03 | 1.35 | 0.548 | 0.835 | 1.67E-05 | 0.0126 | 0.263 | 0.0973 |
| Q20751 | 3.01 | 3.26 | 0.522 | 1.5 | 7.79E-06 | 3.22E-06 | 0.25 | 0.00399 |
| Q20502 | 3 | 1.2 | 1.36 | 0.551 | 9.80E-07 | 0.00506 | 0.00215 | 0.149 |
| P34528 | 2.99 | 3.56 | 1.19 | 3.78 | 0.000429 | 8.68E-05 | 0.0896 | 4.79E-05 |
| H2KYE0 | 2.98 | 1.48 | 0.181 | 1.26 | 0.000174 | 0.0248 | 0.762 | 0.0496 |
| Q21153 | 2.98 | 1.37 | 1.49 | 0.905 | 3.13E-06 | 0.00399 | 0.00224 | 0.039 |
| Q9U382 | 2.97 | 1.27 | 0.413 | 0.726 | 0.0012 | 0.105 | 0.581 | 0.337 |
| Q10941 | 2.96 | 1 | 0.651 | 0.876 | 0.000519 | 0.15 | 0.339 | 0.204 |
| Q18090 | 2.94 | 0.323 | 0.655 | 0.83 | 0.000391 | 0.618 | 0.319 | 0.211 |
| Q27257 | 2.94 | 0.479 | 0.193 | 0.179 | 2.63E-05 | 0.332 | 0.691 | 0.712 |
| Q09603 | 2.93 | 1.63 | 0.25 | 2.05 | 0.000504 | 0.0252 | 0.706 | 0.00715 |
| Q22021 | 2.92 | 1.38 | 0.751 | 0.496 | 0.000333 | 0.0416 | 0.243 | 0.434 |
| Q9Y041 | 2.9 | 2.06 | 1.06 | 1.81 | 1.99E-06 | 7.52E-05 | 0.0126 | 0.000257 |
| Q17334 | 2.88 | -0.422 | 0.301 | -0.235 | 9.47E-07 | 0.242 | 0.398 | 0.507 |
| B3WFV9 | 2.87 | 0.401 | 2.21 | 1.06 | 0.000816 | 0.561 | 0.00549 | 0.138 |
| Q21702 | 2.85 | 0.751 | 0.705 | 1.28 | 0.000203 | 0.209 | 0.237 | 0.0413 |
| Q10663 | 2.84 | -0.684 | 0.0121 | -0.908 | 0.00301 | 0.402 | 0.988 | 0.27 |
| Q27497 | 2.84 | 2.63 | 0.359 | 3.16 | 0.000164 | 0.00033 | 0.528 | 5.91E-05 |
| Q23384 | 2.83 | 0.576 | 0.27 | 0.388 | 1.44E-05 | 0.205 | 0.543 | 0.386 |
| O02082 | 2.82 | 3.44 | -0.874 | 2.83 | 0.00174 | 0.000343 | 0.25 | 0.00171 |
| O16453 | 2.82 | 1.42 | 0.971 | 1.08 | 0.00041 | 0.0355 | 0.134 | 0.0999 |
| Q9XXR5 | 2.81 | 0.666 | 0.116 | -0.00395 | 9.53E-05 | 0.219 | 0.825 | 0.994 |
| Q22505 | 2.79 | 1.86 | -0.524 | 2.01 | 0.000315 | 0.00681 | 0.386 | 0.00417 |
| G5EBV4 | 2.75 | 0.783 | -0.252 | 0.96 | 0.000462 | 0.215 | 0.682 | 0.134 |
| O02286 | 2.75 | 1.53 | -0.144 | 1.42 | 9.98E-07 | 0.000408 | 0.671 | 0.000777 |
| O16689 | 2.74 | 1.43 | 0.621 | 1.42 | 0.000215 | 0.0215 | 0.278 | 0.0224 |
| Q8WQA4 | 2.73 | -0.733 | 0.789 | -1.28 | 0.00132 | 0.299 | 0.265 | 0.081 |
| A0A5S9MMA9 | 2.73 | 0.7 | 0.571 | 0.699 | 5.44E-05 | 0.163 | 0.249 | 0.163 |
| P41994 | 2.71 | -0.0387 | 1.05 | 0.969 | 0.00687 | 0.965 | 0.241 | 0.276 |
| Q23381 | 2.7 | 2.61 | -1.64 | 3.1 | 0.0169 | 0.02 | 0.121 | 0.00767 |
| Q5FC71 | 2.7 | 0.328 | 0.187 | 0.77 | 0.00295 | 0.668 | 0.806 | 0.321 |
| P91390 | 2.69 | 1.39 | 1.07 | 1.35 | 0.00485 | 0.105 | 0.205 | 0.115 |
| Q07750 | 2.69 | 2.36 | -1.43 | 1.12 | 0.0117 | 0.0234 | 0.145 | 0.247 |
| O76367 | 2.68 | -0.202 | -0.224 | 0.614 | 0.0016 | 0.772 | 0.748 | 0.385 |
| Q9N538 | 2.68 | 0.542 | 0.678 | 0.611 | 0.00184 | 0.451 | 0.348 | 0.397 |
| Q95PZ1 | 2.66 | 1.11 | 0.18 | 1.4 | 0.000183 | 0.0528 | 0.737 | 0.0189 |
| Q94392 | 2.65 | 0.045 | -0.285 | -0.195 | 3.00E-04 | 0.936 | 0.614 | 0.73 |
| Q22633 | 2.63 | 1.85 | 1.06 | 1.28 | 0.000193 | 0.00337 | 0.0615 | 0.0279 |
| G5EFS2 | 2.62 | 0.103 | 0.53 | -0.113 | 0.000162 | 0.843 | 0.318 | 0.828 |
| G5EGT7 | 2.62 | 0.787 | 1.4 | 0.549 | 0.013 | 0.406 | 0.15 | 0.56 |
| Q9N5Y2 | 2.59 | 0.814 | -0.104 | 0.21 | 5.61E-05 | 0.0939 | 0.821 | 0.649 |
| Q03575 | 2.58 | 0.426 | -0.151 | 0.565 | 0.00523 | 0.593 | 0.849 | 0.48 |
| A0A486WV52 | 2.58 | -0.414 | -0.307 | 0.622 | 0.00207 | 0.553 | 0.659 | 0.377 |
| Q9GUF2 | 2.58 | 1.36 | 1.05 | 2.59 | 3.42E-05 | 0.0068 | 0.0286 | 3.32E-05 |

| | | | | | | | | |
|---|---|---|---|---|---|---|---|---|
| Q20223 | 2.57 | 1.45 | 0.179 | 1.5 | 4.20E-05 | 0.00512 | 0.689 | 0.00415 |
| B4DCT1 | 2.57 | 1.12 | 0.413 | 0.778 | 0.0015 | 0.11 | 0.537 | 0.253 |
| P52814 | 2.56 | 3.78 | 0.196 | 2.72 | 0.0174 | 0.00138 | 0.839 | 0.0126 |
| Q9U2J6 | 2.56 | 1.32 | 0.0359 | 1.96 | 0.00102 | 0.051 | 0.954 | 0.00684 |
| P17139 | 2.55 | 0.633 | -0.489 | -0.239 | 4.81E-05 | 0.171 | 0.284 | 0.594 |
| Q23445 | 2.55 | 2.56 | 0.462 | 2.67 | 0.00297 | 0.00282 | 0.524 | 0.00208 |
| Q9U2D9 | 2.55 | 0.203 | 0.534 | 1.83 | 0.00019 | 0.693 | 0.309 | 0.00282 |
| Q18358 | 2.54 | 1 | 0.273 | 0.215 | 0.00817 | 0.245 | 0.745 | 0.798 |
| L8E833 | 2.54 | -0.341 | -0.314 | 1.84 | 0.0485 | 0.776 | 0.793 | 0.14 |
| Q93204 | 2.53 | -0.943 | 0.11 | -0.938 | 0.00433 | 0.225 | 0.884 | 0.227 |
| A0A1N7SYN7 | 2.52 | -0.502 | -0.228 | -0.194 | 0.00129 | 0.437 | 0.721 | 0.761 |
| B5BM47 | 2.52 | 1.19 | 0.45 | 1.08 | 0.00193 | 0.0922 | 0.506 | 0.124 |
| O45865 | 2.52 | 0.713 | -0.869 | 1.03 | 0.000178 | 0.173 | 0.102 | 0.0569 |
| Q19655 | 2.52 | 0.263 | -0.188 | 0.875 | 1.45E-05 | 0.507 | 0.634 | 0.0397 |
| Q22601 | 2.52 | 1.35 | 1.03 | 0.952 | 0.000493 | 0.0292 | 0.0854 | 0.11 |
| P24894 | 2.51 | 0.376 | -0.643 | 1.06 | 0.00187 | 0.575 | 0.343 | 0.128 |
| Q04457 | 2.51 | 1.06 | 0.0169 | 0.0521 | 0.000622 | 0.0848 | 0.977 | 0.928 |
| G5EDE8 | 2.51 | 0.961 | 0.119 | 1.34 | 0.000653 | 0.116 | 0.839 | 0.0357 |
| Q9TYL2 | 2.5 | 1.32 | 0.36 | 0.463 | 0.00081 | 0.0413 | 0.549 | 0.442 |
| Q21285 | 2.48 | 1.05 | 0.202 | 1.98 | 0.00155 | 0.117 | 0.753 | 0.00726 |
| A0A2C9C352 | 2.47 | 2.03 | 1.67 | 1.47 | 0.0046 | 0.0152 | 0.0385 | 0.0639 |
| A0A8D9I6U5 | 2.47 | 1.9 | 1.92 | 2.55 | 2.87E-05 | 0.000355 | 0.000319 | 2.12E-05 |
| O45783 | 2.47 | -0.541 | 0.494 | -1.19 | 0.0164 | 0.56 | 0.594 | 0.211 |
| Q95QL1 | 2.47 | 0.0767 | 0.722 | 0.548 | 8.47E-05 | 0.867 | 0.131 | 0.244 |
| Q9GZH5 | 2.47 | 1.14 | 0.12 | 0.646 | 0.00743 | 0.172 | 0.881 | 0.427 |
| O17732 | 2.46 | 1.6 | 0.249 | 0.977 | 0.000793 | 0.0147 | 0.672 | 0.111 |
| Q18864 | 2.46 | 0.241 | -0.488 | 0.361 | 0.00376 | 0.739 | 0.502 | 0.618 |
| Q9TXY8 | 2.46 | -1.11 | -0.307 | -0.045 | 0.0392 | 0.32 | 0.78 | 0.967 |
| C9IY21 | 2.45 | 0.861 | 0.128 | 0.644 | 0.0084 | 0.299 | 0.875 | 0.433 |
| O02089 | 2.45 | 0.667 | 1.41 | -0.612 | 0.00118 | 0.287 | 0.0346 | 0.327 |
| O44509 | 2.45 | 1.33 | 0.36 | 0.62 | 0.00105 | 0.042 | 0.555 | 0.314 |
| Q22352 | 2.45 | 2.63 | 0.139 | 1.45 | 0.00876 | 0.00572 | 0.866 | 0.0929 |
| Q9N4A5 | 2.45 | 1.4 | 0.833 | 0.716 | 0.00801 | 0.0988 | 0.311 | 0.382 |
| Q9XU13 | 2.45 | 2.67 | 0.482 | 2.97 | 0.00897 | 0.00512 | 0.559 | 0.0025 |
| Q18040 | 2.44 | 1.89 | -0.106 | 1.73 | 0.00218 | 0.0115 | 0.872 | 0.0187 |
| O17271 | 2.43 | 0.486 | 0.181 | 0.668 | 0.00356 | 0.494 | 0.798 | 0.351 |
| Q18625 | 2.43 | 1.42 | 2.04 | 1.18 | 0.00275 | 0.0526 | 0.00869 | 0.1 |
| O45218 | 2.42 | 1.75 | 0.206 | 1.83 | 0.000327 | 0.00424 | 0.693 | 0.00301 |
| O45941 | 2.41 | 1.47 | -0.262 | 3.66 | 0.00319 | 0.0482 | 0.705 | 9.97E-05 |
| P30627 | 2.41 | 0.489 | -0.253 | 0.189 | 1.08E-05 | 0.196 | 0.493 | 0.607 |
| Q21067 | 2.41 | 0.41 | -0.673 | -0.15 | 5.10E-05 | 0.342 | 0.129 | 0.724 |
| G5EFS5 | 2.41 | 0.565 | 0.384 | -0.217 | 3.40E-05 | 0.181 | 0.354 | 0.598 |
| O44608 | 2.41 | 0.369 | -0.0306 | 0.555 | 7.57E-05 | 0.409 | 0.945 | 0.221 |
| O61832 | 2.41 | 0.834 | -0.286 | 0.562 | 5.57E-05 | 0.0679 | 0.508 | 0.204 |

| | | | | | | | | |
|---|---|---|---|---|---|---|---|---|
| Q21994 | 2.41 | 1.04 | 0.645 | 1.56 | 0.00178 | 0.116 | 0.319 | 0.0258 |
| G5EF86 | 2.41 | 1.14 | 1.25 | 2.07 | 0.0183 | 0.226 | 0.187 | 0.0377 |
| Q17405 | 2.4 | -0.618 | -1.06 | -0.626 | 0.0127 | 0.475 | 0.227 | 0.469 |
| A7DTF0 | 2.37 | 1.22 | 0.249 | -0.1 | 0.00482 | 0.106 | 0.73 | 0.889 |
| O62246 | 2.37 | 0.217 | 0.487 | 1.58 | 0.00442 | 0.76 | 0.497 | 0.0396 |
| Q9XWD1 | 2.37 | 0.359 | 0.0783 | -0.24 | 0.000327 | 0.484 | 0.878 | 0.639 |
| G5EEK9 | 2.36 | 0.844 | 0.82 | 1.24 | 2.39E-05 | 0.0429 | 0.0483 | 0.00549 |
| P36609 | 2.36 | 2.89 | -0.482 | 0.0571 | 0.051 | 0.0207 | 0.67 | 0.96 |
| Q09657 | 2.36 | -0.634 | -0.701 | -0.303 | 0.000127 | 0.18 | 0.141 | 0.511 |
| O17071 | 2.35 | 0.652 | 0.251 | 0.344 | 0.000244 | 0.195 | 0.609 | 0.485 |
| Q17802 | 2.35 | 2.21 | 3.08 | 3.41 | 0.00325 | 0.005 | 0.000388 | 0.000156 |
| Q94234 | 2.35 | 1.35 | 0.398 | 0.393 | 9.50E-05 | 0.00771 | 0.374 | 0.379 |
| O17245 | 2.35 | 0.449 | 0.399 | -0.203 | 0.00015 | 0.341 | 0.396 | 0.662 |
| P91372 | 2.34 | 0.272 | 0.315 | -0.226 | 0.0017 | 0.659 | 0.609 | 0.713 |
| Q21742 | 2.34 | 2.32 | 0.69 | 1.65 | 0.00288 | 0.00307 | 0.305 | 0.0236 |
| Q2L6V7 | 2.34 | 0.73 | -0.12 | 0.352 | 0.00016 | 0.132 | 0.796 | 0.453 |
| O01257 | 2.33 | 0.296 | -0.295 | 0.446 | 0.000803 | 0.596 | 0.597 | 0.428 |
| Q94055 | 2.33 | 0.605 | -0.465 | 1.26 | 0.00727 | 0.428 | 0.54 | 0.111 |
| G5EFC1 | 2.32 | 1.75 | 0.175 | 1.83 | 0.000427 | 0.00379 | 0.734 | 0.00277 |
| Q69Z13 | 2.32 | 1.56 | 0.602 | 0.985 | 0.0016 | 0.02 | 0.328 | 0.12 |
| Q06561 | 2.31 | 1.09 | 1.33 | 0.871 | 0.000268 | 0.0378 | 0.0145 | 0.0885 |
| Q23500 | 2.31 | 2.17 | 0.287 | 1.86 | 8.04E-05 | 0.000143 | 0.504 | 0.000578 |
| C3JXD8 | 2.31 | 3.79 | 1.3 | 4.18 | 0.00939 | 0.000223 | 0.113 | 8.76E-05 |
| Q23091 | 2.31 | 2 | 1.51 | 1.24 | 0.00455 | 0.0113 | 0.0443 | 0.0905 |
| O01761 | 2.31 | 1.7 | 1.97 | 0.574 | 0.0808 | 0.188 | 0.13 | 0.647 |
| G5EC23 | 2.27 | 0.822 | 0.787 | 1.46 | 0.000133 | 0.0795 | 0.0916 | 0.00467 |
| P53585 | 2.27 | 1.58 | 0.559 | 3.73 | 0.00705 | 0.0457 | 0.45 | 0.000147 |
| Q9N362 | 2.27 | 3.22 | 0.00634 | 2.38 | 0.000109 | 2.88E-06 | 0.988 | 7.00E-05 |
| Q9N4I0 | 2.26 | 1.46 | 0.0712 | 0.179 | 0.00121 | 0.0201 | 0.9 | 0.753 |
| O44549 | 2.26 | -0.182 | -0.387 | 0.864 | 0.0123 | 0.82 | 0.629 | 0.289 |
| P34517 | 2.25 | 0.31 | -0.159 | 0.704 | 0.022 | 0.728 | 0.858 | 0.433 |
| Q9XTT9 | 2.24 | 0.734 | 0.931 | 1.36 | 0.00378 | 0.274 | 0.171 | 0.0528 |
| O45011 | 2.24 | -1.6 | 1.6 | 0.0366 | 0.00347 | 0.0248 | 0.0248 | 0.955 |
| Q9TYL9 | 2.24 | 1.94 | -0.175 | 0.431 | 0.0025 | 0.00672 | 0.778 | 0.49 |
| Q23642 | 2.24 | 1.26 | 2.46 | 0.536 | 0.0446 | 0.235 | 0.0297 | 0.605 |
| Q09279 | 2.23 | 0.635 | 0.274 | -0.157 | 0.00106 | 0.259 | 0.619 | 0.775 |
| Q18938 | 2.23 | 0.563 | -0.195 | 0.575 | 0.00467 | 0.41 | 0.773 | 0.4 |
| Q09490 | 2.23 | 1.45 | -0.0937 | 2.18 | 0.0255 | 0.125 | 0.918 | 0.0286 |
| Q09629 | 2.22 | 0.421 | -0.592 | -0.391 | 0.0133 | 0.598 | 0.461 | 0.624 |
| P90860 | 2.22 | -0.236 | -0.908 | 1.18 | 0.013 | 0.766 | 0.262 | 0.152 |
| Q95XX1 | 2.2 | 0.605 | 0.531 | -0.691 | 7.11E-05 | 0.147 | 0.199 | 0.101 |
| P92020 | 2.2 | 1.17 | -0.233 | 0.376 | 0.00146 | 0.0536 | 0.681 | 0.51 |
| Q95017 | 2.2 | 1.89 | 1.03 | 1.2 | 0.0274 | 0.0527 | 0.269 | 0.201 |
| Q21850 | 2.18 | 0.804 | 0.642 | -0.154 | 0.000554 | 0.122 | 0.21 | 0.757 |
| P34496 | 2.17 | 1.67 | 0.626 | 0.598 | 0.00337 | 0.0165 | 0.325 | 0.346 |
| G5ECW7 | 2.17 | 3.41 | 0.311 | 2.71 | 0.00054 | 6.62E-06 | 0.532 | 7.12E-05 |

| | | | | | | | | |
|---|---|---|---|---|---|---|---|---|
| A0A0K3A RI2 | 2.16 | 0.514 | 0.444 | -0.211 | 0.00142 | 0.36 | 0.427 | 0.704 |
| C0HLB3 | 2.15 | 1.77 | -0.124 | 0.928 | 0.00995 | 0.028 | 0.866 | 0.219 |
| P34574 | 2.15 | 1.79 | -0.164 | 2.13 | 0.000102 | 0.000541 | 0.688 | 0.000113 |
| Q7YZW5 | 2.15 | 1.83 | 1.04 | 1.1 | 0.000248 | 0.000977 | 0.0336 | 0.0257 |
| A5Z2T8 | 2.15 | -0.714 | 0.221 | 0.272 | 0.00239 | 0.238 | 0.708 | 0.646 |
| H2KYM0 | 2.15 | -0.238 | -0.35 | -0.338 | 0.000587 | 0.632 | 0.482 | 0.498 |
| O45502 | 2.15 | 3.28 | 0.54 | 2.53 | 0.000244 | 3.30E-06 | 0.239 | 5.20E-05 |
| Q17474 | 2.15 | 1.12 | -0.246 | 1.09 | 0.00341 | 0.0867 | 0.693 | 0.095 |
| A0A131M BU3 | 2.14 | 2.7 | 1.23 | 2.85 | 0.00663 | 0.00129 | 0.0877 | 0.00083 |
| G5ECG7 | 2.14 | -0.314 | 0.415 | 0.465 | 0.00729 | 0.651 | 0.552 | 0.505 |
| H2L023 | 2.14 | 0.414 | -0.142 | -0.209 | 0.00297 | 0.498 | 0.814 | 0.73 |
| Q27GU2 | 2.14 | 0.887 | 0.126 | 1.26 | 0.011 | 0.244 | 0.865 | 0.107 |
| G5EDD4 | 2.13 | 0.773 | 0.098 | 0.359 | 0.0024 | 0.2 | 0.867 | 0.542 |
| Q19202 | 2.12 | 1.07 | 0.582 | 0.771 | 0.00192 | 0.0739 | 0.311 | 0.185 |
| Q8MXI1 | 2.12 | 1.46 | 0.598 | 0.505 | 0.0023 | 0.0222 | 0.31 | 0.389 |
| Q17624 | 2.11 | 0.894 | 0.16 | 1.02 | 0.00151 | 0.116 | 0.768 | 0.076 |
| Q9XVS1 | 2.1 | 0.879 | -0.673 | 0.466 | 0.00468 | 0.18 | 0.299 | 0.467 |
| A0A0K3A XF6 | 2.1 | 0.0971 | 1.01 | 0.461 | 0.00752 | 0.887 | 0.155 | 0.504 |
| P91491 | 2.1 | 1.06 | 0.273 | 0.846 | 0.000965 | 0.0546 | 0.596 | 0.115 |
| Q17512 | 2.1 | -0.202 | 0.0331 | 1.79 | 0.00352 | 0.74 | 0.957 | 0.00968 |
| O76449 | 2.09 | 0.0759 | -0.387 | -0.135 | 0.00197 | 0.892 | 0.491 | 0.808 |
| Q09489 | 2.09 | 1.85 | 0.12 | 1.33 | 0.000253 | 0.000738 | 0.783 | 0.00806 |
| Q19126 | 2.08 | 0.437 | -1.47 | -0.135 | 0.00147 | 0.42 | 0.0146 | 0.801 |
| Q20589 | 2.08 | 0.463 | -0.0563 | 1.08 | 0.00775 | 0.499 | 0.934 | 0.127 |
| Q9N3H3 | 2.08 | 0.387 | -0.53 | 0.461 | 0.019 | 0.629 | 0.51 | 0.566 |
| Q9U329 | 2.08 | 0.349 | -0.603 | 1.84 | 0.0131 | 0.64 | 0.422 | 0.0248 |
| O16202 | 2.07 | 1.59 | 0.244 | 2.18 | 0.000353 | 0.00283 | 0.587 | 0.000219 |
| O62183 | 2.06 | 0.868 | 0.197 | 0.5 | 0.0041 | 0.169 | 0.747 | 0.418 |
| G5EBR1 | 2.05 | 2.34 | 0.224 | 1.91 | 5.95E-05 | 1.51E-05 | 0.545 | 0.00012 |
| Q20780 | 2.05 | -0.425 | -0.586 | -0.432 | 1.62E-05 | 0.201 | 0.0859 | 0.194 |
| Q9XVR3 | 2.05 | 0.852 | 0.253 | 0.74 | 0.00462 | 0.183 | 0.683 | 0.243 |
| G5EEH6 | 2.05 | 0.793 | 0.688 | 1.22 | 0.0237 | 0.343 | 0.409 | 0.153 |
| A0A2C9C 320 | 2.03 | 2.21 | -0.26 | 1.05 | 0.00207 | 0.00107 | 0.635 | 0.0707 |
| E0AHA7 | 2.03 | 1.3 | -0.221 | 1.27 | 0.00135 | 0.0227 | 0.67 | 0.025 |
| Q21322 | 2.03 | 1.56 | 0.555 | 0.946 | 0.00882 | 0.0352 | 0.419 | 0.178 |
| Q8MYN1 | 2.03 | 2.06 | -0.181 | 0.806 | 0.0102 | 0.00938 | 0.794 | 0.258 |
| Q9XUU9 | 2.02 | 1.37 | 0.183 | 0.133 | 0.000885 | 0.0128 | 0.708 | 0.785 |
| G3MU66 | 2.02 | -0.284 | -0.654 | 0.0992 | 0.00393 | 0.635 | 0.283 | 0.868 |
| O16521 | 2.02 | 0.573 | -0.124 | 0.269 | 0.00347 | 0.334 | 0.831 | 0.646 |
| O45106 | 2.02 | 0.414 | 0.0282 | -0.266 | 0.00212 | 0.451 | 0.959 | 0.626 |
| Q19722 | 2.01 | 3.36 | 0.183 | 3 | 0.000548 | 3.38E-06 | 0.691 | 1.18E-05 |
| O17643 | 2.01 | 0.214 | -0.124 | 0.926 | 0.00309 | 0.708 | 0.828 | 0.121 |
| Q9N3W5 | 2 | -0.031 | -0.521 | -0.558 | 0.00501 | 0.96 | 0.399 | 0.367 |

| | | | | | | | | |
|---|---|---|---|---|---|---|---|---|
| P34492 | 1.99 | 1.08 | 0.469 | 0.406 | 0.00129 | 0.0458 | 0.359 | 0.425 |
| Q10667 | 1.98 | 1.3 | 0.519 | 1.25 | 0.00882 | 0.065 | 0.438 | 0.075 |
| O17939 | 1.98 | 3.23 | 0.387 | 2.01 | 0.08 | 0.00813 | 0.717 | 0.075 |
| Q86MI3 | 1.98 | 0.31 | 0.249 | 0.587 | 0.00395 | 0.596 | 0.67 | 0.322 |
| G5EGH7 | 1.97 | 2.97 | 0.531 | 1.91 | 0.00683 | 0.000291 | 0.405 | 0.00811 |
| Q20140 | 1.97 | 0.865 | 0.322 | 1.26 | 0.000409 | 0.0616 | 0.462 | 0.0104 |
| Q21525 | 1.97 | -0.242 | 0.394 | -0.25 | 0.00439 | 0.683 | 0.509 | 0.673 |
| Q8TA47 | 1.97 | 0.241 | -0.461 | 2.01 | 0.0052 | 0.691 | 0.451 | 0.00455 |
| Q09607 | 1.96 | 1.82 | -2.05 | 1.18 | 0.00926 | 0.0143 | 0.007 | 0.0901 |
| A0A3B1DVR7 | 1.96 | 1.07 | 0.374 | 0.917 | 0.00128 | 0.0457 | 0.454 | 0.0802 |
| G5EDW8 | 1.96 | -0.396 | -0.796 | 1.17 | 0.0132 | 0.575 | 0.268 | 0.112 |
| Q22067 | 1.95 | 0.782 | -0.329 | 0.962 | 5.23E-05 | 0.0369 | 0.347 | 0.0132 |
| O45060 | 1.95 | -1.5 | -0.397 | -0.839 | 0.0135 | 0.0473 | 0.574 | 0.244 |
| Q8MXH3 | 1.95 | -0.0544 | -0.124 | -0.409 | 0.0224 | 0.944 | 0.872 | 0.598 |
| Q95XJ0 | 1.95 | 0.828 | 0.684 | 0.525 | 0.0101 | 0.227 | 0.314 | 0.436 |
| Q23089 | 1.94 | 1.41 | 0.56 | 0.283 | 0.000307 | 0.00374 | 0.19 | 0.496 |
| Q03206 | 1.93 | 0.227 | -0.162 | 0.505 | 0.00314 | 0.681 | 0.769 | 0.367 |
| A0A486WWU7 | 1.93 | 0.546 | 0.252 | -0.396 | 0.0031 | 0.33 | 0.649 | 0.476 |
| G5EE41 | 1.93 | -1.18 | 0.183 | -2 | 0.00469 | 0.0579 | 0.754 | 0.00363 |
| Q9U238 | 1.93 | 1.05 | 0.973 | 2.68 | 0.0175 | 0.165 | 0.196 | 0.00224 |
| Q18115 | 1.92 | 1.6 | 2.5 | 1.06 | 0.00591 | 0.0174 | 0.000873 | 0.0947 |
| P91180 | 1.92 | -1.44 | -1.08 | -1.85 | 0.0555 | 0.139 | 0.261 | 0.0646 |
| Q9N3Y1 | 1.92 | 2.66 | 0.766 | 2.3 | 0.00233 | 0.000151 | 0.159 | 0.000551 |
| Q9U2Q9 | 1.91 | 2.82 | 0.0799 | 2.17 | 0.00842 | 0.000477 | 0.9 | 0.00363 |
| Q9XU97 | 1.91 | -0.0919 | -0.26 | -0.37 | 0.00976 | 0.887 | 0.689 | 0.57 |
| Q21018 | 1.91 | 0.435 | 0.275 | 1.36 | 0.0194 | 0.556 | 0.709 | 0.0813 |
| G5ECR0 | 1.9 | 2 | -0.561 | 2.28 | 0.00569 | 0.00413 | 0.351 | 0.00157 |
| G5EGU1 | 1.9 | 0.926 | 0.0979 | 0.327 | 0.00293 | 0.101 | 0.855 | 0.545 |
| Q21481 | 1.9 | 1.47 | 0.144 | 1.03 | 0.00285 | 0.0145 | 0.788 | 0.07 |
| P34575 | 1.89 | -0.201 | -0.997 | 0.406 | 9.45E-05 | 0.574 | 0.0128 | 0.264 |
| Q95YB2 | 1.88 | 0.875 | -0.103 | 1.34 | 0.00964 | 0.184 | 0.872 | 0.05 |
| P46561 | 1.87 | 0.465 | -0.525 | 0.47 | 0.000237 | 0.24 | 0.188 | 0.236 |
| Q9XUT0 | 1.87 | 1.66 | 0.231 | 0.873 | 0.0086 | 0.0168 | 0.711 | 0.175 |
| Q93235 | 1.86 | 0.861 | -0.435 | 1.47 | 0.00643 | 0.16 | 0.466 | 0.0238 |
| O76672 | 1.86 | -2.19 | -0.974 | 0.0221 | 0.141 | 0.0865 | 0.427 | 0.985 |
| G5ECU1 | 1.85 | -0.277 | -0.399 | -0.117 | 0.00421 | 0.616 | 0.472 | 0.831 |
| Q10013 | 1.85 | 0.413 | -0.371 | 1.74 | 0.0121 | 0.53 | 0.572 | 0.0173 |
| A0A2C9C2E8 | 1.85 | 0.711 | 2.87 | 0.847 | 0.24 | 0.644 | 0.0775 | 0.583 |
| G5EE96 | 1.84 | -0.206 | -0.327 | -0.281 | 0.00542 | 0.718 | 0.568 | 0.623 |
| Q23449 | 1.84 | 1.26 | -0.334 | 1.93 | 0.00882 | 0.056 | 0.589 | 0.00672 |
| Q21884 | 1.84 | 0.611 | -0.0131 | 0.675 | 0.00667 | 0.307 | 0.982 | 0.261 |
| Q9NAB0 | 1.84 | 0.511 | 1.03 | 1.26 | 0.000592 | 0.238 | 0.0267 | 0.00897 |
| Q9XVL1 | 1.83 | 0.151 | -0.353 | 0.548 | 0.00566 | 0.791 | 0.537 | 0.343 |

| | | | | | | | | |
|---|---|---|---|---|---|---|---|---|
| A0A0K3AVS5 | 1.82 | -0.646 | -1.01 | 2.01 | 0.03 | 0.406 | 0.203 | 0.0185 |
| P90889 | 1.82 | -1.79 | 0.963 | -0.92 | 0.0498 | 0.0535 | 0.274 | 0.296 |
| Q21569 | 1.82 | 1.23 | 0.248 | 0.0226 | 0.00154 | 0.0193 | 0.601 | 0.962 |
| Q9GYK2 | 1.82 | 1.36 | 0.918 | 0.0165 | 0.0336 | 0.0989 | 0.254 | 0.983 |
| Q9GZH4 | 1.81 | -0.185 | 2.16 | 0.615 | 0.0277 | 0.805 | 0.011 | 0.417 |
| A0A4V0IK01 | 1.81 | 1.65 | 0.727 | 1.2 | 0.0109 | 0.0181 | 0.257 | 0.0705 |
| Q22508 | 1.81 | 1.11 | 0.883 | 3.31 | 0.0787 | 0.264 | 0.37 | 0.00375 |
| P34618 | 1.8 | -0.651 | -0.213 | -0.228 | 0.0172 | 0.345 | 0.754 | 0.737 |
| Q9XW92 | 1.8 | 1.29 | -0.0816 | 1.42 | 0.000208 | 0.0031 | 0.825 | 0.00159 |
| G5EC10 | 1.8 | 2.48 | -0.666 | 0.747 | 0.0546 | 0.0121 | 0.452 | 0.399 |
| Q93540 | 1.8 | 1.36 | 0.968 | 0.735 | 0.0122 | 0.0463 | 0.142 | 0.258 |
| Q20819 | 1.79 | 0.701 | -0.394 | 0.0345 | 0.00248 | 0.172 | 0.431 | 0.944 |
| Q93379 | 1.79 | 2.23 | -0.785 | 0.978 | 0.056 | 0.0215 | 0.377 | 0.275 |
| O17776 | 1.79 | 0.618 | -2.7 | -0.301 | 0.0418 | 0.451 | 0.00448 | 0.711 |
| A0A2I2LG59 | 1.79 | 1.53 | 0.379 | 0.593 | 0.048 | 0.0853 | 0.653 | 0.484 |
| Q17740 | 1.79 | 1.07 | 0.641 | 0.693 | 0.0222 | 0.147 | 0.373 | 0.337 |
| O02485 | 1.78 | 1.18 | 1.67 | 0.414 | 0.0129 | 0.0806 | 0.0182 | 0.517 |
| G5EFK4 | 1.78 | 1.17 | -1.19 | 1.94 | 0.0567 | 0.193 | 0.185 | 0.0396 |
| Q18680 | 1.77 | 0.93 | 0.407 | 0.506 | 0.000182 | 0.0188 | 0.263 | 0.17 |
| Q21633 | 1.77 | 0.652 | 0.33 | 0.846 | 0.0435 | 0.426 | 0.684 | 0.305 |
| U4PBU8 | 1.77 | 1.28 | 0.321 | 1.88 | 0.0349 | 0.114 | 0.678 | 0.0265 |
| Q03565 | 1.76 | 0.575 | -0.931 | 1.45 | 0.023 | 0.417 | 0.197 | 0.0538 |
| H2L0C8 | 1.76 | 0.517 | -1.65 | 1.51 | 0.127 | 0.642 | 0.151 | 0.186 |
| Q17994 | 1.76 | -0.558 | -1.23 | -0.253 | 0.000239 | 0.143 | 0.00426 | 0.492 |
| Q21233 | 1.76 | 0.113 | -0.591 | 0.0378 | 0.0144 | 0.861 | 0.364 | 0.953 |
| Q9BL46 | 1.76 | 1.15 | 0.346 | 1.24 | 0.00222 | 0.0282 | 0.474 | 0.0194 |
| Q9N384 | 1.75 | 1.85 | -0.616 | 1.92 | 0.0139 | 0.00996 | 0.338 | 0.00799 |
| P53588 | 1.74 | 1.09 | 0.738 | 0.516 | 0.0141 | 0.0997 | 0.253 | 0.418 |
| H2KYP2 | 1.74 | 1.31 | 0.798 | 0.569 | 0.0045 | 0.0229 | 0.142 | 0.286 |
| H2KZV5 | 1.74 | 1.3 | 0.336 | 1.43 | 0.0415 | 0.117 | 0.671 | 0.0869 |
| Q09236 | 1.73 | 0.405 | 2.65 | 2.01 | 0.0384 | 0.6 | 0.00358 | 0.0189 |
| Q21603 | 1.73 | 0.811 | 0.371 | 0.878 | 0.008 | 0.17 | 0.519 | 0.139 |
| Q20168 | 1.72 | 1.89 | 0.961 | 1.29 | 0.0124 | 0.007 | 0.13 | 0.0494 |
| O62198 | 1.72 | -0.18 | -0.137 | -0.206 | 0.0107 | 0.762 | 0.818 | 0.728 |
| H2KZJ5 | 1.72 | 0.237 | -0.239 | -0.208 | 0.0555 | 0.778 | 0.776 | 0.805 |
| Q2XN18 | 1.72 | 1.43 | 1.42 | 1.05 | 0.0325 | 0.069 | 0.0709 | 0.17 |
| O18054 | 1.71 | 1.39 | 1.67 | 1.71 | 0.0106 | 0.0311 | 0.0119 | 0.0106 |
| Q94360 | 1.71 | 0.926 | -0.225 | 0.769 | 0.00479 | 0.0913 | 0.666 | 0.154 |
| Q9GZH3 | 1.7 | -0.334 | -0.215 | -0.852 | 0.023 | 0.624 | 0.752 | 0.222 |
| H2L064 | 1.69 | 0.177 | 1.09 | -0.0917 | 0.0118 | 0.765 | 0.0821 | 0.877 |
| P54688 | 1.68 | 0.494 | 0.659 | 0.783 | 0.00101 | 0.243 | 0.127 | 0.0742 |
| Q9XXU9 | 1.68 | 1.59 | 0.187 | 1.38 | 0.000637 | 0.00102 | 0.633 | 0.00299 |
| A5HU97 | 1.68 | -0.778 | -0.407 | -1.33 | 0.0284 | 0.276 | 0.563 | 0.0735 |
| G4SPY0 | 1.68 | 0.503 | -0.746 | 0.771 | 0.00588 | 0.348 | 0.172 | 0.159 |

| | | | | | | | | |
|---|---|---|---|---|---|---|---|---|
| H2L048 | 1.67 | -0.545 | 0.897 | -0.237 | 0.0179 | 0.396 | 0.172 | 0.709 |
| G5EE67 | 1.66 | 0.917 | 0.0352 | -0.352 | 0.0265 | 0.191 | 0.959 | 0.606 |
| O45781 | 1.66 | -1.09 | -0.255 | 0.0779 | 0.00238 | 0.0291 | 0.578 | 0.864 |
| A0A1I6CM94 | 1.65 | 1.26 | 0.568 | 0.878 | 0.0156 | 0.0539 | 0.359 | 0.164 |
| H2KZS2 | 1.65 | -0.431 | 0.738 | -0.579 | 0.0436 | 0.57 | 0.336 | 0.447 |
| Q21241 | 1.64 | 0.729 | 0.856 | 1.17 | 0.00294 | 0.132 | 0.0811 | 0.0222 |
| H2L2J5 | 1.63 | 0.505 | -0.208 | 0.649 | 0.00933 | 0.367 | 0.707 | 0.251 |
| P90779 | 1.63 | 0.203 | 0.0337 | 0.0987 | 0.00184 | 0.639 | 0.938 | 0.819 |
| P91423 | 1.63 | 1.14 | -0.928 | 0.877 | 0.0107 | 0.0581 | 0.114 | 0.134 |
| P53703 | 1.62 | -0.0851 | 0.441 | 0.118 | 0.000956 | 0.829 | 0.273 | 0.766 |
| P41956 | 1.61 | 1.12 | 0.406 | -0.394 | 0.00975 | 0.055 | 0.461 | 0.475 |
| G5EFH8 | 1.6 | 1.59 | 0.443 | 1.37 | 0.0145 | 0.015 | 0.452 | 0.032 |
| Q19537 | 1.6 | 1.15 | 1.33 | 1.49 | 0.0103 | 0.0524 | 0.0275 | 0.0158 |
| Q9TXQ1 | 1.59 | 0.894 | -0.00874 | 0.29 | 0.0103 | 0.118 | 0.987 | 0.598 |
| P34526 | 1.58 | 1.03 | 0.753 | 0.549 | 0.011 | 0.0767 | 0.184 | 0.326 |
| Q9TYW1 | 1.58 | 0.964 | 0.309 | 0.667 | 0.00212 | 0.0374 | 0.473 | 0.134 |
| O45574 | 1.58 | 1.72 | 0.808 | 1.83 | 0.00367 | 0.00201 | 0.0961 | 0.00126 |
| Q9GYU3 | 1.58 | 0.721 | -0.00534 | -0.0505 | 0.00389 | 0.137 | 0.991 | 0.914 |
| D0VWM8 | 1.58 | 0.196 | 0.693 | -0.34 | 0.0706 | 0.812 | 0.406 | 0.68 |
| G5EEA2 | 1.58 | 1.19 | 2.58 | 0.897 | 0.0282 | 0.085 | 0.00136 | 0.186 |
| H9G340 | 1.58 | 0.725 | 0.472 | -0.577 | 0.0922 | 0.42 | 0.597 | 0.52 |
| Q18926 | 1.58 | 0.905 | 0.991 | 0.962 | 0.0198 | 0.152 | 0.12 | 0.13 |
| Q19064 | 1.58 | 0.415 | 0.838 | 0.648 | 0.0373 | 0.556 | 0.243 | 0.362 |
| Q05062 | 1.57 | 0.705 | 0.513 | 1.3 | 0.00141 | 0.0965 | 0.216 | 0.00542 |
| Q23330 | 1.56 | -0.982 | 0.315 | -1.31 | 0.00362 | 0.0449 | 0.49 | 0.0107 |
| P34659 | 1.56 | 1.52 | 0.533 | -0.401 | 0.0461 | 0.0501 | 0.465 | 0.581 |
| A9Z1L1 | 1.55 | 0.836 | -0.0349 | 0.617 | 0.0162 | 0.162 | 0.952 | 0.294 |
| G5EFI4 | 1.55 | 0.972 | -0.297 | 0.487 | 0.000864 | 0.0188 | 0.43 | 0.204 |
| Q9TZL8 | 1.55 | 0.992 | 0.0436 | 2.06 | 0.0816 | 0.25 | 0.959 | 0.0259 |
| O44662 | 1.54 | -1.23 | 0.603 | -1.44 | 0.0838 | 0.159 | 0.479 | 0.105 |
| P50093 | 1.54 | -2.73 | -2.01 | -3.22 | 0.0107 | 0.000136 | 0.00182 | 2.67E-05 |
| Q20546 | 1.54 | 0.539 | 0.529 | 1.04 | 0.0138 | 0.34 | 0.349 | 0.0767 |
| P55956 | 1.53 | -2.21 | 1.28 | 2.09 | 0.121 | 0.0321 | 0.188 | 0.0407 |
| Q09543 | 1.53 | 1.37 | 0.229 | 2.42 | 0.0161 | 0.0277 | 0.688 | 0.000713 |
| O76360 | 1.53 | 0.624 | 0.369 | 0.657 | 0.0945 | 0.477 | 0.672 | 0.455 |
| Q19983 | 1.52 | 1.72 | 0.612 | 0.554 | 0.00672 | 0.00292 | 0.22 | 0.264 |
| Q20330 | 1.52 | 0.538 | 0.856 | -0.398 | 0.025 | 0.387 | 0.178 | 0.52 |
| Q17417 | 1.52 | 0.935 | 0.776 | 0.44 | 0.0713 | 0.249 | 0.335 | 0.58 |
| Q20637 | 1.51 | 0.45 | 0.318 | 0.0974 | 0.0161 | 0.427 | 0.571 | 0.862 |
| O01803 | 1.51 | 0.85 | 0.115 | 1.05 | 0.138 | 0.392 | 0.906 | 0.293 |
| Q21824 | 1.5 | 0.298 | -1.09 | 0.527 | 0.0168 | 0.597 | 0.0674 | 0.355 |
| Q18198 | 1.5 | 0.653 | 0.598 | 1.37 | 0.00634 | 0.183 | 0.22 | 0.0108 |
| Q75MI6 | 1.5 | 0.562 | 0.699 | 0.5 | 0.189 | 0.612 | 0.529 | 0.652 |
| G5EE42 | 1.49 | 0.147 | 0.412 | -0.36 | 0.0152 | 0.788 | 0.455 | 0.514 |
| G5EFQ2 | 1.48 | 0.878 | 0.994 | 0.177 | 0.0116 | 0.107 | 0.0713 | 0.733 |
| Q09508 | 1.47 | -0.594 | -0.268 | -0.15 | 0.000719 | 0.102 | 0.442 | 0.665 |

| | | | | | | | | |
|---|---|---|---|---|---|---|---|---|
| Q9N4M4 | 1.47 | 0.332 | -0.2 | 0.392 | 0.00469 | 0.458 | 0.652 | 0.383 |
| Q21284 | 1.47 | 0.00976 | 0.329 | -0.194 | 0.00622 | 0.983 | 0.483 | 0.677 |
| O02158 | 1.47 | 1.67 | -0.181 | 0.0229 | 0.111 | 0.0744 | 0.837 | 0.979 |
| Q9U296 | 1.47 | 1.04 | -0.196 | 2.54 | 0.227 | 0.385 | 0.868 | 0.0467 |
| P34369 | 1.46 | 0.401 | 1.01 | 0.213 | 0.00341 | 0.348 | 0.0285 | 0.614 |
| P90735 | 1.44 | 0.54 | 0.155 | 0.651 | 0.000663 | 0.124 | 0.645 | 0.0686 |
| P91871 | 1.43 | 0.165 | 0.245 | -0.173 | 0.015 | 0.753 | 0.64 | 0.741 |
| Q23597 | 1.43 | -0.464 | -0.295 | 0.0693 | 0.00289 | 0.261 | 0.469 | 0.864 |
| Q21193 | 1.42 | 0.999 | 0.825 | 1.07 | 0.000984 | 0.011 | 0.0297 | 0.0072 |
| P90893 | 1.42 | 1.91 | 0.779 | 1.9 | 0.0669 | 0.0187 | 0.295 | 0.019 |
| Q22763 | 1.42 | 1.16 | 2.57 | 0.414 | 0.0845 | 0.149 | 0.00464 | 0.596 |
| Q8MXR2 | 1.42 | 0.293 | 0.116 | 0.384 | 0.0777 | 0.7 | 0.879 | 0.614 |
| Q9XVZ2 | 1.42 | 0.534 | 1.31 | 0.505 | 0.071 | 0.475 | 0.0925 | 0.498 |
| Q18599 | 1.41 | -2.32 | 0.0429 | -1.72 | 0.00116 | 1.11E-05 | 0.903 | 0.000215 |
| Q22038 | 1.41 | 1.48 | -1.3 | -0.927 | 0.207 | 0.187 | 0.241 | 0.398 |
| Q8IAA5 | 1.4 | 0.0597 | -0.237 | -0.393 | 0.00922 | 0.899 | 0.616 | 0.41 |
| Q09541 | 1.4 | 1.36 | 0.86 | 1.65 | 0.0237 | 0.0275 | 0.141 | 0.00999 |
| G5EF32 | 1.4 | 0.459 | 0.134 | 0.419 | 0.0187 | 0.399 | 0.803 | 0.44 |
| P90900 | 1.39 | -1.16 | 0.167 | -1.24 | 0.116 | 0.185 | 0.843 | 0.159 |
| G5EC71 | 1.39 | 1.19 | -0.483 | 0.891 | 0.112 | 0.168 | 0.564 | 0.294 |
| P12844 | 1.38 | 0.451 | 0.516 | 0.0282 | 0.000938 | 0.192 | 0.139 | 0.933 |
| P17329 | 1.38 | -0.404 | -0.229 | 0.0488 | 0.000331 | 0.188 | 0.446 | 0.869 |
| P55954 | 1.38 | -2.1 | -2.43 | -1.85 | 0.156 | 0.0384 | 0.0194 | 0.064 |
| Q23551 | 1.37 | 0.96 | 0.603 | -0.227 | 0.00705 | 0.0435 | 0.185 | 0.608 |
| A0A168H9W7 | 1.37 | 0.627 | 1.44 | 0.00507 | 0.0108 | 0.2 | 0.00808 | 0.991 |
| G5EDV3 | 1.37 | 1.38 | -1.33 | -0.279 | 0.179 | 0.177 | 0.191 | 0.777 |
| Q22620 | 1.37 | 2.47 | 2.33 | 1.4 | 0.0688 | 0.00321 | 0.00476 | 0.0637 |
| Q9BMU4 | 1.36 | 0.13 | -0.0811 | 0.779 | 0.0528 | 0.842 | 0.901 | 0.244 |
| O45394 | 1.35 | 0.813 | 0.165 | 0.345 | 0.00205 | 0.0388 | 0.65 | 0.349 |
| O62327 | 1.33 | 0.417 | 0.582 | 0.241 | 0.000955 | 0.212 | 0.0895 | 0.461 |
| P34690 | 1.33 | 1.58 | -0.201 | 1.52 | 0.015 | 0.00543 | 0.682 | 0.00679 |
| O01824 | 1.33 | 1.4 | 0.63 | 0.861 | 0.0363 | 0.0288 | 0.289 | 0.155 |
| Q95Y96 | 1.33 | 0.858 | 0.285 | 0.638 | 0.0647 | 0.216 | 0.674 | 0.352 |
| Q93714 | 1.32 | -0.14 | 0.0709 | 0.194 | 0.00792 | 0.746 | 0.87 | 0.655 |
| Q56VX9 | 1.32 | 3.85 | 1.05 | 3.33 | 0.0132 | 1.04E-06 | 0.0415 | 5.36E-06 |
| Q20779 | 1.31 | -0.272 | -0.713 | 0.0738 | 0.0443 | 0.652 | 0.248 | 0.902 |
| H2L2C9 | 1.31 | 1.74 | -0.0945 | 0.7 | 0.0115 | 0.00177 | 0.837 | 0.143 |
| Q965T2 | 1.31 | 0.476 | -0.198 | 0.498 | 0.0602 | 0.47 | 0.763 | 0.451 |
| Q9U3H4 | 1.3 | -0.0118 | 0.541 | 0.242 | 0.0104 | 0.979 | 0.237 | 0.589 |
| B6VQ62 | 1.3 | 1.43 | 0.208 | 0.53 | 0.0411 | 0.0265 | 0.724 | 0.373 |
| G5ECK7 | 1.3 | 0.976 | 0.43 | 0.799 | 0.0299 | 0.0918 | 0.438 | 0.16 |
| G5EF53 | 1.29 | 0.731 | 0.0107 | 0.164 | 0.018 | 0.152 | 0.983 | 0.739 |
| O17680 | 1.28 | 0.987 | 0.871 | 0.911 | 0.00111 | 0.00692 | 0.0144 | 0.0112 |
| Q22101 | 1.28 | -2.27 | 0.532 | -0.735 | 0.00412 | 3.04E-05 | 0.176 | 0.069 |
| G5EFJ3 | 1.28 | 0.1 | 0.21 | 0.642 | 0.0571 | 0.874 | 0.739 | 0.316 |
| Q18817 | 1.28 | 1.68 | 1.93 | 0.242 | 0.0615 | 0.0184 | 0.00845 | 0.707 |

| | | | | | | | | |
|---|---|---|---|---|---|---|---|---|
| Q20049 | 1.28 | 0.47 | 0.191 | 0.301 | 0.0395 | 0.416 | 0.738 | 0.6 |
| O01592 | 1.27 | -0.736 | -0.745 | -0.244 | 0.069 | 0.272 | 0.266 | 0.71 |
| P90868 | 1.27 | 2.24 | -1.49 | 2.07 | 0.0343 | 0.000999 | 0.0153 | 0.00183 |
| A0A0K3AQS9 | 1.27 | 0.44 | -0.0782 | 0.107 | 0.0587 | 0.488 | 0.901 | 0.865 |
| G5EDC6 | 1.27 | 0.338 | 0.666 | 0.269 | 0.0639 | 0.6 | 0.309 | 0.676 |
| G5EED5 | 1.27 | -0.627 | 0.876 | 0.214 | 0.188 | 0.505 | 0.356 | 0.819 |
| Q564Q1 | 1.26 | -0.367 | -0.0027 | 0.387 | 0.0221 | 0.465 | 0.996 | 0.442 |
| Q9N4F3 | 1.26 | 0.249 | -0.484 | 0.819 | 0.0532 | 0.682 | 0.43 | 0.191 |
| B2D6P1 | 1.26 | 1.21 | -0.555 | -0.29 | 0.14 | 0.157 | 0.503 | 0.725 |
| Q18227 | 1.26 | 1.24 | 1.21 | 0.911 | 0.0624 | 0.0664 | 0.0719 | 0.165 |
| Q21544 | 1.25 | -1.49 | -1.22 | -0.15 | 0.0895 | 0.0479 | 0.0953 | 0.829 |
| Q22562 | 1.25 | 0.766 | 0.279 | 0.929 | 0.00224 | 0.038 | 0.417 | 0.0149 |
| A0A0K3ARC0 | 1.24 | 0.539 | -0.851 | 0.341 | 0.0834 | 0.429 | 0.22 | 0.614 |
| O17695 | 1.24 | 0.503 | -0.505 | 0.296 | 0.188 | 0.584 | 0.582 | 0.746 |
| P46975 | 1.24 | 1.67 | -0.281 | 0.988 | 0.144 | 0.0563 | 0.731 | 0.237 |
| Q19420 | 1.24 | 1.36 | -0.0565 | 0.146 | 0.0729 | 0.052 | 0.931 | 0.822 |
| G5EGP8 | 1.23 | -1.52 | -2.06 | -2.04 | 0.144 | 0.0767 | 0.0209 | 0.0222 |
| Q22498 | 1.23 | 0.354 | -0.385 | 0.481 | 0.0395 | 0.525 | 0.49 | 0.39 |
| Q09665 | 1.22 | 0.267 | -1.74 | 0.501 | 0.0848 | 0.691 | 0.0196 | 0.459 |
| Q9XWT3 | 1.22 | -0.654 | 0.728 | 0.621 | 0.189 | 0.47 | 0.422 | 0.492 |
| P34686 | 1.21 | -1.17 | 0.283 | -1.27 | 0.114 | 0.125 | 0.699 | 0.0997 |
| Q10454 | 1.21 | 0.622 | -0.176 | 0.565 | 0.00179 | 0.0684 | 0.585 | 0.0947 |
| Q17686 | 1.21 | 3.14 | -1.66 | 1.6 | 0.0615 | 0.000125 | 0.0148 | 0.0178 |
| Q22020 | 1.21 | 0.814 | -0.0676 | -0.215 | 0.0252 | 0.114 | 0.891 | 0.663 |
| Q09422 | 1.21 | 0.655 | 0.703 | 0.558 | 0.085 | 0.331 | 0.298 | 0.406 |
| Q9TXX0 | 1.21 | 0.695 | 0.264 | 0.581 | 0.128 | 0.367 | 0.729 | 0.45 |
| Q21888 | 1.2 | 1.52 | 0.51 | 0.641 | 0.0332 | 0.0096 | 0.33 | 0.226 |
| Q93896 | 1.19 | 0.449 | -0.0712 | -0.643 | 0.0183 | 0.331 | 0.875 | 0.171 |
| O01578 | 1.18 | 0.204 | -0.877 | -0.125 | 0.0565 | 0.724 | 0.144 | 0.829 |
| O61199 | 1.18 | -1.67 | 0.069 | -0.629 | 0.0849 | 0.0205 | 0.915 | 0.34 |
| P12845 | 1.17 | 0.326 | 1.11 | 0.205 | 0.00826 | 0.406 | 0.0112 | 0.598 |
| Q93761 | 1.17 | -0.789 | -2.06 | -0.549 | 0.0254 | 0.114 | 0.000637 | 0.261 |
| Q20684 | 1.17 | -1.91 | -0.569 | -0.851 | 0.0102 | 0.000272 | 0.171 | 0.0488 |
| Q09450 | 1.17 | -0.183 | -0.111 | 0.513 | 0.0588 | 0.751 | 0.848 | 0.381 |
| G5EEQ8 | 1.16 | 0.0413 | 2.22 | -0.672 | 0.0989 | 0.95 | 0.00447 | 0.322 |
| G5EER0 | 1.16 | -1.46 | -0.222 | -2.13 | 0.203 | 0.116 | 0.802 | 0.0281 |
| Q18550 | 1.16 | 1.85 | 1.14 | 1.05 | 0.0155 | 0.000628 | 0.0172 | 0.026 |
| O01805 | 1.15 | 0.818 | -0.304 | 0.865 | 0.0403 | 0.13 | 0.559 | 0.11 |
| H1UBK1 | 1.15 | -0.0637 | -0.862 | -0.391 | 0.176 | 0.938 | 0.302 | 0.634 |
| A0A679L8M9 | 1.15 | -0.188 | 0.379 | -0.166 | 0.0761 | 0.758 | 0.537 | 0.786 |
| O76687 | 1.15 | 0.868 | 0.037 | -0.00937 | 0.0315 | 0.0934 | 0.94 | 0.985 |
| Q18280 | 1.14 | -0.229 | 0.21 | -0.129 | 0.0158 | 0.589 | 0.62 | 0.76 |
| E8MDW0 | 1.14 | 0.794 | 0.739 | 1.13 | 0.106 | 0.248 | 0.281 | 0.107 |
| G5EBK3 | 1.13 | 0.72 | 0.341 | 0.579 | 0.0146 | 0.0966 | 0.412 | 0.174 |

| | | | | | | | | |
|---|---|---|---|---|---|---|---|---|
| Q95PZ7 | 1.13 | 3.35 | 0.594 | 3.38 | 0.198 | 0.00131 | 0.488 | 0.00122 |
| P34669 | 1.13 | 0.54 | 0.512 | 0.789 | 0.035 | 0.282 | 0.307 | 0.125 |
| Q9TZH6 | 1.13 | 0.85 | 1.36 | 0.714 | 0.265 | 0.397 | 0.184 | 0.475 |
| O17836 | 1.12 | 1.75 | 0.446 | 1.58 | 0.0125 | 0.000546 | 0.273 | 0.00123 |
| Q21217 | 1.12 | 0.607 | -0.597 | -0.767 | 0.211 | 0.489 | 0.497 | 0.385 |
| Q9TXH9 | 1.12 | 0.674 | 1.4 | 0.0386 | 0.114 | 0.326 | 0.0523 | 0.954 |
| O76430 | 1.12 | 1.25 | 0.972 | 0.465 | 0.116 | 0.0822 | 0.169 | 0.498 |
| Q21962 | 1.12 | 0.115 | 0.659 | 0.0973 | 0.0533 | 0.832 | 0.236 | 0.857 |
| P49632 | 1.11 | 0.508 | 0.0653 | 1.29 | 0.00576 | 0.157 | 0.85 | 0.00204 |
| Q10657 | 1.11 | 0.186 | -0.648 | 0.406 | 0.014 | 0.645 | 0.123 | 0.321 |
| Q20719 | 1.11 | -0.178 | -0.409 | 0.518 | 0.201 | 0.832 | 0.627 | 0.539 |
| Q22170 | 1.11 | 0.631 | 0.333 | -0.226 | 0.0459 | 0.232 | 0.52 | 0.662 |
| Q86NC2 | 1.11 | 0.46 | -0.137 | 0.463 | 0.0561 | 0.402 | 0.801 | 0.399 |
| P50306 | 1.1 | 1.22 | 0.305 | 0.839 | 0.0398 | 0.0246 | 0.538 | 0.105 |
| P51875 | 1.1 | -0.192 | -0.137 | 0.552 | 0.0924 | 0.757 | 0.824 | 0.379 |
| A0A486WX07 | 1.1 | 1.15 | 1.84 | 0.663 | 0.17 | 0.155 | 0.0303 | 0.399 |
| V6CLP5 | 1.09 | 0.403 | 0.77 | 0.0587 | 0.00642 | 0.255 | 0.0398 | 0.865 |
| O44782 | 1.09 | 0.0709 | 1.53 | 0.678 | 0.0623 | 0.897 | 0.013 | 0.228 |
| Q9U2Z1 | 1.09 | 1.36 | 0.761 | 0.813 | 0.168 | 0.0919 | 0.329 | 0.298 |
| P02567 | 1.08 | 0.499 | 0.855 | 0.252 | 0.0142 | 0.218 | 0.0443 | 0.525 |
| P52713 | 1.08 | -0.318 | -0.138 | 0.311 | 0.00746 | 0.37 | 0.693 | 0.381 |
| Q20053 | 1.08 | 0.508 | -0.636 | 1.25 | 0.0502 | 0.329 | 0.226 | 0.0264 |
| Q21752 | 1.08 | -0.636 | -1.58 | -0.106 | 0.0108 | 0.106 | 0.000786 | 0.777 |
| O17218 | 1.08 | 2.29 | -0.412 | 2.18 | 0.113 | 0.00305 | 0.53 | 0.0043 |
| Q95ZL1 | 1.08 | -0.0711 | 0.235 | 0.16 | 0.0106 | 0.849 | 0.532 | 0.67 |
| D5MCN2 | 1.08 | 0.197 | 0.39 | 1.04 | 0.145 | 0.782 | 0.587 | 0.158 |
| O44400 | 1.07 | 0.891 | 0.488 | 0.529 | 0.00805 | 0.0222 | 0.18 | 0.149 |
| P41938 | 1.07 | 0.246 | -0.982 | 0.686 | 0.0508 | 0.632 | 0.0712 | 0.193 |
| Q20173 | 1.07 | 0.59 | -0.863 | 0.339 | 0.0124 | 0.137 | 0.0369 | 0.38 |
| Q18688 | 1.06 | 2.03 | -0.0211 | 1.96 | 0.0233 | 0.000248 | 0.96 | 0.00033 |
| Q03577 | 1.06 | 1.51 | 0.114 | 0.643 | 0.185 | 0.066 | 0.883 | 0.41 |
| Q21831 | 1.06 | 0.691 | 0.675 | -0.443 | 0.142 | 0.326 | 0.337 | 0.524 |
| A0A4V0IJD5 | 1.05 | 0.709 | 0.337 | -0.203 | 0.0282 | 0.121 | 0.446 | 0.644 |
| Q20224 | 1.05 | 0.568 | 0.164 | 1.85 | 0.139 | 0.409 | 0.81 | 0.0151 |
| P31161 | 1.04 | -0.622 | -1.02 | -0.0113 | 0.035 | 0.185 | 0.0382 | 0.98 |
| Q9U1W1 | 1.04 | -2.86 | 1.81 | -2.5 | 0.348 | 0.0181 | 0.112 | 0.0346 |
| A0A0K3AT05 | 1.04 | 0.646 | 0.154 | -0.295 | 0.0353 | 0.168 | 0.733 | 0.517 |
| A8WFK2 | 1.04 | -1.85 | -2.05 | -1.94 | 0.315 | 0.0857 | 0.0588 | 0.0724 |
| Q19007 | 1.04 | 2.05 | 0.0228 | 1.9 | 0.0167 | 0.000103 | 0.953 | 0.000204 |
| O02058 | 1.04 | 0.724 | 0.114 | 0.0629 | 0.0612 | 0.176 | 0.826 | 0.903 |
| A0A0K3AUJ9 | 1.03 | -0.201 | 0.284 | -0.147 | 0.0062 | 0.539 | 0.388 | 0.652 |
| Q21355 | 1.03 | -1.71 | 1.46 | -2.04 | 0.189 | 0.0371 | 0.0696 | 0.016 |
| O45443 | 1.03 | -0.454 | -1.2 | -1.05 | 0.0647 | 0.393 | 0.0357 | 0.0621 |

| | | | | | | | | |
|---|---|---|---|---|---|---|---|---|
| Q4R127 | 1.03 | -1.87 | 0.86 | -2.25 | 0.0871 | 0.0049 | 0.147 | 0.00132 |
| P34559 | 1.03 | -0.263 | -0.449 | -0.0332 | 0.174 | 0.721 | 0.543 | 0.964 |
| Q10943 | 1.02 | 1.4 | -1.47 | 1.67 | 0.118 | 0.0383 | 0.0306 | 0.0167 |
| Q9XUT9 | 1.02 | 0.845 | -1.52 | 0.457 | 0.181 | 0.264 | 0.0546 | 0.539 |
| Q17967 | 1.01 | 0.81 | -0.23 | 1.28 | 0.0286 | 0.0712 | 0.587 | 0.00808 |
| Q20576 | 1.01 | 0.904 | 0.791 | 0.902 | 0.0637 | 0.0922 | 0.136 | 0.0929 |
| Q18823 | 1 | 0.974 | 1.56 | -0.254 | 0.15 | 0.16 | 0.0326 | 0.705 |
| G5EG13 | 1 | -0.0709 | 0.268 | -0.611 | 0.242 | 0.932 | 0.748 | 0.468 |
| Q9U229 | 1 | -2.12 | 0.0179 | -1.94 | 0.389 | 0.0802 | 0.988 | 0.108 |
| O02495 | 0.993 | -0.589 | 0.168 | 0.369 | 0.101 | 0.315 | 0.771 | 0.524 |
| Q9N4Z0 | 0.992 | 0.253 | -0.0308 | -0.29 | 0.301 | 0.788 | 0.974 | 0.758 |
| O18178 | 0.989 | 0.867 | 0.523 | 0.495 | 0.204 | 0.262 | 0.492 | 0.515 |
| P10567 | 0.988 | 0.211 | -0.731 | 0.0678 | 0.0515 | 0.656 | 0.137 | 0.886 |
| P43510 | 0.987 | 0.384 | -0.646 | 0.451 | 0.0942 | 0.496 | 0.259 | 0.426 |
| P27639 | 0.985 | 2.46 | 0.156 | 2.02 | 0.0651 | 0.000205 | 0.755 | 0.00109 |
| O01812 | 0.984 | 0.363 | -0.215 | 0.57 | 0.0236 | 0.364 | 0.587 | 0.163 |
| Q9N5S7 | 0.984 | 0.563 | 0.346 | 0.361 | 0.133 | 0.377 | 0.584 | 0.567 |
| O02365 | 0.977 | 0.447 | 0.196 | -0.892 | 0.0381 | 0.311 | 0.653 | 0.0551 |
| Q09359 | 0.977 | 0.731 | 0.28 | 0.651 | 0.101 | 0.21 | 0.622 | 0.261 |
| Q18120 | 0.974 | 0.764 | 0.352 | 0.348 | 0.0527 | 0.119 | 0.456 | 0.462 |
| O45864 | 0.972 | -1.83 | -0.582 | 0.0563 | 0.301 | 0.0631 | 0.53 | 0.951 |
| Q20107 | 0.972 | 1.17 | -1.55 | 1.13 | 0.154 | 0.0911 | 0.0304 | 0.102 |
| Q23604 | 0.967 | 0.21 | -0.321 | -0.681 | 0.0993 | 0.707 | 0.567 | 0.234 |
| C1P636 | 0.965 | 1.76 | 0.703 | 1.92 | 0.0597 | 0.00224 | 0.158 | 0.00114 |
| K8ESM2 | 0.961 | 0.301 | -0.754 | -0.701 | 0.291 | 0.736 | 0.404 | 0.437 |
| G5EGP4 | 0.96 | 0.744 | 0.157 | 0.52 | 0.0643 | 0.142 | 0.747 | 0.295 |
| Q9XUL7 | 0.958 | 1.07 | -0.608 | 0.218 | 0.261 | 0.212 | 0.469 | 0.793 |
| O76836 | 0.952 | 0.862 | -0.187 | 1.16 | 0.209 | 0.253 | 0.8 | 0.132 |
| Q9N3D9 | 0.95 | -0.272 | 1.22 | -0.56 | 0.269 | 0.747 | 0.16 | 0.508 |
| O01530 | 0.93 | -0.427 | 0.11 | 0.449 | 0.0414 | 0.32 | 0.794 | 0.296 |
| Q19853 | 0.925 | 0.97 | -1.5 | 0.638 | 0.213 | 0.193 | 0.0529 | 0.383 |
| P28548 | 0.925 | 0.507 | 0.589 | -0.0917 | 0.125 | 0.385 | 0.316 | 0.874 |
| Q9BL34 | 0.921 | -1.29 | -1.62 | -1.26 | 0.212 | 0.0874 | 0.0377 | 0.0958 |
| Q18276 | 0.92 | -0.562 | -0.67 | 0.979 | 0.419 | 0.619 | 0.554 | 0.39 |
| A0A4V0IIR6 | 0.915 | 1.31 | -0.717 | 0.889 | 0.0524 | 0.00902 | 0.119 | 0.0585 |
| P46502 | 0.913 | -0.22 | -0.561 | -0.331 | 0.141 | 0.713 | 0.354 | 0.581 |
| P21137 | 0.912 | 1.29 | 0.21 | 1.6 | 0.124 | 0.0373 | 0.713 | 0.0126 |
| P20163 | 0.903 | -1.14 | 0.993 | -0.61 | 0.248 | 0.149 | 0.206 | 0.429 |
| O44565 | 0.902 | 0.0521 | 0.714 | -0.802 | 0.235 | 0.944 | 0.343 | 0.289 |
| Q9XTB5 | 0.893 | 0.263 | 1.24 | 0.532 | 0.296 | 0.754 | 0.154 | 0.528 |
| O62146 | 0.891 | -1.22 | -0.642 | -0.653 | 0.171 | 0.0675 | 0.316 | 0.308 |
| Q22781 | 0.889 | -0.176 | -0.689 | 0.318 | 0.081 | 0.716 | 0.167 | 0.512 |
| A0A5E4LXR0 | 0.888 | -0.749 | 1.96 | 0.295 | 0.119 | 0.183 | 0.00261 | 0.589 |
| Q9N4J8 | 0.886 | 2.56 | 1.22 | 2.19 | 0.0929 | 0.000139 | 0.0261 | 0.000555 |
| P91856 | 0.879 | 0.96 | -0.33 | 0.954 | 0.0618 | 0.0438 | 0.458 | 0.045 |

| | | | | | | | | |
|---|---|---|---|---|---|---|---|---|
| Q17754 | 0.878 | -0.399 | -0.816 | -0.708 | 0.319 | 0.646 | 0.354 | 0.419 |
| P30625 | 0.876 | 0.367 | 0.464 | 0.268 | 0.0328 | 0.337 | 0.23 | 0.479 |
| Q19240 | 0.867 | -1.97 | -0.81 | -2.22 | 0.303 | 0.0295 | 0.334 | 0.0162 |
| P08898 | 0.866 | 0.925 | -1.09 | 0.844 | 0.0934 | 0.0752 | 0.0405 | 0.101 |
| Q8T3D2 | 0.865 | 1.12 | 0.608 | 0.138 | 0.278 | 0.166 | 0.441 | 0.86 |
| Q20634 | 0.863 | -3.11 | 0.00183 | -3.86 | 0.0485 | 2.06E-06 | 0.996 | 1.61E-07 |
| G5EEL9 | 0.859 | 0.236 | 0.412 | -0.994 | 0.102 | 0.638 | 0.415 | 0.0622 |
| P34685 | 0.858 | 0.763 | 0.766 | -0.00394 | 0.096 | 0.135 | 0.133 | 0.994 |
| O01816 | 0.854 | 0.384 | -0.448 | 0.931 | 0.0518 | 0.355 | 0.283 | 0.036 |
| Q7Z072 | 0.854 | -1.4 | 0.187 | -0.9 | 0.066 | 0.00555 | 0.669 | 0.0543 |
| G4S034 | 0.848 | 0.491 | 0.0347 | 0.117 | 0.0168 | 0.138 | 0.913 | 0.714 |
| Q9XVM0 | 0.846 | 0.478 | -0.0638 | 0.31 | 0.251 | 0.51 | 0.929 | 0.667 |
| Q19626 | 0.84 | 0.418 | -0.218 | 0.396 | 0.0125 | 0.175 | 0.468 | 0.198 |
| O45679 | 0.838 | 0.519 | 0.325 | 0.603 | 0.0189 | 0.122 | 0.32 | 0.0766 |
| P34339 | 0.83 | 1.48 | 0.894 | 1.29 | 0.413 | 0.154 | 0.379 | 0.212 |
| Q9XUE5 | 0.824 | 0.344 | 0.349 | 0.782 | 0.207 | 0.59 | 0.585 | 0.23 |
| Q23588 | 0.823 | 0.949 | -1.18 | 1.13 | 0.256 | 0.194 | 0.112 | 0.126 |
| O17915 | 0.822 | 1.87 | 0.21 | 1.4 | 0.0899 | 0.00102 | 0.648 | 0.00783 |
| Q9NA75 | 0.818 | 0.198 | -0.139 | -0.0795 | 0.172 | 0.732 | 0.81 | 0.891 |
| Q20228 | 0.817 | 2.35 | -0.439 | 2.1 | 0.208 | 0.00199 | 0.49 | 0.0044 |
| Q21000 | 0.815 | -1.98 | 1.18 | -1.47 | 0.138 | 0.00192 | 0.0391 | 0.0131 |
| P02566 | 0.815 | 0.466 | 0.294 | 0.229 | 0.0239 | 0.169 | 0.375 | 0.487 |
| Q9NDC9 | 0.811 | 0.397 | 0.68 | 0.795 | 0.271 | 0.584 | 0.353 | 0.281 |
| P62784 | 0.806 | 0.326 | -0.518 | 0.791 | 0.0991 | 0.486 | 0.275 | 0.105 |
| Q18211 | 0.801 | -1.75 | 0.717 | -2.1 | 0.115 | 0.0025 | 0.154 | 0.000598 |
| Q11176 | 0.799 | 0.391 | 0.106 | -0.658 | 0.337 | 0.634 | 0.897 | 0.427 |
| H2KYN6 | 0.797 | 0.635 | 0.687 | 0.783 | 0.191 | 0.292 | 0.256 | 0.198 |
| Q9XXR4 | 0.796 | -2.05 | 0.784 | -1.9 | 0.153 | 0.00162 | 0.158 | 0.00291 |
| Q86NH9 | 0.796 | 0.463 | 0.755 | -0.0543 | 0.0791 | 0.289 | 0.0943 | 0.899 |
| Q9BLB6 | 0.793 | -1.69 | -0.186 | -1.73 | 0.449 | 0.118 | 0.858 | 0.111 |
| Q19058 | 0.781 | -2.92 | -0.827 | -2.99 | 0.15 | 5.91E-05 | 0.129 | 4.65E-05 |
| Q19749 | 0.78 | -0.382 | -0.107 | -0.159 | 0.0362 | 0.275 | 0.755 | 0.644 |
| Q20062 | 0.778 | -0.246 | -0.146 | 0.0924 | 0.16 | 0.646 | 0.785 | 0.862 |
| G5EDP2 | 0.775 | -0.395 | 0.000992 | -0.721 | 0.0954 | 0.377 | 0.998 | 0.119 |
| P91455 | 0.775 | -0.738 | -2.11 | -1.07 | 0.427 | 0.449 | 0.0427 | 0.277 |
| G5EES3 | 0.762 | -0.569 | 0.189 | -0.134 | 0.234 | 0.369 | 0.762 | 0.83 |
| O17759 | 0.76 | -0.274 | 0.295 | 0.0302 | 0.0697 | 0.49 | 0.457 | 0.939 |
| Q9BIC3 | 0.757 | -0.322 | -0.284 | -0.216 | 0.0526 | 0.382 | 0.44 | 0.556 |
| Q22100 | 0.756 | -1.89 | -0.32 | -1.06 | 0.405 | 0.0497 | 0.722 | 0.247 |
| G5ECA7 | 0.751 | -0.0673 | -0.559 | -0.209 | 0.124 | 0.885 | 0.243 | 0.655 |
| Q22494 | 0.749 | 0.748 | 1.06 | 0.364 | 0.0744 | 0.0747 | 0.0165 | 0.364 |
| Q9N5E4 | 0.742 | -1.29 | -1.74 | -1.45 | 0.264 | 0.063 | 0.0165 | 0.0388 |
| O01799 | 0.74 | 0.136 | -0.143 | -1.26 | 0.341 | 0.859 | 0.852 | 0.116 |
| Q17761 | 0.739 | 1.12 | -1.3 | 0.171 | 0.103 | 0.0191 | 0.0084 | 0.693 |
| O01302 | 0.737 | 0.625 | -0.335 | 0.546 | 0.162 | 0.231 | 0.513 | 0.292 |
| P52717 | 0.735 | 0.756 | -0.0718 | 1.43 | 0.136 | 0.126 | 0.879 | 0.00812 |
| O02108 | 0.735 | -0.453 | -0.717 | 0.401 | 0.292 | 0.511 | 0.304 | 0.56 |

| | | | | | | | | |
|---|---|---|---|---|---|---|---|---|
| Q9N4H7 | 0.734 | 1.19 | 2.21 | 0.962 | 0.141 | 0.0245 | 0.000352 | 0.0602 |
| Q95QS3 | 0.731 | 2.39 | 0.204 | 3.04 | 0.269 | 0.00214 | 0.753 | 0.000298 |
| Q9XXF9 | 0.727 | 0.757 | -0.214 | -0.786 | 0.215 | 0.198 | 0.709 | 0.182 |
| G5EBJ7 | 0.726 | 0.562 | 0.0698 | 0.521 | 0.0812 | 0.168 | 0.859 | 0.199 |
| O16487 | 0.725 | 0.358 | 0.0601 | 0.101 | 0.134 | 0.445 | 0.897 | 0.828 |
| H2KYQ5 | 0.723 | 0.19 | 0.529 | -0.125 | 0.0472 | 0.576 | 0.133 | 0.712 |
| Q9XXK1 | 0.72 | -0.258 | -0.404 | -0.031 | 0.0191 | 0.358 | 0.159 | 0.911 |
| P46563 | 0.718 | 0.273 | -0.366 | 0.101 | 0.0403 | 0.404 | 0.268 | 0.755 |
| O18000 | 0.715 | 0.85 | -1.93 | 1.15 | 0.0622 | 0.0304 | 8.56E-05 | 0.00562 |
| Q27874 | 0.713 | 0.907 | 0.282 | 0.706 | 0.36 | 0.249 | 0.714 | 0.365 |
| P91283 | 0.713 | -0.932 | -0.416 | 0.557 | 0.491 | 0.371 | 0.686 | 0.589 |
| Q9XTZ2 | 0.713 | -1.66 | -0.117 | -0.13 | 0.461 | 0.1 | 0.903 | 0.892 |
| Q9XVJ3 | 0.713 | -2.12 | -2.19 | -2.47 | 0.582 | 0.116 | 0.105 | 0.0712 |
| Q9GS00 | 0.712 | 0.249 | -0.218 | -0.0724 | 0.189 | 0.637 | 0.679 | 0.89 |
| P34455 | 0.71 | -1.94 | -0.514 | -0.893 | 0.0449 | 3.30E-05 | 0.133 | 0.0152 |
| Q18212 | 0.705 | 1.6 | -0.4 | 1.59 | 0.124 | 0.00234 | 0.369 | 0.00255 |
| O45552 | 0.699 | -3.95 | -0.749 | -2.05 | 0.226 | 5.20E-06 | 0.196 | 0.00232 |
| Q95XS1 | 0.697 | 0.94 | 0.757 | 0.598 | 0.355 | 0.218 | 0.316 | 0.425 |
| Q9TYS3 | 0.693 | 0.28 | -0.171 | 0.433 | 0.0973 | 0.484 | 0.668 | 0.286 |
| O45444 | 0.69 | 0.615 | -0.583 | 0.783 | 0.0656 | 0.097 | 0.114 | 0.0399 |
| O02323 | 0.687 | 1.14 | -0.233 | 0.541 | 0.159 | 0.0278 | 0.622 | 0.261 |
| Q20655 | 0.686 | 0.319 | -0.0197 | 0.549 | 0.0564 | 0.349 | 0.953 | 0.118 |
| B9WRT9 | 0.682 | -0.214 | 1.97 | -1.06 | 0.133 | 0.625 | 0.000427 | 0.0265 |
| Q22799 | 0.679 | -0.869 | -0.765 | 1.12 | 0.339 | 0.226 | 0.283 | 0.124 |
| A0A2C9C 377 | 0.678 | 0.485 | 1.4 | 0.447 | 0.308 | 0.461 | 0.046 | 0.496 |
| G5EE90 | 0.677 | 0.995 | -0.236 | -0.184 | 0.171 | 0.0524 | 0.622 | 0.701 |
| G4RUC0 | 0.674 | 1.03 | 0.247 | 0.958 | 0.244 | 0.0846 | 0.662 | 0.106 |
| G5ECE7 | 0.674 | 1.24 | 0.094 | 1.12 | 0.315 | 0.0751 | 0.886 | 0.105 |
| Q93315 | 0.673 | 0.871 | 0.859 | 0.331 | 0.28 | 0.168 | 0.174 | 0.589 |
| P36573 | 0.665 | -0.617 | -0.448 | -0.25 | 0.0745 | 0.0957 | 0.215 | 0.48 |
| P17343 | 0.658 | 0.279 | -0.776 | 0.294 | 0.18 | 0.56 | 0.118 | 0.539 |
| P53795 | 0.656 | -0.64 | -0.736 | 0.409 | 0.374 | 0.385 | 0.32 | 0.576 |
| A4F319 | 0.651 | -0.228 | -0.2 | 1.54 | 0.396 | 0.763 | 0.791 | 0.0574 |
| Q23237 | 0.648 | 0.638 | -1.37 | 0.994 | 0.157 | 0.163 | 0.00699 | 0.0377 |
| P91253 | 0.643 | -0.471 | -1.8 | -0.265 | 0.132 | 0.261 | 0.000529 | 0.52 |
| B7FAR9 | 0.641 | -0.704 | 0.858 | 0.178 | 0.49 | 0.449 | 0.359 | 0.847 |
| P52899 | 0.637 | -2.01 | -0.758 | -0.363 | 0.326 | 0.00626 | 0.246 | 0.57 |
| Q09285 | 0.637 | 0.219 | -0.167 | 0.247 | 0.34 | 0.739 | 0.8 | 0.707 |
| P91859 | 0.634 | -0.544 | 0.839 | -0.58 | 0.421 | 0.488 | 0.291 | 0.461 |
| P52275 | 0.632 | 1.52 | -0.293 | 1.01 | 0.142 | 0.00223 | 0.483 | 0.0256 |
| Q09584 | 0.631 | -0.119 | 2.66 | -1.12 | 0.501 | 0.898 | 0.0114 | 0.239 |
| Q95QW0 | 0.629 | 1.31 | -1.4 | 1.87 | 0.391 | 0.0873 | 0.0699 | 0.0201 |
| Q20507 | 0.628 | 0.612 | -2.47 | 1.1 | 0.356 | 0.368 | 0.0022 | 0.117 |
| Q21551 | 0.628 | 1.09 | 0.348 | 0.133 | 0.228 | 0.0458 | 0.496 | 0.793 |
| Q23683 | 0.624 | 4.4 | 0.458 | 3.92 | 0.41 | 3.55E-05 | 0.544 | 0.000111 |
| O01876 | 0.622 | 0.482 | -0.365 | 0.0785 | 0.41 | 0.522 | 0.626 | 0.916 |

| | | | | | | | | |
|---|---|---|---|---|---|---|---|---|
| Q9GYF1 | 0.621 | 0.361 | -0.473 | 0.0869 | 0.134 | 0.371 | 0.246 | 0.827 |
| Q18036 | 0.618 | -0.84 | -1.65 | -0.687 | 0.241 | 0.118 | 0.00576 | 0.195 |
| P49595 | 0.618 | -0.544 | 0.828 | -0.394 | 0.326 | 0.385 | 0.194 | 0.527 |
| O02328 | 0.611 | 1.52 | -0.248 | -0.712 | 0.352 | 0.0315 | 0.702 | 0.281 |
| Q21313 | 0.609 | 0.809 | 0.302 | 0.245 | 0.0816 | 0.0261 | 0.367 | 0.462 |
| Q9XWL1 | 0.607 | -0.9 | 0.192 | -1.09 | 0.185 | 0.0581 | 0.667 | 0.0259 |
| Q2HQL4 | 0.605 | -0.101 | 0.365 | -0.518 | 0.411 | 0.889 | 0.617 | 0.479 |
| Q09510 | 0.604 | -0.149 | -0.538 | -0.222 | 0.333 | 0.808 | 0.387 | 0.718 |
| K8FDY0 | 0.603 | -1.98 | -0.215 | -2.59 | 0.321 | 0.00466 | 0.719 | 0.000613 |
| Q9XX57 | 0.601 | 0.481 | -0.654 | 0.995 | 0.242 | 0.344 | 0.205 | 0.0627 |
| G5EEA8 | 0.598 | 0.186 | -0.575 | 0.29 | 0.117 | 0.612 | 0.13 | 0.432 |
| P34654 | 0.595 | -1.18 | 0.0648 | -0.533 | 0.543 | 0.237 | 0.947 | 0.586 |
| O16226 | 0.594 | -1.36 | 0.236 | -1.86 | 0.348 | 0.0429 | 0.706 | 0.00904 |
| Q19324 | 0.593 | 0.913 | 0.0491 | 1.05 | 0.274 | 0.102 | 0.926 | 0.0644 |
| Q22235 | 0.589 | -0.246 | -0.00864 | 0.231 | 0.228 | 0.607 | 0.986 | 0.629 |
| Q23315 | 0.589 | 0.326 | -0.312 | 0.889 | 0.434 | 0.663 | 0.676 | 0.244 |
| Q7Z1Q3 | 0.587 | 0.459 | -0.0829 | -0.0955 | 0.068 | 0.144 | 0.784 | 0.752 |
| Q9UAV5 | 0.582 | 0.074 | -0.178 | -0.0426 | 0.081 | 0.814 | 0.574 | 0.892 |
| Q27249 | 0.581 | -0.329 | 0.0244 | -0.708 | 0.319 | 0.568 | 0.966 | 0.229 |
| G4SLH0 | 0.578 | 0.951 | 1.44 | -0.703 | 0.332 | 0.121 | 0.0251 | 0.242 |
| Q19790 | 0.576 | 0.0812 | -0.013 | -2.63 | 0.49 | 0.922 | 0.987 | 0.00601 |
| Q27511 | 0.572 | 1.28 | 0.319 | 0.807 | 0.133 | 0.0032 | 0.389 | 0.0411 |
| G5EBH7 | 0.572 | -3.87 | 1.2 | -1.3 | 0.607 | 0.00318 | 0.289 | 0.252 |
| Q22370 | 0.571 | -3.73 | -5.02 | -3.29 | 0.571 | 0.00204 | 0.00017 | 0.00497 |
| Q9TYX1 | 0.571 | 0.29 | 0.571 | 0.771 | 0.403 | 0.668 | 0.403 | 0.264 |
| O02640 | 0.568 | -1.21 | -0.274 | -0.824 | 0.117 | 0.00321 | 0.432 | 0.0293 |
| Q9NAB2 | 0.558 | 1.55 | -0.0462 | 1.22 | 0.231 | 0.00381 | 0.919 | 0.0163 |
| P91463 | 0.558 | 1.17 | -0.0621 | 0.154 | 0.31 | 0.0449 | 0.908 | 0.776 |
| Q21151 | 0.558 | 1.34 | 0.586 | -0.277 | 0.325 | 0.0283 | 0.302 | 0.62 |
| H2L0G8 | 0.557 | 0.0321 | 0.922 | -1.08 | 0.285 | 0.95 | 0.0874 | 0.0499 |
| Q17886 | 0.556 | -0.941 | 0.654 | -0.848 | 0.572 | 0.344 | 0.507 | 0.392 |
| O17892 | 0.555 | 0.184 | 0.201 | 1.32 | 0.4 | 0.777 | 0.758 | 0.0584 |
| Q19437 | 0.552 | 0.525 | 0.86 | 0.557 | 0.38 | 0.403 | 0.18 | 0.376 |
| Q966C7 | 0.548 | 0.927 | 0.951 | 1.06 | 0.179 | 0.0316 | 0.0281 | 0.016 |
| Q93831 | 0.545 | -1.06 | -0.683 | -1.67 | 0.419 | 0.126 | 0.314 | 0.0232 |
| P52011 | 0.538 | 0.361 | 0.244 | 0.066 | 0.133 | 0.302 | 0.481 | 0.847 |
| P34697 | 0.537 | -0.867 | -0.0582 | -0.754 | 0.163 | 0.0325 | 0.875 | 0.0579 |
| P34629 | 0.535 | 0.981 | 0.188 | 0.852 | 0.254 | 0.0466 | 0.682 | 0.0787 |
| G5EBP4 | 0.534 | 0.409 | 0.11 | 0.239 | 0.0921 | 0.188 | 0.715 | 0.431 |
| O17921 | 0.527 | 0.606 | -0.191 | 0.859 | 0.119 | 0.077 | 0.557 | 0.0172 |
| H9G2T4 | 0.525 | 0.532 | -0.382 | 0.494 | 0.0931 | 0.0893 | 0.21 | 0.112 |
| Q21624 | 0.522 | -0.146 | -0.798 | -0.269 | 0.516 | 0.855 | 0.326 | 0.736 |
| O17328 | 0.522 | 0.275 | -0.377 | 0.447 | 0.241 | 0.529 | 0.391 | 0.312 |
| O45815 | 0.512 | -0.0731 | 0.0585 | -0.284 | 0.145 | 0.829 | 0.862 | 0.406 |
| Q9UAQ6 | 0.511 | 0.206 | 0.0145 | 0.391 | 0.253 | 0.638 | 0.974 | 0.377 |
| P46822 | 0.499 | 0.515 | -0.0835 | -0.00536 | 0.525 | 0.513 | 0.915 | 0.995 |
| Q23655 | 0.494 | 1.15 | -0.27 | -0.86 | 0.703 | 0.378 | 0.834 | 0.508 |

| | | | | | | | | |
|---|---|---|---|---|---|---|---|---|
| P45971 | 0.492 | 2.26 | 0.456 | 2.2 | 0.593 | 0.0254 | 0.621 | 0.0289 |
| P17140 | 0.48 | -3.31 | -0.173 | -2.76 | 0.319 | 5.59E-06 | 0.715 | 3.89E-05 |
| P34462 | 0.478 | 0.501 | 0.538 | -0.0565 | 0.178 | 0.159 | 0.133 | 0.869 |
| P91303 | 0.475 | -1.21 | 0.313 | -0.424 | 0.29 | 0.014 | 0.48 | 0.343 |
| O01685 | 0.475 | -0.362 | 1.02 | 0.722 | 0.32 | 0.446 | 0.0437 | 0.14 |
| Q0G819 | 0.474 | -0.387 | 0.929 | -0.607 | 0.389 | 0.48 | 0.104 | 0.274 |
| P91134 | 0.472 | -0.442 | -0.471 | 0.88 | 0.655 | 0.676 | 0.656 | 0.409 |
| Q9XVE9 | 0.471 | 1.83 | -0.0519 | 2.05 | 0.339 | 0.00181 | 0.915 | 0.000744 |
| Q22968 | 0.471 | -1.11 | -0.92 | -1.18 | 0.469 | 0.101 | 0.168 | 0.0837 |
| G5EFZ1 | 0.469 | 0.0585 | -1.48 | 0.0655 | 0.499 | 0.932 | 0.0463 | 0.924 |
| O16517 | 0.469 | -0.678 | -1.27 | -0.754 | 0.454 | 0.284 | 0.0551 | 0.235 |
| Q27504 | 0.467 | -1.24 | -0.496 | -1.11 | 0.535 | 0.114 | 0.511 | 0.152 |
| Q9NLD1 | 0.466 | 0.914 | -0.581 | 1.06 | 0.29 | 0.0488 | 0.192 | 0.0255 |
| Q18032 | 0.463 | -1.11 | 0.31 | 0.0313 | 0.367 | 0.0424 | 0.543 | 0.951 |
| D0IMZ5 | 0.462 | 0.244 | 0.651 | 0.185 | 0.143 | 0.427 | 0.0464 | 0.544 |
| O45569 | 0.461 | 1.46 | 0.633 | 1.11 | 0.496 | 0.0443 | 0.354 | 0.115 |
| O18180 | 0.46 | -0.604 | -1.1 | -0.666 | 0.277 | 0.159 | 0.0168 | 0.123 |
| P34385 | 0.457 | 0.289 | 1.55 | 0.127 | 0.577 | 0.723 | 0.0732 | 0.876 |
| Q17490 | 0.456 | 0.05 | -0.664 | -0.0646 | 0.465 | 0.935 | 0.293 | 0.917 |
| P53014 | 0.454 | -0.0691 | -0.507 | -0.136 | 0.341 | 0.883 | 0.29 | 0.772 |
| Q94246 | 0.452 | -0.475 | 0.45 | -0.182 | 0.195 | 0.174 | 0.197 | 0.593 |
| Q86NC1 | 0.45 | 0.866 | 1.39 | 1.11 | 0.387 | 0.108 | 0.0158 | 0.0449 |
| G5EFF8 | 0.442 | -0.462 | 0.475 | 0.329 | 0.396 | 0.376 | 0.363 | 0.525 |
| Q9BKQ7 | 0.442 | 1.12 | 0.973 | 0.419 | 0.397 | 0.0441 | 0.0751 | 0.422 |
| Q17335 | 0.438 | -0.132 | -0.845 | -0.341 | 0.475 | 0.828 | 0.178 | 0.576 |
| Q19782 | 0.436 | -0.418 | -0.413 | -1.87 | 0.666 | 0.679 | 0.683 | 0.0791 |
| A0A5E4LYH5 | 0.435 | -0.848 | 0.754 | -0.0995 | 0.419 | 0.127 | 0.171 | 0.852 |
| Q19052 | 0.434 | -0.192 | -0.0667 | -0.446 | 0.554 | 0.793 | 0.927 | 0.544 |
| P49180 | 0.433 | 1.74 | 0.296 | 1.85 | 0.28 | 0.000506 | 0.455 | 0.000297 |
| Q09983 | 0.432 | 1.24 | 1.87 | 1.25 | 0.582 | 0.128 | 0.0292 | 0.125 |
| Q27245 | 0.431 | 0.36 | -1.35 | 0.407 | 0.212 | 0.293 | 0.00112 | 0.237 |
| P52819 | 0.43 | 1.6 | -0.296 | 1.75 | 0.405 | 0.00669 | 0.563 | 0.00367 |
| O44441 | 0.418 | -1.66 | -0.811 | -2.13 | 0.314 | 0.00102 | 0.0625 | 0.000115 |
| Q9NAH4 | 0.416 | -2.76 | -1.05 | -2.49 | 0.682 | 0.0149 | 0.308 | 0.0251 |
| Q07749 | 0.414 | -1.39 | -1.5 | -0.558 | 0.65 | 0.141 | 0.114 | 0.542 |
| Q18785 | 0.414 | 0.297 | 0.725 | 0.515 | 0.374 | 0.522 | 0.131 | 0.273 |
| D1MN68 | 0.41 | 0.86 | 0.765 | 0.845 | 0.491 | 0.16 | 0.208 | 0.167 |
| Q22866 | 0.407 | -0.232 | 0.489 | -0.432 | 0.35 | 0.59 | 0.264 | 0.322 |
| P46562 | 0.403 | 0.452 | -0.02 | 0.203 | 0.277 | 0.225 | 0.956 | 0.577 |
| Q19289 | 0.402 | -0.151 | -0.416 | -0.378 | 0.314 | 0.7 | 0.298 | 0.342 |
| G5ECG8 | 0.399 | 0.121 | 0.131 | -0.737 | 0.22 | 0.703 | 0.68 | 0.0328 |
| O17214 | 0.396 | -1.6 | -1.73 | -0.88 | 0.479 | 0.0109 | 0.00678 | 0.129 |
| P90916 | 0.394 | -0.266 | 0.435 | -0.148 | 0.356 | 0.529 | 0.309 | 0.726 |
| Q23451 | 0.389 | 1.27 | -1.36 | 1.01 | 0.591 | 0.0952 | 0.0754 | 0.175 |
| Q20588 | 0.383 | -0.298 | -0.22 | -0.818 | 0.52 | 0.615 | 0.71 | 0.181 |
| O62289 | 0.383 | -0.7 | 0.578 | -0.224 | 0.451 | 0.179 | 0.262 | 0.658 |

| | | | | | | | | |
|---|---|---|---|---|---|---|---|---|
| O17586 | 0.379 | 1.16 | -0.274 | 1.88 | 0.55 | 0.0817 | 0.664 | 0.00891 |
| P30642 | 0.373 | 0.92 | -0.231 | 0.433 | 0.365 | 0.0372 | 0.572 | 0.297 |
| Q19842 | 0.372 | -1.2 | -0.257 | -0.697 | 0.383 | 0.0115 | 0.544 | 0.114 |
| Q9N4A7 | 0.37 | 0.691 | -0.685 | 1.33 | 0.484 | 0.201 | 0.204 | 0.0215 |
| Q20239 | 0.37 | 0.493 | -0.0581 | 0.16 | 0.552 | 0.43 | 0.925 | 0.795 |
| Q21929 | 0.363 | -0.379 | -0.636 | -0.666 | 0.556 | 0.539 | 0.308 | 0.287 |
| Q21443 | 0.362 | 0.0837 | -0.131 | -0.137 | 0.292 | 0.804 | 0.697 | 0.684 |
| O02267 | 0.359 | -1.82 | 0.353 | -2.93 | 0.395 | 0.000558 | 0.402 | 5.25E-06 |
| Q94272 | 0.357 | -0.49 | 0.749 | -0.614 | 0.578 | 0.447 | 0.252 | 0.344 |
| A0A2C9C2Y0 | 0.354 | 1.09 | 0.415 | 0.675 | 0.509 | 0.0557 | 0.439 | 0.216 |
| Q9XTU6 | 0.353 | 0.903 | 0.205 | 0.821 | 0.538 | 0.129 | 0.72 | 0.165 |
| P49596 | 0.349 | -1.7 | 0.353 | -1.3 | 0.601 | 0.0212 | 0.597 | 0.0662 |
| Q95005 | 0.345 | -1.51 | -1.43 | -2.17 | 0.595 | 0.0323 | 0.0404 | 0.00424 |
| A0A061AD47 | 0.345 | -2.91 | 0.806 | -3.48 | 0.415 | 5.98E-06 | 0.07 | 7.88E-07 |
| P54216 | 0.345 | 0.0307 | -0.307 | -0.186 | 0.279 | 0.921 | 0.334 | 0.554 |
| P10299 | 0.341 | -0.0437 | -1.32 | -0.317 | 0.485 | 0.928 | 0.015 | 0.517 |
| P98080 | 0.334 | 0.411 | -0.845 | 0.272 | 0.518 | 0.429 | 0.116 | 0.598 |
| Q965S0 | 0.333 | -0.693 | 0.513 | -1.12 | 0.415 | 0.103 | 0.217 | 0.0138 |
| Q22018 | 0.332 | -1.23 | -0.536 | -0.942 | 0.51 | 0.0259 | 0.294 | 0.0759 |
| A0A4V6M4Y3 | 0.332 | -1.19 | 0.354 | -1.65 | 0.626 | 0.0953 | 0.603 | 0.0266 |
| Q93244 | 0.329 | -0.484 | -0.0311 | -0.308 | 0.462 | 0.285 | 0.944 | 0.49 |
| P55955 | 0.322 | -0.103 | 0.264 | -0.166 | 0.391 | 0.781 | 0.479 | 0.654 |
| O01542 | 0.316 | 1.17 | -0.0454 | -0.181 | 0.431 | 0.00952 | 0.909 | 0.648 |
| Q23248 | 0.316 | 1.58 | 0.202 | 2.84 | 0.748 | 0.124 | 0.837 | 0.0107 |
| Q20476 | 0.315 | 1.01 | 0.0191 | 0.579 | 0.718 | 0.258 | 0.982 | 0.509 |
| Q9NF11 | 0.315 | 0.693 | -0.441 | 0.244 | 0.569 | 0.22 | 0.427 | 0.658 |
| Q19967 | 0.313 | 2.06 | 0.102 | -0.119 | 0.771 | 0.0714 | 0.925 | 0.912 |
| K8ERV5 | 0.312 | -1.32 | -0.0964 | -0.458 | 0.616 | 0.047 | 0.876 | 0.463 |
| Q9NES7 | 0.311 | -1.49 | -1.98 | -1.42 | 0.695 | 0.0757 | 0.0231 | 0.0888 |
| P91917 | 0.307 | 1.17 | -1.08 | 1.25 | 0.546 | 0.0344 | 0.0472 | 0.0248 |
| O02268 | 0.307 | 0.866 | 0.72 | 0.151 | 0.582 | 0.135 | 0.208 | 0.785 |
| P92005 | 0.307 | 0.404 | 1.35 | 0.271 | 0.793 | 0.73 | 0.26 | 0.817 |
| O44480 | 0.306 | 1.89 | 0.153 | 2.21 | 0.542 | 0.0018 | 0.759 | 0.000518 |
| Q7JLY3 | 0.299 | -1.05 | 0.833 | -1.83 | 0.529 | 0.04 | 0.0945 | 0.00148 |
| Q27894 | 0.299 | 0.961 | -0.0298 | 0.8 | 0.44 | 0.0231 | 0.938 | 0.0519 |
| P35129 | 0.296 | 0.591 | -1.12 | -0.589 | 0.707 | 0.456 | 0.168 | 0.457 |
| Q18240 | 0.296 | 0.357 | -0.613 | 1.13 | 0.712 | 0.658 | 0.45 | 0.173 |
| Q9XVI9 | 0.296 | 0.417 | 0.197 | 0.369 | 0.718 | 0.611 | 0.81 | 0.653 |
| P53489 | 0.294 | 1.49 | -0.0989 | 2.4 | 0.652 | 0.0352 | 0.879 | 0.00216 |
| Q20728 | 0.294 | -0.341 | -0.325 | -0.385 | 0.757 | 0.719 | 0.732 | 0.685 |
| P34382 | 0.291 | -0.289 | -1.21 | -0.929 | 0.621 | 0.624 | 0.0546 | 0.13 |
| P91427 | 0.281 | 0.0383 | -0.596 | 0.183 | 0.437 | 0.915 | 0.113 | 0.611 |
| Q17698 | 0.277 | 0.0801 | -1.14 | 0.442 | 0.577 | 0.871 | 0.0338 | 0.378 |
| Q18359 | 0.273 | -0.861 | -1.08 | -0.32 | 0.534 | 0.0637 | 0.0245 | 0.467 |

| | | | | | | | | |
|---|---|---|---|---|---|---|---|---|
| Q93615 | 0.271 | 0.452 | -0.549 | 0.0539 | 0.47 | 0.235 | 0.154 | 0.885 |
| P27604 | 0.27 | 1.08 | -0.541 | 1.17 | 0.479 | 0.0112 | 0.167 | 0.00725 |
| O44512 | 0.266 | -0.654 | -1.4 | -1.59 | 0.729 | 0.399 | 0.0849 | 0.0527 |
| Q21265 | 0.265 | -0.914 | 0.0785 | -0.29 | 0.496 | 0.0307 | 0.839 | 0.457 |
| Q3LFN1 | 0.264 | 1.1 | -0.498 | 1.02 | 0.518 | 0.0157 | 0.231 | 0.0231 |
| Q18020 | 0.26 | -0.452 | -0.459 | -0.991 | 0.662 | 0.45 | 0.442 | 0.11 |
| Q9N4X8 | 0.26 | 0.264 | -0.18 | 0.703 | 0.631 | 0.626 | 0.74 | 0.206 |
| P52554 | 0.258 | 0.104 | -0.0663 | -2.17 | 0.602 | 0.833 | 0.893 | 0.000518 |
| Q19264 | 0.253 | 0.602 | -0.277 | 0.686 | 0.533 | 0.151 | 0.495 | 0.105 |
| Q9N5M2 | 0.253 | -0.0774 | 0.133 | -0.269 | 0.745 | 0.921 | 0.864 | 0.73 |
| Q9N4L8 | 0.25 | -2.17 | -0.281 | -2.66 | 0.738 | 0.0105 | 0.707 | 0.00285 |
| Q22099 | 0.249 | 0.941 | 0.912 | 0.881 | 0.481 | 0.0162 | 0.0191 | 0.0227 |
| Q21276 | 0.246 | 1.21 | 1.59 | 0.833 | 0.817 | 0.267 | 0.151 | 0.439 |
| Q9N456 | 0.244 | -1.28 | 0.661 | -1.51 | 0.721 | 0.0778 | 0.341 | 0.0416 |
| Q94271 | 0.242 | 1.54 | -0.714 | 0.794 | 0.74 | 0.0492 | 0.336 | 0.286 |
| Q86DC6 | 0.24 | -0.048 | -0.823 | -0.464 | 0.629 | 0.923 | 0.112 | 0.355 |
| Q95008 | 0.237 | 0.376 | -0.915 | -0.641 | 0.759 | 0.627 | 0.247 | 0.411 |
| Q22347 | 0.235 | -1.27 | -0.998 | -0.53 | 0.599 | 0.0113 | 0.0383 | 0.244 |
| Q17770 | 0.232 | -0.151 | -0.368 | 0.077 | 0.561 | 0.703 | 0.361 | 0.846 |
| Q9N5V3 | 0.229 | 2.66 | -0.975 | 2.03 | 0.695 | 0.000388 | 0.111 | 0.00327 |
| Q22850 | 0.228 | -2.39 | 1.07 | -2.01 | 0.8 | 0.0175 | 0.245 | 0.0393 |
| Q19579 | 0.226 | 0.851 | 0.17 | 0.414 | 0.786 | 0.315 | 0.838 | 0.62 |
| Q09544 | 0.225 | -1.23 | -0.469 | -0.812 | 0.566 | 0.00649 | 0.241 | 0.0526 |
| P27420 | 0.224 | -0.0487 | -0.944 | 0.38 | 0.576 | 0.903 | 0.0303 | 0.348 |
| Q9XVT2 | 0.219 | -0.364 | 0.243 | -0.737 | 0.733 | 0.571 | 0.705 | 0.261 |
| Q23312 | 0.218 | 1.28 | -0.407 | 1.52 | 0.651 | 0.0165 | 0.401 | 0.00617 |
| Q19973 | 0.218 | -0.26 | -1.31 | 0.839 | 0.739 | 0.691 | 0.0611 | 0.212 |
| A0A4V0IJ<br>W4 | 0.216 | -0.188 | 0.514 | -1.31 | 0.792 | 0.819 | 0.533 | 0.125 |
| P91127 | 0.214 | -0.102 | 0.482 | -0.776 | 0.644 | 0.825 | 0.305 | 0.109 |
| G5EEG4 | 0.211 | -1.43 | -0.478 | -1.47 | 0.665 | 0.00954 | 0.334 | 0.00835 |
| P54811 | 0.204 | 0.454 | 0.292 | 0.42 | 0.498 | 0.145 | 0.336 | 0.175 |
| Q23621 | 0.204 | -0.52 | -0.698 | -0.322 | 0.507 | 0.106 | 0.0358 | 0.302 |
| G5ED31 | 0.2 | -3.03 | -2.24 | -2.17 | 0.788 | 0.000995 | 0.00829 | 0.01 |
| G5EFP2 | 0.199 | -1.41 | -0.633 | -0.76 | 0.799 | 0.0872 | 0.423 | 0.338 |
| P90904 | 0.199 | 0.781 | 1.66 | 0.369 | 0.747 | 0.218 | 0.0161 | 0.551 |
| H2L2J0 | 0.196 | -0.917 | 0.599 | -1.45 | 0.711 | 0.1 | 0.269 | 0.0146 |
| Q95Y90 | 0.195 | 1.86 | 0.866 | 1.8 | 0.586 | 0.000115 | 0.0268 | 0.000158 |
| Q9GRZ9 | 0.194 | 0.0938 | -0.0759 | -1.27 | 0.761 | 0.883 | 0.905 | 0.0637 |
| H2KYI5 | 0.193 | -0.19 | 0.773 | 0.838 | 0.837 | 0.839 | 0.414 | 0.377 |
| Q23280 | 0.192 | 0.894 | -1.42 | 1.14 | 0.693 | 0.0815 | 0.00983 | 0.032 |
| Q09476 | 0.189 | -2.25 | 0.278 | -2.55 | 0.633 | 4.87E-05 | 0.484 | 1.31E-05 |
| Q27527 | 0.188 | -0.484 | -0.229 | -0.46 | 0.533 | 0.122 | 0.45 | 0.14 |
| O62102 | 0.187 | 0.715 | -0.812 | 0.452 | 0.665 | 0.113 | 0.0757 | 0.303 |
| O45924 | 0.184 | -1.06 | -2.06 | -1.89 | 0.825 | 0.215 | 0.025 | 0.0371 |
| Q7K6X4 | 0.181 | -0.951 | 1.08 | -1.39 | 0.721 | 0.0754 | 0.0469 | 0.0138 |
| Q9XXE2 | 0.18 | -0.207 | -0.346 | -0.093 | 0.685 | 0.642 | 0.441 | 0.834 |

| | | | | | | | | |
|---|---|---|---|---|---|---|---|---|
| Q20660 | 0.176 | -1.89 | 0.397 | -1.6 | 0.654 | 0.00023 | 0.319 | 0.00098 |
| G5EFQ9 | 0.176 | 0.129 | 0.393 | 0.658 | 0.688 | 0.768 | 0.375 | 0.148 |
| Q22922 | 0.175 | -0.151 | -0.816 | 0.491 | 0.788 | 0.817 | 0.222 | 0.454 |
| Q5CCJ2 | 0.173 | 0.551 | -0.34 | -0.0666 | 0.73 | 0.28 | 0.499 | 0.894 |
| G5EEL2 | 0.171 | 0.953 | 1.08 | -0.948 | 0.742 | 0.0831 | 0.0529 | 0.0845 |
| O62431 | 0.169 | 2.35 | 0.522 | 1.75 | 0.863 | 0.0285 | 0.595 | 0.0897 |
| Q17489 | 0.165 | 0.117 | 0.874 | 0.169 | 0.682 | 0.77 | 0.0443 | 0.676 |
| P52652 | 0.159 | 0.656 | 0.0652 | -0.0258 | 0.822 | 0.359 | 0.926 | 0.971 |
| Q93716 | 0.157 | 0.847 | 0.469 | 0.238 | 0.752 | 0.104 | 0.351 | 0.633 |
| Q93934 | 0.154 | 1.49 | -0.35 | 1.23 | 0.686 | 0.00138 | 0.364 | 0.00529 |
| Q9N4F0 | 0.15 | 0.335 | 0.592 | 0.206 | 0.739 | 0.461 | 0.202 | 0.648 |
| Q9BKP8 | 0.149 | 1.16 | 0.0685 | 1.4 | 0.769 | 0.0348 | 0.892 | 0.0142 |
| Q93637 | 0.147 | -0.217 | 0.75 | -0.28 | 0.699 | 0.571 | 0.0646 | 0.466 |
| Q18136 | 0.146 | 1.16 | 0.456 | 0.476 | 0.705 | 0.00829 | 0.247 | 0.228 |
| P34340 | 0.145 | 0.692 | 1.76 | -0.592 | 0.79 | 0.216 | 0.0054 | 0.285 |
| O45550 | 0.143 | 0.675 | 0.179 | 0.39 | 0.768 | 0.18 | 0.713 | 0.428 |
| O16264 | 0.137 | -1.48 | -0.922 | -1.49 | 0.886 | 0.137 | 0.343 | 0.136 |
| O17919 | 0.134 | 0.907 | 0.159 | 0.606 | 0.723 | 0.0288 | 0.675 | 0.126 |
| P53439 | 0.13 | 0.772 | 0.254 | 0.874 | 0.797 | 0.143 | 0.618 | 0.101 |
| Q21230 | 0.128 | 1.66 | 1.26 | 0.998 | 0.813 | 0.00765 | 0.0334 | 0.0821 |
| Q9TXI3 | 0.128 | -0.813 | -0.537 | -1.01 | 0.853 | 0.25 | 0.441 | 0.159 |
| Q9GUM7 | 0.127 | -0.293 | -0.693 | -1.56 | 0.776 | 0.514 | 0.136 | 0.00313 |
| Q19087 | 0.126 | 0.606 | -0.123 | 0.39 | 0.713 | 0.0918 | 0.719 | 0.263 |
| P91477 | 0.123 | -0.0476 | -2.01 | 0.029 | 0.865 | 0.948 | 0.0139 | 0.968 |
| P29691 | 0.121 | 1.12 | 0.0569 | 1.3 | 0.703 | 0.00291 | 0.857 | 0.000933 |
| O16458 | 0.118 | 1.46 | -0.218 | 0.803 | 0.837 | 0.0215 | 0.705 | 0.177 |
| Q19102 | 0.114 | -1.14 | -0.857 | -0.5 | 0.718 | 0.00246 | 0.0149 | 0.128 |
| Q9XX15 | 0.107 | 0.955 | 1.21 | 0.234 | 0.857 | 0.124 | 0.057 | 0.694 |
| P34453 | 0.105 | 0.292 | -0.14 | -0.822 | 0.882 | 0.682 | 0.844 | 0.259 |
| G5EDN6 | 0.104 | 0.558 | 0.729 | -0.0624 | 0.861 | 0.356 | 0.233 | 0.916 |
| O01802 | 0.098 | 1.57 | 0.0767 | 1.27 | 0.832 | 0.00396 | 0.868 | 0.0143 |
| Q18786 | 0.095 | -3.06 | -0.828 | -2.59 | 0.91 | 0.00237 | 0.332 | 0.0074 |
| Q9N4L9 | 0.0892 | -0.048 | -0.45 | -1.71 | 0.91 | 0.951 | 0.571 | 0.045 |
| Q18709 | 0.0876 | -0.533 | 0.0121 | -0.385 | 0.858 | 0.285 | 0.98 | 0.435 |
| P19826 | 0.0858 | -1.14 | 0.465 | 0.439 | 0.918 | 0.183 | 0.577 | 0.598 |
| Q19724 | 0.0853 | -1.02 | -0.477 | -1.09 | 0.924 | 0.265 | 0.597 | 0.238 |
| Q18685 | 0.0835 | -1.3 | 0.705 | -1.04 | 0.899 | 0.0651 | 0.295 | 0.131 |
| E3W741 | 0.0807 | -3.81 | 0.95 | -4.03 | 0.938 | 0.00224 | 0.367 | 0.00147 |
| Q19063 | 0.0785 | -1.51 | 1.02 | -1.17 | 0.919 | 0.0662 | 0.202 | 0.147 |
| O01593 | 0.0775 | -0.0814 | -0.346 | -0.41 | 0.934 | 0.931 | 0.713 | 0.663 |
| Q19775 | 0.0764 | -1.86 | -1.69 | -1.64 | 0.94 | 0.0826 | 0.112 | 0.122 |
| P91177 | 0.0751 | -0.704 | 0.437 | 0.568 | 0.9 | 0.248 | 0.466 | 0.348 |
| Q9U1W8 | 0.0694 | 1.65 | 0.971 | 0.848 | 0.895 | 0.00663 | 0.0822 | 0.124 |
| Q9N537 | 0.0643 | -1.03 | -0.331 | 0.145 | 0.937 | 0.216 | 0.684 | 0.858 |
| O61217 | 0.0605 | -0.204 | 0.377 | 0.103 | 0.865 | 0.571 | 0.301 | 0.773 |
| Q1XFY4 | 0.0531 | 0.382 | 0.23 | 0.156 | 0.914 | 0.442 | 0.64 | 0.751 |
| Q95Y72 | 0.0475 | 0.381 | 0.509 | 0.584 | 0.91 | 0.37 | 0.236 | 0.177 |

| | | | | | | | | |
|---|---|---|---|---|---|---|---|---|
| Q17993 | 0.0439 | 0.388 | -0.153 | 0.758 | 0.939 | 0.504 | 0.791 | 0.201 |
| A0A5E4LWV7 | 0.0415 | -0.882 | 0.0652 | -1.7 | 0.965 | 0.363 | 0.946 | 0.0919 |
| O76371 | 0.0411 | -1.71 | -0.0638 | -0.438 | 0.935 | 0.00391 | 0.899 | 0.392 |
| O16305 | 0.0404 | 0.384 | -0.305 | 0.34 | 0.934 | 0.437 | 0.535 | 0.49 |
| Q9N3B9 | 0.0391 | 0.279 | -0.493 | -0.0342 | 0.945 | 0.622 | 0.388 | 0.952 |
| Q95X44 | 0.0349 | -0.348 | -0.107 | -0.394 | 0.921 | 0.333 | 0.762 | 0.275 |
| A0A486WVD1 | 0.0341 | -3.3 | -0.801 | -3.45 | 0.955 | 8.42E-05 | 0.204 | 5.37E-05 |
| Q17348 | 0.0312 | 1 | -0.0865 | 0.376 | 0.921 | 0.00579 | 0.783 | 0.242 |
| Q9BI71 | 0.0291 | -1.37 | 0.607 | -1.9 | 0.95 | 0.00907 | 0.201 | 0.000909 |
| O44895 | 0.0258 | -2.54 | 0.253 | -3.17 | 0.968 | 0.00138 | 0.696 | 0.000207 |
| Q17473 | 0.025 | -0.241 | -0.576 | -0.109 | 0.939 | 0.463 | 0.0933 | 0.738 |
| Q04908 | 0.0248 | 0.854 | 0.665 | 0.188 | 0.951 | 0.05 | 0.117 | 0.644 |
| P34346 | 0.023 | -2.26 | -0.0353 | -0.452 | 0.977 | 0.0124 | 0.965 | 0.575 |
| O17626 | 0.023 | 0.341 | -0.000426 | 0.376 | 0.945 | 0.313 | 0.999 | 0.268 |
| Q9GRY9 | 0.0213 | -2.1 | 0.509 | -0.475 | 0.964 | 0.000527 | 0.295 | 0.328 |
| P90961 | 0.0187 | -0.465 | 0.128 | 0.275 | 0.959 | 0.211 | 0.722 | 0.45 |
| P49181 | 0.0174 | 1.25 | 0.205 | 1.41 | 0.967 | 0.00974 | 0.63 | 0.00454 |
| G5EF37 | 0.0164 | -0.12 | -0.49 | 0.0555 | 0.976 | 0.825 | 0.373 | 0.919 |
| Q93695 | 0.0153 | 0.598 | -0.0951 | 0.129 | 0.973 | 0.204 | 0.835 | 0.778 |
| Q21338 | 0.0148 | 1.38 | 1.18 | 0.519 | 0.982 | 0.0525 | 0.0917 | 0.437 |
| P54412 | 0.0147 | 1.28 | 0.0612 | 1.08 | 0.968 | 0.00311 | 0.867 | 0.00944 |
| O44995 | 0.0144 | 0.0361 | -0.166 | -0.562 | 0.984 | 0.959 | 0.814 | 0.429 |
| Q9N4Y8 | 0.0119 | -0.284 | 0.773 | 0.701 | 0.989 | 0.745 | 0.381 | 0.426 |
| Q21024 | 0.00821 | 0.751 | -0.661 | 0.594 | 0.986 | 0.133 | 0.182 | 0.228 |
| O45562 | 0.00661 | -2.75 | 0.491 | -2.78 | 0.992 | 0.00115 | 0.478 | 0.00105 |
| Q9TZC4 | 0.00253 | 0.908 | -0.9 | 1.74 | 0.997 | 0.197 | 0.201 | 0.0215 |
| Q9TZS5 | 0.00179 | 0.621 | -0.955 | 1.2 | 0.998 | 0.363 | 0.17 | 0.0907 |
| Q93778 | 0.00121 | 0.739 | 0.235 | 2.82 | 0.999 | 0.333 | 0.754 | 0.00188 |
| Q9U2X0 | -0.00181 | 1.54 | -0.331 | 0.463 | 0.996 | 0.00113 | 0.395 | 0.239 |
| Q17430 | -0.00191 | 0.484 | 0.239 | 0.327 | 0.997 | 0.403 | 0.677 | 0.57 |
| P41932 | -0.00473 | 0.448 | -0.143 | 0.228 | 0.988 | 0.162 | 0.645 | 0.465 |
| P51404 | -0.00525 | 1.34 | -0.185 | 1.36 | 0.989 | 0.00333 | 0.632 | 0.00296 |
| Q9XUW5 | -0.00934 | 1.12 | 0.524 | 1.12 | 0.989 | 0.119 | 0.451 | 0.118 |
| Q965Q1 | -0.0106 | -0.608 | -0.0834 | 0.162 | 0.981 | 0.197 | 0.855 | 0.724 |
| Q09289 | -0.0109 | -1.48 | 0.329 | -1.15 | 0.986 | 0.0305 | 0.6 | 0.0813 |
| P54813 | -0.0183 | 0.796 | 0.585 | 2 | 0.97 | 0.113 | 0.234 | 0.000841 |
| Q23670 | -0.022 | 0.724 | 1.26 | 0.795 | 0.982 | 0.456 | 0.203 | 0.414 |
| O45012 | -0.0242 | 0.906 | 1.28 | 0.735 | 0.95 | 0.0304 | 0.00434 | 0.0707 |
| Q95Y73 | -0.0306 | 4.27 | 1.05 | 3.4 | 0.973 | 0.000244 | 0.247 | 0.00162 |
| A0A061AKY5 | -0.0341 | -1.49 | 1.27 | -0.183 | 0.973 | 0.154 | 0.218 | 0.856 |
| Q95Y92 | -0.0419 | -0.03 | 1.87 | -0.232 | 0.95 | 0.964 | 0.013 | 0.729 |
| Q9U2F6 | -0.0433 | 0.0577 | 1.06 | 0.417 | 0.936 | 0.916 | 0.0667 | 0.448 |
| O01541 | -0.0456 | 1.14 | -0.0793 | 0.0985 | 0.943 | 0.0941 | 0.902 | 0.878 |
| Q23120 | -0.0461 | 0.179 | -0.00884 | 0.172 | 0.898 | 0.62 | 0.98 | 0.634 |

| | | | | | | | | |
|---|---|---|---|---|---|---|---|---|
| P90778 | -0.048 | -1.09 | -0.125 | -1.18 | 0.963 | 0.305 | 0.904 | 0.266 |
| Q95YF3 | -0.0489 | 1.49 | 0.849 | 1.53 | 0.918 | 0.00636 | 0.0891 | 0.00542 |
| P34539 | -0.0494 | -0.291 | -1.2 | -0.919 | 0.938 | 0.649 | 0.0768 | 0.165 |
| P11141 | -0.0531 | -0.435 | -0.759 | 0.0501 | 0.885 | 0.246 | 0.0532 | 0.891 |
| Q95XR0 | -0.0547 | -0.529 | -0.292 | 0.979 | 0.953 | 0.572 | 0.754 | 0.303 |
| G5ECG0 | -0.0603 | 0.0897 | -0.0839 | -0.345 | 0.921 | 0.882 | 0.89 | 0.572 |
| P34328 | -0.0648 | -0.704 | 0.0111 | -1.17 | 0.887 | 0.139 | 0.981 | 0.021 |
| O02155 | -0.0698 | 0.479 | 0.905 | -2.11 | 0.938 | 0.594 | 0.321 | 0.0309 |
| P91914 | -0.071 | 1.38 | -0.722 | 1.7 | 0.911 | 0.0444 | 0.266 | 0.0167 |
| O62122 | -0.0731 | 0.109 | -1.04 | -0.337 | 0.921 | 0.882 | 0.17 | 0.648 |
| Q9U302 | -0.0818 | -0.00967 | 0.416 | 0.0747 | 0.837 | 0.981 | 0.304 | 0.851 |
| H2KYV3 | -0.0871 | -0.312 | 1.33 | -0.789 | 0.883 | 0.601 | 0.0386 | 0.198 |
| Q17763 | -0.0872 | -1.78 | -1.23 | -1.49 | 0.828 | 0.000501 | 0.00746 | 0.00208 |
| Q9TXU7 | -0.0885 | -0.509 | 0.122 | -0.569 | 0.809 | 0.179 | 0.739 | 0.136 |
| Q18124 | -0.0912 | -0.356 | -0.76 | 0.332 | 0.94 | 0.77 | 0.535 | 0.785 |
| P46548 | -0.0913 | 1.36 | -1.14 | 0.583 | 0.881 | 0.0394 | 0.0792 | 0.347 |
| Q86LS4 | -0.0923 | -0.0392 | 0.469 | 0.467 | 0.862 | 0.941 | 0.383 | 0.384 |
| P49041 | -0.0968 | 1.07 | -0.0629 | 0.742 | 0.754 | 0.00331 | 0.839 | 0.0284 |
| G5EET8 | -0.0981 | -0.239 | -0.0986 | -0.0898 | 0.806 | 0.553 | 0.805 | 0.822 |
| Q19328 | -0.102 | 1.34 | 0.349 | 1.08 | 0.758 | 0.00106 | 0.302 | 0.00526 |
| Q95QV8 | -0.104 | 0.166 | -0.0352 | 0.426 | 0.782 | 0.661 | 0.926 | 0.27 |
| O17861 | -0.106 | -2.07 | -2.11 | -1.66 | 0.928 | 0.0935 | 0.0881 | 0.17 |
| Q93791 | -0.108 | -1.41 | 0.645 | -1.86 | 0.875 | 0.056 | 0.355 | 0.0154 |
| G5EFL5 | -0.111 | -1.35 | 0.601 | -1.04 | 0.829 | 0.0179 | 0.253 | 0.0589 |
| Q18026 | -0.111 | 0.598 | -0.565 | 0.263 | 0.833 | 0.264 | 0.29 | 0.616 |
| G4SI07 | -0.111 | 0.498 | 0.277 | 0.498 | 0.695 | 0.095 | 0.337 | 0.095 |
| Q9XWK2 | -0.111 | 1.19 | 0.278 | 1.01 | 0.835 | 0.0393 | 0.603 | 0.0744 |
| Q94230 | -0.112 | 0.151 | -0.286 | 0.287 | 0.738 | 0.653 | 0.4 | 0.399 |
| Q09610 | -0.114 | -1.39 | -2.14 | 0.57 | 0.934 | 0.324 | 0.138 | 0.681 |
| P43509 | -0.117 | 1.27 | 0.555 | 1.45 | 0.88 | 0.116 | 0.475 | 0.075 |
| O62106 | -0.127 | 0.546 | -0.888 | 0.55 | 0.791 | 0.265 | 0.0798 | 0.261 |
| O45346 | -0.128 | -2.58 | 0.501 | -3.11 | 0.833 | 0.00072 | 0.415 | 0.000137 |
| Q21826 | -0.132 | 1.06 | 0.569 | 0.444 | 0.838 | 0.119 | 0.387 | 0.497 |
| G5EFV4 | -0.135 | -0.126 | -0.0459 | 0.114 | 0.754 | 0.771 | 0.915 | 0.79 |
| Q21750 | -0.137 | 0.62 | -0.639 | 0.892 | 0.749 | 0.162 | 0.151 | 0.0523 |
| Q9TXI4 | -0.139 | -2.5 | -0.545 | -1.79 | 0.704 | 7.46E-06 | 0.152 | 0.000218 |
| Q11067 | -0.141 | -1.52 | -0.239 | 0.0153 | 0.748 | 0.0033 | 0.585 | 0.972 |
| P04970 | -0.143 | 1.13 | 0.183 | 1.01 | 0.847 | 0.144 | 0.804 | 0.185 |
| Q9XVP0 | -0.144 | 1.3 | -0.0431 | 1.21 | 0.747 | 0.0107 | 0.923 | 0.0157 |
| Q7JKP6 | -0.147 | -1.11 | 0.597 | -0.61 | 0.827 | 0.115 | 0.382 | 0.372 |
| G3MU72 | -0.151 | 0.23 | 0.0489 | 0.864 | 0.734 | 0.605 | 0.912 | 0.0672 |
| Q18066 | -0.153 | -0.473 | 0.362 | -0.771 | 0.624 | 0.145 | 0.256 | 0.0247 |
| P19626 | -0.155 | -1.08 | -0.304 | -1.13 | 0.681 | 0.0113 | 0.425 | 0.00855 |
| Q20448 | -0.156 | 0.109 | 1.71 | -0.796 | 0.825 | 0.878 | 0.0271 | 0.27 |
| Q17561 | -0.157 | -2.12 | 0.183 | -2.3 | 0.758 | 0.000861 | 0.72 | 0.000436 |
| P91240 | -0.159 | -0.963 | 2.16 | -1.96 | 0.9 | 0.451 | 0.104 | 0.136 |
| B1V8A0 | -0.164 | -0.605 | 2.56 | -1.05 | 0.893 | 0.622 | 0.0516 | 0.396 |

| | | | | | | | | |
|---|---|---|---|---|---|---|---|---|
| Q9XUN9 | -0.165 | -1.9 | -0.239 | -1.7 | 0.713 | 0.000714 | 0.594 | 0.00172 |
| P27798 | -0.166 | -0.699 | -0.596 | -0.193 | 0.74 | 0.177 | 0.245 | 0.699 |
| P09588 | -0.169 | 0.486 | -0.341 | 0.548 | 0.672 | 0.235 | 0.399 | 0.183 |
| Q19128 | -0.171 | 0.509 | 0.165 | 0.00915 | 0.76 | 0.369 | 0.768 | 0.987 |
| P48152 | -0.172 | 1.34 | 0.0716 | 1.31 | 0.67 | 0.00457 | 0.859 | 0.00536 |
| P52015 | -0.173 | 0.331 | 0.205 | 0.316 | 0.566 | 0.279 | 0.498 | 0.301 |
| P52709 | -0.174 | 0.279 | 0.0516 | 0.465 | 0.623 | 0.433 | 0.883 | 0.199 |
| Q10121 | -0.175 | -1.22 | -0.506 | -0.53 | 0.828 | 0.144 | 0.532 | 0.513 |
| Q22752 | -0.178 | 0.87 | -0.235 | 0.288 | 0.746 | 0.129 | 0.67 | 0.602 |
| Q23307 | -0.195 | -0.334 | 0.699 | -0.829 | 0.608 | 0.383 | 0.0808 | 0.0426 |
| G5EFG4 | -0.197 | 1.45 | 0.288 | 0.868 | 0.735 | 0.0235 | 0.621 | 0.15 |
| P34662 | -0.198 | 1.62 | -0.388 | 1.48 | 0.6 | 0.000635 | 0.31 | 0.0013 |
| B0M0L8 | -0.202 | -2.36 | -1.08 | -2.34 | 0.766 | 0.00323 | 0.125 | 0.00344 |
| Q95PX7 | -0.202 | 0.556 | 0.377 | -0.17 | 0.813 | 0.518 | 0.66 | 0.843 |
| G5EEF0 | -0.203 | -0.154 | 0.508 | -0.183 | 0.578 | 0.672 | 0.176 | 0.616 |
| Q9TZ68 | -0.209 | -2.83 | -1.33 | -3.31 | 0.79 | 0.00261 | 0.107 | 0.000766 |
| Q21926 | -0.213 | 1.56 | -0.00806 | 1.4 | 0.733 | 0.0233 | 0.99 | 0.0381 |
| Q21614 | -0.218 | 2.22 | -0.448 | 0.889 | 0.822 | 0.0346 | 0.644 | 0.364 |
| Q18100 | -0.219 | -0.617 | 0.99 | -0.711 | 0.645 | 0.207 | 0.0524 | 0.15 |
| Q94269 | -0.219 | 0.0787 | -0.857 | 0.323 | 0.557 | 0.832 | 0.0339 | 0.39 |
| Q20203 | -0.222 | -2.82 | -2.14 | -3.39 | 0.812 | 0.00823 | 0.0347 | 0.00239 |
| A0A8S4Q9F9 | -0.226 | 0.324 | 0.894 | -0.0144 | 0.689 | 0.568 | 0.129 | 0.98 |
| Q17880 | -0.226 | 0.754 | -0.382 | 0.676 | 0.754 | 0.305 | 0.598 | 0.356 |
| Q93408 | -0.229 | -1.16 | 0.0252 | -1.67 | 0.755 | 0.129 | 0.972 | 0.036 |
| P34255 | -0.232 | -3.13 | -1.34 | -3.09 | 0.788 | 0.00238 | 0.134 | 0.00264 |
| Q9XVB4 | -0.233 | 0.553 | 0.831 | -0.239 | 0.684 | 0.341 | 0.161 | 0.676 |
| P52013 | -0.235 | -0.775 | -0.524 | -0.481 | 0.506 | 0.0412 | 0.15 | 0.184 |
| Q86S66 | -0.235 | 0.323 | -0.0152 | 0.0937 | 0.566 | 0.432 | 0.97 | 0.818 |
| O17345 | -0.237 | 0.753 | 0.644 | -0.105 | 0.598 | 0.109 | 0.166 | 0.815 |
| Q21021 | -0.238 | 0.425 | -0.138 | 0.609 | 0.539 | 0.279 | 0.719 | 0.129 |
| G5EBF3 | -0.247 | -0.986 | -0.0508 | -0.329 | 0.661 | 0.0958 | 0.928 | 0.56 |
| P46550 | -0.25 | -0.00541 | 0.00368 | 0.735 | 0.665 | 0.993 | 0.995 | 0.216 |
| Q95ZW2 | -0.253 | -0.269 | -1.5 | -0.42 | 0.681 | 0.663 | 0.026 | 0.497 |
| Q9BL61 | -0.254 | 0.632 | 0.312 | 0.0598 | 0.56 | 0.16 | 0.475 | 0.89 |
| O01692 | -0.258 | 0.666 | 0.398 | 0.939 | 0.461 | 0.0713 | 0.262 | 0.0157 |
| P48154 | -0.258 | 1.44 | -0.138 | 1.27 | 0.45 | 0.000717 | 0.683 | 0.00193 |
| P46769 | -0.262 | 1.11 | 0.201 | 1.14 | 0.495 | 0.0105 | 0.601 | 0.00902 |
| Q21746 | -0.262 | 0.192 | -0.352 | 0.12 | 0.429 | 0.561 | 0.292 | 0.716 |
| Q6A573 | -0.262 | 0.962 | 0.537 | 0.552 | 0.702 | 0.174 | 0.436 | 0.425 |
| Q9N4I4 | -0.263 | 1.03 | -0.183 | 1.2 | 0.509 | 0.0187 | 0.645 | 0.00807 |
| P30629 | -0.266 | -1.66 | 1.28 | -0.819 | 0.688 | 0.0229 | 0.0686 | 0.227 |
| Q05036 | -0.269 | 0.787 | 0.719 | -0.585 | 0.729 | 0.32 | 0.362 | 0.456 |
| Q21215 | -0.27 | 0.304 | -0.945 | 0.725 | 0.482 | 0.429 | 0.0243 | 0.073 |
| Q9N3X2 | -0.27 | 0.944 | -0.0386 | 0.827 | 0.423 | 0.0122 | 0.908 | 0.0245 |
| Q95Y93 | -0.271 | -1.32 | 0.74 | -0.359 | 0.486 | 0.00373 | 0.0708 | 0.358 |
| Q18231 | -0.293 | 0.522 | 0.218 | 0.676 | 0.514 | 0.252 | 0.626 | 0.145 |

| | | | | | | | | |
|---|---|---|---|---|---|---|---|---|
| Q18164 | -0.294 | 0.3 | 1.2 | -0.647 | 0.648 | 0.641 | 0.078 | 0.322 |
| Q9TW45 | -0.298 | 0.13 | 1.27 | -0.282 | 0.694 | 0.863 | 0.11 | 0.709 |
| Q19591 | -0.3 | 0.33 | -0.421 | -0.563 | 0.627 | 0.594 | 0.498 | 0.367 |
| Q19286 | -0.301 | -0.59 | -0.0702 | -1.12 | 0.517 | 0.214 | 0.879 | 0.0264 |
| Q21268 | -0.31 | 0.695 | 0.293 | -0.0387 | 0.544 | 0.185 | 0.566 | 0.939 |
| Q27389 | -0.312 | 0.887 | -0.076 | 0.973 | 0.333 | 0.0128 | 0.81 | 0.00741 |
| Q18409 | -0.315 | -1.67 | 0.254 | -1.62 | 0.818 | 0.235 | 0.853 | 0.247 |
| Q23058 | -0.315 | 0.599 | 0.859 | -0.315 | 0.575 | 0.294 | 0.14 | 0.575 |
| Q8MXD9 | -0.315 | 0.619 | 0.308 | 0.0279 | 0.406 | 0.114 | 0.416 | 0.94 |
| H2KYC9 | -0.318 | 0.419 | 1.06 | 0.123 | 0.656 | 0.558 | 0.15 | 0.863 |
| G5ED07 | -0.319 | -0.751 | -0.485 | -0.687 | 0.412 | 0.0663 | 0.219 | 0.09 |
| A5HU98 | -0.323 | -0.497 | -0.568 | -0.22 | 0.696 | 0.548 | 0.494 | 0.79 |
| G5EFE3 | -0.328 | -1.53 | 0.546 | -1.02 | 0.385 | 0.000932 | 0.158 | 0.0148 |
| G5EC31 | -0.33 | 0.0916 | -0.57 | 0.826 | 0.638 | 0.896 | 0.42 | 0.248 |
| Q9BKU5 | -0.334 | 0.485 | -0.764 | 0.784 | 0.482 | 0.312 | 0.121 | 0.112 |
| O45948 | -0.334 | -1.26 | 1.49 | 0.189 | 0.825 | 0.409 | 0.332 | 0.9 |
| P51403 | -0.336 | 1.08 | -0.248 | 1 | 0.35 | 0.00772 | 0.488 | 0.0123 |
| Q9N368 | -0.341 | 0.565 | 0.773 | 0.445 | 0.355 | 0.136 | 0.0483 | 0.233 |
| Q19766 | -0.341 | -1.34 | -1.13 | -0.58 | 0.503 | 0.0176 | 0.0388 | 0.262 |
| G5ECX8 | -0.344 | 0.0157 | 1.15 | -0.0245 | 0.625 | 0.982 | 0.118 | 0.972 |
| Q22338 | -0.354 | 0.00603 | 0.11 | -0.482 | 0.558 | 0.992 | 0.855 | 0.427 |
| A0A4V0INA2 | -0.356 | -3.51 | 0.578 | -2.88 | 0.58 | 7.27E-05 | 0.374 | 0.00045 |
| Q21155 | -0.361 | -1.26 | 2.12 | -1.92 | 0.726 | 0.231 | 0.054 | 0.0771 |
| Q17435 | -0.364 | -0.28 | 1.14 | 0.608 | 0.547 | 0.643 | 0.0745 | 0.321 |
| P09446 | -0.367 | -0.0288 | -0.677 | -0.0173 | 0.241 | 0.925 | 0.0406 | 0.955 |
| Q22918 | -0.368 | -1.5 | -0.0666 | -1.15 | 0.441 | 0.00628 | 0.888 | 0.0265 |
| O45148 | -0.371 | -0.403 | -0.539 | -0.783 | 0.432 | 0.394 | 0.259 | 0.11 |
| O61742 | -0.376 | -1.47 | -0.643 | -2.03 | 0.597 | 0.0529 | 0.37 | 0.0112 |
| O44144 | -0.377 | -1.67 | 1.7 | -2.55 | 0.669 | 0.074 | 0.0699 | 0.0107 |
| Q9XW41 | -0.378 | -0.31 | 0.109 | -0.13 | 0.2 | 0.287 | 0.702 | 0.649 |
| Q9N358 | -0.383 | 1.19 | -1.05 | 0.887 | 0.535 | 0.0688 | 0.104 | 0.163 |
| O45418 | -0.384 | 0.601 | -0.0599 | 1.07 | 0.402 | 0.197 | 0.895 | 0.0311 |
| Q9U1X9 | -0.384 | 0.823 | 0.282 | 0.791 | 0.388 | 0.0773 | 0.523 | 0.0881 |
| G5EDT4 | -0.386 | 0.646 | 0.0102 | 0.861 | 0.24 | 0.0595 | 0.975 | 0.0162 |
| O16266 | -0.386 | -1.69 | 0.577 | -1.71 | 0.356 | 0.000939 | 0.175 | 0.000876 |
| Q9U3P5 | -0.387 | -0.778 | -1.3 | 1.73 | 0.735 | 0.499 | 0.267 | 0.145 |
| Q95Y51 | -0.388 | 0.463 | 0.256 | -0.514 | 0.459 | 0.379 | 0.624 | 0.33 |
| Q9XVS2 | -0.396 | -4.34 | -1.04 | -3.89 | 0.657 | 0.000221 | 0.256 | 0.000568 |
| P34661 | -0.397 | -0.234 | -1.52 | -0.665 | 0.561 | 0.731 | 0.0395 | 0.336 |
| Q9N599 | -0.402 | -0.316 | -1.5 | -0.082 | 0.613 | 0.69 | 0.0735 | 0.917 |
| P47991 | -0.403 | 0.491 | 0.423 | 0.728 | 0.257 | 0.171 | 0.235 | 0.0509 |
| Q95ZY7 | -0.408 | -1.61 | 0.42 | -0.941 | 0.454 | 0.00896 | 0.441 | 0.0978 |
| O16294 | -0.409 | -2.55 | -1.18 | -0.397 | 0.548 | 0.00186 | 0.0981 | 0.56 |
| P47209 | -0.421 | 0.598 | -0.104 | 0.323 | 0.188 | 0.0697 | 0.738 | 0.307 |
| Q22053 | -0.424 | 0.835 | 0.0785 | 0.872 | 0.217 | 0.0233 | 0.814 | 0.0187 |
| P50140 | -0.424 | -0.733 | -0.582 | -0.348 | 0.201 | 0.036 | 0.0864 | 0.288 |

| | | | | | | | | |
|---|---|---|---|---|---|---|---|---|
| Q20683 | -0.428 | 0.537 | 1.02 | -0.123 | 0.477 | 0.376 | 0.103 | 0.837 |
| Q10039 | -0.431 | 1.1 | -0.815 | 0.777 | 0.384 | 0.0376 | 0.111 | 0.128 |
| V6CKJ1 | -0.433 | -1.08 | 0.756 | -0.779 | 0.268 | 0.0125 | 0.0641 | 0.0571 |
| Q9NEZ5 | -0.436 | -0.237 | 0.379 | -0.742 | 0.189 | 0.466 | 0.25 | 0.0342 |
| A0A2K5A<br>U04 | -0.437 | 0.002 | -0.297 | -0.51 | 0.588 | 0.998 | 0.712 | 0.528 |
| Q7YZH4 | -0.437 | -0.061 | 0.996 | -0.341 | 0.461 | 0.917 | 0.106 | 0.564 |
| Q9N5B3 | -0.45 | 0.717 | -0.302 | 0.493 | 0.14 | 0.0261 | 0.312 | 0.109 |
| Q09533 | -0.452 | 0.793 | 0.0735 | 0.93 | 0.205 | 0.0352 | 0.832 | 0.0161 |
| Q9BKQ9 | -0.461 | -1.23 | -0.00734 | -1.25 | 0.257 | 0.00706 | 0.985 | 0.00645 |
| O16785 | -0.462 | 0.554 | 1.52 | -0.567 | 0.589 | 0.518 | 0.0893 | 0.508 |
| H2KZZ2 | -0.463 | -1.24 | 0.757 | -1.3 | 0.4 | 0.0359 | 0.177 | 0.0291 |
| O45946 | -0.463 | -0.068 | 0.932 | 0.245 | 0.305 | 0.878 | 0.0504 | 0.581 |
| P50432 | -0.464 | -0.0827 | -0.109 | 0.224 | 0.165 | 0.798 | 0.734 | 0.49 |
| Q20970 | -0.47 | -0.624 | 0.536 | -0.154 | 0.255 | 0.138 | 0.198 | 0.703 |
| Q21323 | -0.476 | -0.144 | 0.612 | 0.00455 | 0.483 | 0.83 | 0.37 | 0.995 |
| Q09936 | -0.479 | -1.06 | 0.502 | -1.27 | 0.32 | 0.039 | 0.298 | 0.0162 |
| Q9XWM1 | -0.48 | 0.206 | 0.786 | 0.00703 | 0.368 | 0.696 | 0.15 | 0.989 |
| Q2EEM8 | -0.482 | -4.72 | 0.128 | -4.18 | 0.344 | 1.80E-07 | 0.799 | 7.73E-07 |
| Q27371 | -0.489 | -0.808 | 0.356 | -1.34 | 0.182 | 0.0361 | 0.324 | 0.0018 |
| O17953 | -0.499 | -4.69 | -1.1 | -1.43 | 0.253 | 2.60E-08 | 0.0199 | 0.00419 |
| O61793 | -0.502 | -1.7 | -0.605 | -0.702 | 0.383 | 0.00868 | 0.296 | 0.229 |
| G5EE65 | -0.502 | 0.948 | 0.32 | -0.29 | 0.539 | 0.254 | 0.694 | 0.722 |
| Q22341 | -0.503 | 1.09 | -0.114 | 0.599 | 0.495 | 0.151 | 0.876 | 0.418 |
| O18650 | -0.506 | 2.06 | 0.135 | 1.87 | 0.129 | 1.35E-05 | 0.672 | 3.67E-05 |
| P91020 | -0.506 | -0.406 | -0.721 | -0.142 | 0.147 | 0.238 | 0.0461 | 0.674 |
| O45904 | -0.507 | 1.14 | 0.516 | 0.286 | 0.259 | 0.0197 | 0.251 | 0.517 |
| Q95PW9 | -0.507 | -0.56 | 0.0158 | -0.362 | 0.226 | 0.184 | 0.969 | 0.381 |
| G5EE04 | -0.514 | -0.609 | 0.441 | -0.377 | 0.11 | 0.0624 | 0.165 | 0.23 |
| G5ED29 | -0.515 | -2.78 | 0.329 | -2.58 | 0.32 | 7.23E-05 | 0.521 | 0.000151 |
| O01532 | -0.515 | 0.441 | 1.39 | -0.443 | 0.406 | 0.475 | 0.0362 | 0.473 |
| Q9BKU4 | -0.515 | -1.56 | -0.56 | -2.27 | 0.524 | 0.0682 | 0.49 | 0.0124 |
| Q95ZL3 | -0.516 | -0.722 | 0.812 | -0.638 | 0.326 | 0.176 | 0.132 | 0.229 |
| Q21465 | -0.517 | -0.136 | -0.261 | -1.74 | 0.597 | 0.889 | 0.789 | 0.09 |
| O45734 | -0.518 | -0.926 | -0.299 | -1.05 | 0.134 | 0.013 | 0.374 | 0.00635 |
| Q9N2W7 | -0.528 | -0.84 | -0.355 | -1.04 | 0.207 | 0.0539 | 0.389 | 0.0208 |
| Q9NA39 | -0.534 | -1.05 | 0.629 | -0.856 | 0.209 | 0.0218 | 0.144 | 0.0535 |
| Q9NEU5 | -0.534 | 0.774 | 1.49 | -0.44 | 0.566 | 0.409 | 0.124 | 0.636 |
| O17725 | -0.537 | 0.862 | 0.618 | -0.167 | 0.183 | 0.0412 | 0.129 | 0.669 |
| Q94300 | -0.545 | 1.18 | -0.416 | 1.14 | 0.341 | 0.0517 | 0.465 | 0.0584 |
| G5EDE4 | -0.545 | -1.15 | 1.31 | -0.862 | 0.232 | 0.0198 | 0.00979 | 0.0688 |
| G5EFM6 | -0.545 | -2.39 | 0.351 | -2.89 | 0.318 | 0.000478 | 0.515 | 8.19E-05 |
| Q9U1Q4 | -0.545 | -0.352 | -0.77 | -0.766 | 0.646 | 0.766 | 0.518 | 0.52 |
| O61848 | -0.551 | -1.69 | 0.835 | -1.62 | 0.226 | 0.00172 | 0.0757 | 0.00236 |
| Q22037 | -0.552 | -0.897 | 0.125 | -0.787 | 0.153 | 0.0278 | 0.736 | 0.0492 |
| G5ED01 | -0.555 | 1.2 | 0.885 | 0.762 | 0.212 | 0.0136 | 0.0558 | 0.0941 |
| H2KZR3 | -0.561 | -1.12 | 0.471 | -2.26 | 0.302 | 0.0517 | 0.384 | 0.000726 |

| | | | | | | | | |
|---|---|---|---|---|---|---|---|---|
| G5EDQ2 | -0.563 | -0.53 | 0.702 | -0.406 | 0.307 | 0.335 | 0.207 | 0.457 |
| P34286 | -0.565 | -2.1 | -0.295 | -1.35 | 0.325 | 0.00201 | 0.603 | 0.0293 |
| P53013 | -0.565 | 0.377 | -0.265 | 0.446 | 0.112 | 0.277 | 0.44 | 0.202 |
| Q9UB28 | -0.565 | -0.687 | 0.947 | -1.18 | 0.459 | 0.37 | 0.223 | 0.133 |
| Q19706 | -0.576 | 0.00404 | 0.504 | -0.273 | 0.153 | 0.992 | 0.208 | 0.486 |
| Q93576 | -0.576 | -0.35 | -0.342 | 0.22 | 0.119 | 0.329 | 0.34 | 0.535 |
| Q9U2K8 | -0.578 | -0.918 | 1.02 | 0.572 | 0.444 | 0.232 | 0.187 | 0.449 |
| P49405 | -0.583 | 0.922 | 0.00985 | 0.956 | 0.081 | 0.0101 | 0.975 | 0.0082 |
| O76577 | -0.59 | -0.887 | -0.0249 | -0.576 | 0.172 | 0.0483 | 0.952 | 0.182 |
| H2L0F6 | -0.596 | -2.54 | 0.695 | -3.39 | 0.348 | 0.00103 | 0.276 | 7.81E-05 |
| O16298 | -0.596 | -3.28 | -1.02 | -4.61 | 0.172 | 1.72E-06 | 0.0271 | 2.88E-08 |
| P18948 | -0.597 | 0.716 | -0.0379 | 1.24 | 0.385 | 0.3 | 0.955 | 0.0846 |
| P52821 | -0.602 | 1.27 | -0.59 | 1.22 | 0.223 | 0.018 | 0.232 | 0.0222 |
| P55853 | -0.602 | 0.0917 | -0.84 | 0.241 | 0.129 | 0.809 | 0.041 | 0.528 |
| P91374 | -0.602 | 0.171 | -0.526 | 0.622 | 0.164 | 0.683 | 0.22 | 0.152 |
| Q1XFY2 | -0.602 | -0.233 | 1.84 | 0.092 | 0.303 | 0.685 | 0.00574 | 0.872 |
| Q20276 | -0.607 | 0.812 | -1.42 | 0.36 | 0.312 | 0.182 | 0.0278 | 0.544 |
| P37806 | -0.61 | -1.71 | 0.611 | -1.98 | 0.122 | 0.000409 | 0.121 | 0.000108 |
| Q10661 | -0.612 | -2.26 | 0.0954 | -2.88 | 0.693 | 0.16 | 0.951 | 0.0791 |
| D0PV95 | -0.614 | 0.19 | -0.626 | -0.201 | 0.305 | 0.747 | 0.296 | 0.732 |
| P61866 | -0.617 | 1.03 | -0.382 | 0.393 | 0.0814 | 0.00737 | 0.264 | 0.251 |
| O61880 | -0.617 | 0.104 | -0.13 | -1.57 | 0.235 | 0.837 | 0.797 | 0.00699 |
| Q95QA6 | -0.631 | -1.33 | 0.677 | -1.44 | 0.0891 | 0.00182 | 0.07 | 0.000993 |
| Q09567 | -0.634 | -2.22 | -0.135 | -1.56 | 0.154 | 0.000124 | 0.752 | 0.00243 |
| Q9XVA9 | -0.634 | -1.81 | 1.36 | -1.14 | 0.333 | 0.0128 | 0.0501 | 0.0931 |
| Q966I8 | -0.634 | -0.163 | -0.292 | -1.18 | 0.345 | 0.806 | 0.66 | 0.0915 |
| D3NQB1 | -0.643 | 0.229 | 1.01 | -0.173 | 0.498 | 0.808 | 0.291 | 0.854 |
| G5ECT6 | -0.644 | -2.16 | 0.738 | -0.619 | 0.246 | 0.00121 | 0.187 | 0.264 |
| P48727 | -0.658 | 0.86 | 0.677 | -0.0995 | 0.145 | 0.0634 | 0.135 | 0.819 |
| P50880 | -0.658 | 0.591 | -0.475 | 0.774 | 0.0891 | 0.123 | 0.208 | 0.0498 |
| Q9XXD4 | -0.659 | 0.599 | -1.18 | -0.497 | 0.425 | 0.468 | 0.163 | 0.546 |
| Q564Q6 | -0.66 | 0.814 | 0.737 | 1.16 | 0.283 | 0.19 | 0.233 | 0.071 |
| Q09531 | -0.674 | -1.49 | 0.92 | -1.3 | 0.283 | 0.0271 | 0.15 | 0.0501 |
| P41996 | -0.679 | 0.266 | 1.33 | 1.27 | 0.542 | 0.81 | 0.242 | 0.263 |
| P91457 | -0.681 | -2 | -0.179 | -2.42 | 0.33 | 0.0105 | 0.794 | 0.00306 |
| Q966J6 | -0.681 | -1.78 | 0.0342 | -2.77 | 0.279 | 0.0107 | 0.956 | 0.000442 |
| Q9BKU3 | -0.681 | 0.26 | 0.242 | -1.04 | 0.166 | 0.585 | 0.612 | 0.0422 |
| G5EC87 | -0.686 | -1.46 | -0.0118 | -1.82 | 0.304 | 0.0393 | 0.986 | 0.0136 |
| H2KZY6 | -0.693 | -1.46 | -2.21 | -2.08 | 0.3 | 0.0397 | 0.00405 | 0.0062 |
| Q965K5 | -0.695 | 0.505 | 1.67 | 0.271 | 0.311 | 0.458 | 0.0244 | 0.687 |
| O44444 | -0.706 | -2.64 | 0.591 | -3.05 | 0.163 | 8.18E-05 | 0.239 | 1.93E-05 |
| G5EF87 | -0.706 | 0.0466 | -0.266 | -1.15 | 0.262 | 0.94 | 0.666 | 0.0782 |
| Q9U2Q8 | -0.708 | -3.23 | 0.2 | -2.64 | 0.407 | 0.00163 | 0.813 | 0.00658 |
| Q19246 | -0.709 | -0.897 | -0.625 | -1.74 | 0.403 | 0.293 | 0.459 | 0.0522 |
| O02056 | -0.71 | 0.146 | -0.48 | 0.772 | 0.0569 | 0.676 | 0.182 | 0.0407 |
| O01814 | -0.714 | 0.121 | 0.151 | 0.276 | 0.209 | 0.826 | 0.784 | 0.618 |
| Q21993 | -0.715 | -0.735 | -1.03 | -1.2 | 0.499 | 0.487 | 0.333 | 0.265 |

| | | | | | | | | |
|---|---|---|---|---|---|---|---|---|
| G5ECM9 | -0.722 | -0.4 | 0.278 | -0.651 | 0.244 | 0.512 | 0.646 | 0.291 |
| O16462 | -0.726 | 0.075 | 0.0419 | -0.136 | 0.236 | 0.9 | 0.944 | 0.819 |
| P41988 | -0.727 | 1.13 | 0.183 | 0.86 | 0.112 | 0.0198 | 0.677 | 0.0651 |
| Q17827 | -0.728 | -1.43 | 0.703 | -1.93 | 0.131 | 0.0072 | 0.143 | 0.000822 |
| Q95PW6 | -0.728 | -0.737 | -0.654 | -1.78 | 0.163 | 0.158 | 0.206 | 0.00292 |
| O44145 | -0.729 | -3.59 | 0.169 | -0.474 | 0.255 | 4.61E-05 | 0.788 | 0.453 |
| Q9N4I3 | -0.733 | -0.446 | 0.399 | -0.858 | 0.0639 | 0.241 | 0.291 | 0.0337 |
| Q9TXP0 | -0.734 | 0.658 | 0.298 | 0.412 | 0.0273 | 0.0443 | 0.333 | 0.187 |
| P90849 | -0.735 | -5.31 | -0.661 | -5.65 | 0.053 | 4.97E-10 | 0.0782 | 2.20E-10 |
| Q22716 | -0.735 | 1.03 | -0.407 | 1.26 | 0.0584 | 0.012 | 0.273 | 0.00345 |
| Q11099 | -0.735 | 0.335 | 0.572 | -0.0732 | 0.27 | 0.608 | 0.386 | 0.911 |
| Q17832 | -0.741 | 0.428 | -0.248 | 0.204 | 0.0194 | 0.149 | 0.391 | 0.479 |
| P34447 | -0.746 | -0.0264 | 0.195 | -0.0442 | 0.323 | 0.972 | 0.792 | 0.952 |
| Q19057 | -0.746 | -1.09 | -0.311 | -0.0916 | 0.218 | 0.081 | 0.599 | 0.876 |
| O45226 | -0.748 | -1.55 | -1.27 | 1.15 | 0.577 | 0.258 | 0.35 | 0.393 |
| Q20796 | -0.752 | -0.0903 | 0.0529 | -0.178 | 0.131 | 0.85 | 0.912 | 0.71 |
| G5ECY0 | -0.753 | -0.71 | 0.159 | -1.26 | 0.0429 | 0.0543 | 0.645 | 0.00223 |
| P48158 | -0.756 | 0.866 | 0.395 | 0.858 | 0.0477 | 0.0262 | 0.275 | 0.0275 |
| Q18885 | -0.758 | -0.482 | 0.142 | -0.264 | 0.0771 | 0.245 | 0.726 | 0.518 |
| Q09635 | -0.762 | 0.272 | 0.054 | 0.367 | 0.164 | 0.609 | 0.918 | 0.49 |
| O17406 | -0.763 | -0.506 | 0.261 | -0.897 | 0.0763 | 0.224 | 0.523 | 0.0412 |
| P34504 | -0.765 | 0.399 | 0.265 | -0.144 | 0.164 | 0.456 | 0.619 | 0.786 |
| Q8WTJ4 | -0.766 | 0.983 | 1.23 | -0.707 | 0.267 | 0.16 | 0.0838 | 0.303 |
| P91318 | -0.767 | 0.032 | 1.28 | 0.0776 | 0.18 | 0.954 | 0.0332 | 0.888 |
| Q09390 | -0.768 | -1.72 | -0.842 | -3.66 | 0.279 | 0.0245 | 0.237 | 0.000103 |
| G5EEH9 | -0.769 | -2.26 | 0.0685 | -2.26 | 0.169 | 0.000824 | 0.899 | 0.00081 |
| Q965S8 | -0.773 | -0.611 | -0.737 | -0.85 | 0.288 | 0.398 | 0.311 | 0.245 |
| G5EBP5 | -0.774 | -2.5 | 0.609 | -3.67 | 0.134 | 0.000158 | 0.231 | 2.90E-06 |
| Q9BL03 | -0.775 | -1.42 | -0.285 | -1.38 | 0.0582 | 0.00207 | 0.46 | 0.00258 |
| Q20206 | -0.776 | 0.56 | -0.0958 | 0.719 | 0.0369 | 0.118 | 0.78 | 0.0509 |
| O17389 | -0.78 | -0.411 | 0.323 | -0.621 | 0.0193 | 0.185 | 0.291 | 0.0537 |
| O44156 | -0.783 | -2.18 | 0.307 | -2.84 | 0.321 | 0.0126 | 0.693 | 0.0023 |
| G5EEI4 | -0.794 | -0.498 | 0.244 | -0.398 | 0.0912 | 0.274 | 0.586 | 0.378 |
| C6KRN1 | -0.795 | 0.0602 | 0.236 | -0.352 | 0.121 | 0.902 | 0.631 | 0.477 |
| G5EGK1 | -0.798 | -1.85 | 0.137 | -0.952 | 0.405 | 0.0663 | 0.885 | 0.323 |
| Q9N350 | -0.8 | -2.12 | -0.288 | 0.533 | 0.247 | 0.00656 | 0.67 | 0.434 |
| A0A164D3G3 | -0.806 | -0.759 | 1.09 | -1.37 | 0.175 | 0.2 | 0.0743 | 0.0298 |
| O62220 | -0.807 | 0.0796 | 0.584 | -0.461 | 0.0235 | 0.805 | 0.087 | 0.168 |
| O01806 | -0.813 | -0.139 | 0.00353 | -0.0428 | 0.0439 | 0.711 | 0.992 | 0.909 |
| Q9NAH3 | -0.816 | -0.552 | -0.664 | -0.896 | 0.101 | 0.255 | 0.175 | 0.0746 |
| Q23121 | -0.819 | -1 | -0.378 | -1.06 | 0.0286 | 0.00998 | 0.279 | 0.00719 |
| O01576 | -0.819 | 0.297 | 0.526 | -0.0124 | 0.127 | 0.566 | 0.315 | 0.981 |
| A0A0K3AXM4 | -0.82 | -1.39 | 0.245 | -0.157 | 0.168 | 0.0278 | 0.67 | 0.785 |
| Q9GRZ0 | -0.825 | -0.589 | 0.783 | -0.363 | 0.235 | 0.391 | 0.259 | 0.593 |
| Q9N408 | -0.828 | -1.58 | 0.0188 | -0.781 | 0.138 | 0.00937 | 0.972 | 0.159 |

| | | | | | | | | |
|---|---|---|---|---|---|---|---|---|
| Q17849 | -0.829 | -3.62 | -0.167 | -2.46 | 0.0451 | 1.72E-07 | 0.663 | 1.45E-05 |
| Q9U2H9 | -0.83 | 0.0831 | 0.375 | 0.0862 | 0.0304 | 0.813 | 0.295 | 0.806 |
| O76840 | -0.832 | -0.633 | 0.275 | -0.563 | 0.0238 | 0.0743 | 0.415 | 0.108 |
| Q23050 | -0.838 | -1.9 | 0.442 | -1.81 | 0.0379 | 0.000143 | 0.246 | 0.000222 |
| Q966C6 | -0.848 | 0.562 | -0.371 | 0.839 | 0.0174 | 0.0952 | 0.257 | 0.0183 |
| Q93572 | -0.849 | 0.781 | -0.308 | 0.559 | 0.0295 | 0.0427 | 0.394 | 0.132 |
| P34563 | -0.867 | 0.0376 | -0.968 | 0.272 | 0.0158 | 0.907 | 0.00843 | 0.402 |
| Q21735 | -0.871 | -0.594 | 0.884 | 0.0181 | 0.127 | 0.287 | 0.122 | 0.974 |
| P47207 | -0.88 | 0.379 | -0.127 | 0.46 | 0.0157 | 0.256 | 0.698 | 0.173 |
| Q22135 | -0.883 | 0.491 | 0.556 | 1.17 | 0.224 | 0.491 | 0.436 | 0.114 |
| A0A486W<br>X27 | -0.885 | -1.34 | 0.36 | -1.14 | 0.291 | 0.119 | 0.663 | 0.18 |
| O61977 | -0.895 | -1.49 | 0.42 | -2.09 | 0.102 | 0.0112 | 0.425 | 0.00113 |
| P53596 | -0.896 | -0.937 | -0.0179 | 0.187 | 0.108 | 0.0941 | 0.973 | 0.725 |
| P91128 | -0.899 | 0.682 | -0.458 | 0.796 | 0.0142 | 0.0519 | 0.175 | 0.0265 |
| P48156 | -0.905 | -0.0121 | -0.123 | 0.4 | 0.0247 | 0.974 | 0.737 | 0.284 |
| H2L056 | -0.907 | 0.0228 | 2.01 | -0.728 | 0.278 | 0.978 | 0.0258 | 0.38 |
| Q23487 | -0.913 | -0.429 | -0.243 | -0.136 | 0.0243 | 0.254 | 0.512 | 0.711 |
| G5EDM4 | -0.914 | -1.35 | 0.00109 | -1.2 | 0.00878 | 0.00051 | 0.997 | 0.00131 |
| Q95XG6 | -0.915 | -1.09 | -0.826 | -2.14 | 0.275 | 0.198 | 0.322 | 0.0187 |
| Q8MXJ1 | -0.919 | -1.17 | 0.471 | -1.5 | 0.0203 | 0.00499 | 0.2 | 0.000809 |
| G5EGB1 | -0.922 | -4 | 0.196 | -1.2 | 0.1 | 2.59E-06 | 0.714 | 0.0381 |
| Q22288 | -0.925 | -1.93 | 0.464 | -0.844 | 0.0454 | 0.000435 | 0.288 | 0.0647 |
| O16259 | -0.926 | -0.262 | 0.714 | -0.11 | 0.0672 | 0.583 | 0.149 | 0.816 |
| A3QMC5 | -0.927 | 0.599 | -0.429 | 1.13 | 0.0102 | 0.0758 | 0.191 | 0.00283 |
| A6ZJ46 | -0.929 | -3.02 | -0.247 | -4.05 | 0.139 | 0.000168 | 0.683 | 8.62E-06 |
| Q09601 | -0.936 | 0.268 | 2.08 | -0.532 | 0.2 | 0.705 | 0.00987 | 0.457 |
| O18240 | -0.941 | 0.175 | -0.266 | 1.19 | 0.0446 | 0.687 | 0.542 | 0.0142 |
| G5EFC4 | -0.943 | 0.206 | 0.987 | 0.243 | 0.0559 | 0.656 | 0.0468 | 0.6 |
| Q95XS2 | -0.943 | 0.32 | 0.397 | -1.18 | 0.271 | 0.703 | 0.637 | 0.172 |
| Q21793 | -0.944 | -0.692 | 0.54 | -2.76 | 0.101 | 0.219 | 0.332 | 0.00016 |
| Q20877 | -0.945 | -0.606 | -0.752 | -1.01 | 0.108 | 0.29 | 0.194 | 0.0872 |
| O45499 | -0.955 | 0.56 | -0.41 | 0.511 | 0.0126 | 0.115 | 0.239 | 0.148 |
| O45319 | -0.96 | -0.817 | 0.339 | -0.676 | 0.0161 | 0.0355 | 0.349 | 0.0745 |
| Q18577 | -0.961 | -0.589 | 0.276 | -0.494 | 0.0179 | 0.122 | 0.454 | 0.189 |
| Q18678 | -0.968 | -1.33 | 0.303 | -0.652 | 0.127 | 0.0432 | 0.62 | 0.293 |
| Q21763 | -0.972 | 0.543 | -0.0593 | 0.0747 | 0.0111 | 0.124 | 0.861 | 0.825 |
| A0A1N7S<br>YQ6 | -0.973 | -1.22 | 0.027 | -1.05 | 0.0139 | 0.00334 | 0.939 | 0.00893 |
| Q03561 | -0.975 | 0.477 | 0.998 | -0.443 | 0.139 | 0.456 | 0.131 | 0.487 |
| A0A9J5H<br>VW4 | -0.981 | -0.408 | 0.184 | -0.503 | 0.0698 | 0.428 | 0.718 | 0.33 |
| P15796 | -0.983 | -0.588 | 0.382 | -0.623 | 0.0591 | 0.239 | 0.438 | 0.213 |
| Q20334 | -0.986 | -0.471 | 0.472 | -0.265 | 0.111 | 0.429 | 0.428 | 0.654 |
| V6CJ04 | -0.995 | 0.0305 | -0.341 | -0.576 | 0.067 | 0.952 | 0.507 | 0.27 |
| Q9XXR2 | -0.998 | 0.418 | 1.02 | -0.154 | 0.193 | 0.575 | 0.184 | 0.836 |
| P10771 | -1 | -0.633 | 0.266 | -1.21 | 0.0198 | 0.118 | 0.496 | 0.00672 |

| | | | | | | | | |
|---|---|---|---|---|---|---|---|---|
| P90978 | -1.01 | -2.09 | 0.854 | -2.19 | 0.217 | 0.018 | 0.293 | 0.0141 |
| Q20647 | -1.01 | -0.984 | -0.118 | -1.54 | 0.0482 | 0.0525 | 0.802 | 0.00506 |
| Q9NEN6 | -1.01 | -0.0103 | -0.924 | 0.287 | 0.0154 | 0.978 | 0.0242 | 0.445 |
| O44471 | -1.02 | -1.37 | 0.912 | -1.1 | 0.0289 | 0.00578 | 0.0479 | 0.0204 |
| O45146 | -1.02 | 0.0651 | 1.03 | -0.812 | 0.194 | 0.932 | 0.188 | 0.294 |
| Q9XWP7 | -1.02 | -1.49 | 0.306 | -0.413 | 0.0494 | 0.00698 | 0.528 | 0.396 |
| O76561 | -1.02 | -1.59 | -1.14 | -1.27 | 0.469 | 0.268 | 0.422 | 0.371 |
| P34689 | -1.03 | -0.877 | 1.72 | -1.6 | 0.272 | 0.345 | 0.0764 | 0.0969 |
| G5EFF7 | -1.04 | -1.4 | 0.105 | -0.559 | 0.0316 | 0.00616 | 0.813 | 0.22 |
| G5EDV0 | -1.04 | -0.306 | -0.0449 | 1.54 | 0.328 | 0.77 | 0.966 | 0.157 |
| Q23680 | -1.05 | -2.8 | 0.391 | -0.226 | 0.234 | 0.00524 | 0.651 | 0.792 |
| Q9U263 | -1.05 | 0.284 | 1.47 | -0.122 | 0.155 | 0.69 | 0.0536 | 0.864 |
| Q19335 | -1.05 | -0.00768 | 1.93 | -0.498 | 0.383 | 0.995 | 0.119 | 0.675 |
| Q20724 | -1.05 | 0.0705 | -0.00819 | -0.905 | 0.0826 | 0.902 | 0.989 | 0.13 |
| P91027 | -1.06 | -1.43 | -0.354 | -1.7 | 0.161 | 0.0668 | 0.63 | 0.0334 |
| Q0G820 | -1.07 | -0.46 | 1.27 | -0.252 | 0.219 | 0.59 | 0.15 | 0.766 |
| Q9XWG2 | -1.07 | -3.48 | -0.249 | -2.87 | 0.18 | 0.00043 | 0.747 | 0.00203 |
| Q9BL19 | -1.08 | 0.632 | -0.905 | 0.0968 | 0.0286 | 0.175 | 0.0601 | 0.83 |
| Q9XVS4 | -1.08 | 0.046 | -0.054 | 0.51 | 0.0169 | 0.91 | 0.894 | 0.221 |
| O02637 | -1.08 | -1.11 | 0.978 | -1.08 | 0.151 | 0.141 | 0.191 | 0.151 |
| P34594 | -1.09 | -0.134 | -0.838 | -1.34 | 0.238 | 0.881 | 0.358 | 0.152 |
| Q09930 | -1.1 | -0.144 | -1.31 | 0.726 | 0.144 | 0.842 | 0.0862 | 0.323 |
| Q9Y0V6 | -1.1 | -2.57 | 0.124 | -2.88 | 0.205 | 0.0077 | 0.883 | 0.00363 |
| Q9XWN4 | -1.1 | -3.29 | 1.04 | -4.32 | 0.0301 | 4.55E-06 | 0.0383 | 1.93E-07 |
| Q9XW37 | -1.1 | -0.476 | -0.0978 | -1.02 | 0.0187 | 0.27 | 0.817 | 0.028 |
| Q21568 | -1.11 | -0.801 | -0.158 | -0.748 | 0.00648 | 0.0368 | 0.655 | 0.049 |
| Q9N4G9 | -1.11 | -2.16 | 0.278 | -1.69 | 0.0721 | 0.00209 | 0.634 | 0.0105 |
| Q95ZQ5 | -1.11 | -1.05 | -0.123 | -2.79 | 0.196 | 0.223 | 0.883 | 0.00435 |
| P34416 | -1.11 | 0.497 | -0.223 | -0.426 | 0.0901 | 0.427 | 0.719 | 0.495 |
| Q09581 | -1.12 | -1.94 | -0.268 | -1.44 | 0.00666 | 7.95E-05 | 0.458 | 0.0011 |
| Q9BKS1 | -1.12 | 0.185 | 0.219 | -0.459 | 0.0276 | 0.69 | 0.638 | 0.33 |
| Q9TZE4 | -1.12 | -0.18 | 1.8 | -0.213 | 0.133 | 0.801 | 0.0229 | 0.766 |
| Q19191 | -1.13 | -1.7 | 0.343 | -1.63 | 0.0275 | 0.00237 | 0.467 | 0.00329 |
| D5MCR4 | -1.13 | -1.41 | 0.323 | -1.42 | 0.00301 | 0.000535 | 0.323 | 0.000507 |
| Q9N3B0 | -1.13 | -2.13 | -1.26 | -1.84 | 0.198 | 0.0239 | 0.155 | 0.0461 |
| Q22719 | -1.14 | -2.22 | 0.0825 | -1.68 | 0.016 | 0.000106 | 0.845 | 0.00125 |
| O44451 | -1.15 | -1.62 | -1.35 | -1.82 | 0.17 | 0.0615 | 0.112 | 0.0387 |
| Q20363 | -1.15 | -0.318 | -1.03 | 0.154 | 0.0609 | 0.582 | 0.0907 | 0.79 |
| C7IVR4 | -1.16 | 0.456 | 0.363 | 0.0331 | 0.0325 | 0.365 | 0.468 | 0.947 |
| O01974 | -1.16 | -1.67 | 0.137 | -1.14 | 0.0572 | 0.01 | 0.81 | 0.0615 |
| H2L2A0 | -1.16 | -1.99 | 0.377 | -3.14 | 0.0501 | 0.00253 | 0.497 | 4.81E-05 |
| O01804 | -1.16 | 0.55 | 0.623 | -0.782 | 0.012 | 0.193 | 0.144 | 0.0725 |
| P91398 | -1.16 | 0.385 | 0.101 | 0.0144 | 0.0462 | 0.479 | 0.851 | 0.979 |
| Q20774 | -1.16 | -0.447 | -0.331 | -0.142 | 0.0394 | 0.395 | 0.527 | 0.785 |
| Q21559 | -1.16 | -0.595 | 0.148 | -0.784 | 0.123 | 0.414 | 0.837 | 0.286 |
| O17536 | -1.17 | -0.469 | 1.76 | -0.671 | 0.211 | 0.606 | 0.0682 | 0.464 |
| P06125 | -1.18 | 1.13 | 0.0105 | 1.55 | 0.00852 | 0.0109 | 0.979 | 0.00132 |

| | | | | | | | | |
|---|---|---|---|---|---|---|---|---|
| P19974 | -1.18 | -1.31 | -0.51 | -0.979 | 0.00416 | 0.002 | 0.16 | 0.0131 |
| A0A1I6C M87 | -1.2 | 0.159 | 0.482 | -0.436 | 0.0213 | 0.736 | 0.315 | 0.362 |
| Q9NEW6 | -1.21 | -0.579 | 0.519 | -0.571 | 0.0151 | 0.205 | 0.253 | 0.211 |
| Q9XUV0 | -1.22 | -0.495 | -0.157 | -0.402 | 0.0124 | 0.262 | 0.716 | 0.36 |
| O01868 | -1.23 | 0.00357 | 0.341 | -0.186 | 0.0149 | 0.994 | 0.453 | 0.68 |
| O02639 | -1.23 | 0.329 | 0.276 | 0.228 | 0.0013 | 0.302 | 0.384 | 0.47 |
| A7DT45 | -1.23 | -3.29 | -0.146 | -1.72 | 0.032 | 1.83E-05 | 0.781 | 0.00498 |
| Q9XWS4 | -1.23 | 0.143 | -0.36 | 0.0401 | 0.00346 | 0.687 | 0.319 | 0.91 |
| O62213 | -1.24 | -0.967 | 0.596 | -1.3 | 0.0104 | 0.0366 | 0.176 | 0.00773 |
| Q9XTU9 | -1.24 | -0.284 | -0.0368 | -0.473 | 0.0494 | 0.629 | 0.95 | 0.424 |
| Q10020 | -1.25 | -1.36 | 0.136 | -1.1 | 0.0418 | 0.0284 | 0.81 | 0.068 |
| Q22993 | -1.25 | -0.255 | -0.162 | -0.00283 | 0.0196 | 0.6 | 0.738 | 0.995 |
| E3W759 | -1.25 | -0.388 | 0.122 | -0.805 | 0.073 | 0.555 | 0.851 | 0.23 |
| P34334 | -1.26 | 0.293 | 0.305 | 0.547 | 0.00168 | 0.381 | 0.363 | 0.114 |
| Q9N4D8 | -1.26 | -0.464 | 0.515 | -1.31 | 0.0454 | 0.431 | 0.383 | 0.0382 |
| Q9TYV5 | -1.26 | -0.85 | 0.532 | -1.1 | 0.00474 | 0.0405 | 0.179 | 0.011 |
| P91918 | -1.26 | 1.41 | -0.186 | -0.906 | 0.213 | 0.167 | 0.85 | 0.363 |
| O44572 | -1.27 | 0.628 | 0.54 | 0.253 | 0.271 | 0.579 | 0.633 | 0.822 |
| Q9XTV4 | -1.28 | -4.35 | 0.569 | -3.96 | 0.0127 | 1.60E-07 | 0.226 | 4.92E-07 |
| G5EFB5 | -1.29 | -0.262 | 1.11 | -0.557 | 0.0551 | 0.677 | 0.0939 | 0.382 |
| H2KZV8 | -1.3 | -2.09 | 1.02 | -2.2 | 0.0728 | 0.00775 | 0.151 | 0.00562 |
| Q18943 | -1.3 | 1.78 | -0.243 | 1.94 | 0.326 | 0.186 | 0.852 | 0.151 |
| Q95YC6 | -1.3 | -0.481 | -0.422 | 0.339 | 0.176 | 0.606 | 0.651 | 0.715 |
| Q09511 | -1.32 | -2.8 | 0.0404 | -3.34 | 0.158 | 0.00686 | 0.964 | 0.00205 |
| Q10018 | -1.33 | -1.26 | 0.349 | -1.46 | 0.0201 | 0.0263 | 0.504 | 0.0126 |
| Q067X2 | -1.33 | 1.06 | 0.104 | 1.28 | 0.108 | 0.194 | 0.895 | 0.12 |
| Q9XW16 | -1.34 | -0.141 | -0.148 | 0.36 | 0.0027 | 0.706 | 0.692 | 0.342 |
| O17607 | -1.36 | -1.87 | 0.67 | -1.89 | 0.0996 | 0.0298 | 0.401 | 0.0287 |
| Q10009 | -1.36 | -0.371 | 0.597 | -0.861 | 0.0324 | 0.527 | 0.315 | 0.155 |
| Q21351 | -1.36 | -0.317 | -0.0598 | 0.107 | 0.00104 | 0.35 | 0.858 | 0.75 |
| Q9XW17 | -1.36 | -0.0138 | 0.523 | 0.296 | 0.00464 | 0.973 | 0.216 | 0.475 |
| Q9NAF9 | -1.37 | -0.521 | 0.529 | -0.901 | 0.0375 | 0.395 | 0.388 | 0.152 |
| P48150 | -1.38 | 0.511 | 0.0649 | 0.681 | 0.00061 | 0.124 | 0.838 | 0.047 |
| O62277 | -1.38 | -1.21 | -0.984 | 0.0115 | 0.00107 | 0.00287 | 0.0109 | 0.973 |
| O61749 | -1.38 | -0.79 | 0.669 | -0.391 | 0.1 | 0.331 | 0.409 | 0.627 |
| Q45EK1 | -1.38 | -0.226 | 0.186 | -1.54 | 0.0925 | 0.772 | 0.811 | 0.0644 |
| O01869 | -1.39 | -0.612 | 0.0526 | -0.828 | 0.000961 | 0.0867 | 0.876 | 0.0259 |
| Q9U2A8 | -1.4 | -1.08 | -0.305 | 0.219 | 0.000725 | 0.00499 | 0.361 | 0.51 |
| Q20384 | -1.4 | -1.26 | 0.172 | -1.21 | 0.0302 | 0.0479 | 0.771 | 0.0561 |
| O18236 | -1.43 | 0.138 | -0.286 | -0.853 | 0.113 | 0.872 | 0.74 | 0.331 |
| Q21832 | -1.44 | -1.48 | -0.0124 | -0.94 | 0.00268 | 0.00215 | 0.975 | 0.032 |
| Q9N492 | -1.44 | -1.71 | -0.44 | -2.34 | 0.0511 | 0.0242 | 0.526 | 0.00392 |
| Q93573 | -1.45 | -0.401 | -0.805 | -0.0279 | 0.0039 | 0.356 | 0.0762 | 0.948 |
| Q09979 | -1.45 | -1.24 | -0.793 | -1.28 | 0.0933 | 0.146 | 0.343 | 0.136 |
| Q19948 | -1.46 | -0.24 | 0.573 | -0.571 | 0.046 | 0.724 | 0.405 | 0.406 |
| Q21890 | -1.46 | -0.395 | 0.569 | -1.53 | 0.0298 | 0.523 | 0.362 | 0.0243 |

| | | | | | | | | |
|---|---|---|---|---|---|---|---|---|
| Q9N2W5 | -1.46 | -0.41 | 0.613 | -1.57 | 0.00942 | 0.412 | 0.226 | 0.00592 |
| Q1XFY9 | -1.46 | 0.9 | 0.0213 | 0.721 | 0.00683 | 0.0701 | 0.964 | 0.138 |
| Q20277 | -1.48 | -1.17 | 0.473 | -0.796 | 0.000807 | 0.00485 | 0.196 | 0.0385 |
| P17512 | -1.49 | -0.551 | 2.57 | -1.53 | 0.173 | 0.603 | 0.0263 | 0.161 |
| P83351 | -1.49 | -0.194 | 1.51 | -0.904 | 0.0151 | 0.724 | 0.0143 | 0.115 |
| Q9XVF7 | -1.49 | -0.611 | -0.446 | -0.124 | 0.000709 | 0.0975 | 0.216 | 0.723 |
| G5EES8 | -1.49 | -0.554 | -0.0388 | -1.35 | 0.0585 | 0.456 | 0.958 | 0.0832 |
| Q20603 | -1.5 | 0.407 | -0.302 | 2.2 | 0.0535 | 0.574 | 0.676 | 0.0079 |
| Q22054 | -1.51 | 0.391 | 0.446 | -0.0204 | 0.000397 | 0.251 | 0.193 | 0.951 |
| O17891 | -1.52 | 1.44 | -0.43 | 2.53 | 0.0422 | 0.053 | 0.537 | 0.00233 |
| Q09583 | -1.54 | -2.76 | -0.391 | -2.04 | 0.00548 | 4.04E-05 | 0.416 | 0.000656 |
| O16997 | -1.54 | -0.889 | 0.333 | -0.799 | 0.0305 | 0.187 | 0.611 | 0.232 |
| Q20034 | -1.55 | -0.523 | -0.189 | -0.854 | 0.0694 | 0.518 | 0.814 | 0.298 |
| C6KRJ5 | -1.57 | -0.996 | -0.631 | -1.85 | 0.0161 | 0.105 | 0.291 | 0.0062 |
| Q10906 | -1.57 | -0.83 | 0.504 | -1.31 | 0.119 | 0.395 | 0.602 | 0.189 |
| Q9XWU9 | -1.58 | -1.62 | 0.00368 | -0.898 | 0.000674 | 0.000562 | 0.992 | 0.0269 |
| A9UJN7 | -1.6 | -2.75 | 0.642 | -2.59 | 0.00591 | 7.25E-05 | 0.214 | 0.000129 |
| C1P629 | -1.6 | -0.126 | 0.635 | -1.05 | 0.0536 | 0.87 | 0.416 | 0.186 |
| Q19743 | -1.61 | -1.02 | 0.353 | -0.878 | 0.0522 | 0.202 | 0.649 | 0.266 |
| P49196 | -1.62 | -1.11 | 0.666 | -0.736 | 0.00145 | 0.0168 | 0.127 | 0.0946 |
| Q09248 | -1.62 | -0.81 | 0.369 | -1.98 | 0.0112 | 0.164 | 0.514 | 0.00303 |
| Q19972 | -1.62 | -1.33 | -0.581 | -1.88 | 0.0204 | 0.0508 | 0.364 | 0.00921 |
| H9G2S0 | -1.63 | -3.39 | -0.0863 | -1.67 | 0.00224 | 2.06E-06 | 0.846 | 0.00187 |
| P91277 | -1.64 | -2.71 | 0.329 | -0.94 | 0.0268 | 0.00115 | 0.627 | 0.178 |
| A8WHS3 | -1.64 | -0.0997 | -0.271 | -0.787 | 0.0128 | 0.865 | 0.644 | 0.192 |
| Q18494 | -1.65 | -2.09 | 0.329 | -1.32 | 0.0397 | 0.0126 | 0.659 | 0.0921 |
| P55326 | -1.66 | -4.33 | -0.00159 | -4.41 | 0.0172 | 6.42E-06 | 0.998 | 5.18E-06 |
| H2L2E8 | -1.66 | -1.46 | -1.2 | -1.75 | 0.0652 | 0.101 | 0.172 | 0.0538 |
| Q9BKU8 | -1.69 | -2.87 | 1.06 | -2.83 | 0.103 | 0.0106 | 0.295 | 0.0115 |
| Q9U3F4 | -1.69 | -2.63 | 0.708 | -1.92 | 0.000361 | 4.28E-06 | 0.0693 | 0.00011 |
| P90983 | -1.69 | -0.277 | 0.203 | -0.659 | 0.00487 | 0.591 | 0.694 | 0.213 |
| Q8WQA8 | -1.69 | -0.832 | -0.389 | -0.796 | 7.47E-05 | 0.0164 | 0.223 | 0.0206 |
| Q22615 | -1.69 | -0.866 | -0.136 | -1.78 | 0.133 | 0.427 | 0.899 | 0.115 |
| H2L045 | -1.7 | -0.581 | 0.798 | -0.875 | 0.00661 | 0.295 | 0.157 | 0.123 |
| A5PEX6 | -1.7 | -0.486 | 1.03 | -0.659 | 0.0172 | 0.451 | 0.122 | 0.311 |
| H2KYR1 | -1.71 | -1.3 | 0.314 | -1.49 | 0.00179 | 0.011 | 0.489 | 0.00476 |
| Q93618 | -1.71 | -1.24 | -0.461 | -1.51 | 0.0918 | 0.212 | 0.633 | 0.133 |
| P47208 | -1.72 | 0.466 | 0.465 | -0.0713 | 0.00192 | 0.32 | 0.32 | 0.877 |
| Q09237 | -1.73 | -2.33 | 0.523 | -1.41 | 0.0071 | 0.000832 | 0.357 | 0.0226 |
| Q19869 | -1.73 | 0.0732 | 0.437 | 0.443 | 0.000437 | 0.849 | 0.265 | 0.259 |
| Q9BL39 | -1.73 | -2.49 | -0.81 | -2.92 | 0.0268 | 0.00319 | 0.265 | 0.000937 |
| Q20898 | -1.73 | -0.603 | -0.692 | -1.48 | 0.0346 | 0.427 | 0.363 | 0.0647 |
| A0A486W TM4 | -1.73 | -1.24 | -0.813 | -1.12 | 0.105 | 0.236 | 0.429 | 0.281 |
| Q9U757 | -1.75 | -0.33 | -0.0106 | -1.28 | 0.00203 | 0.486 | 0.982 | 0.0149 |
| P34313 | -1.77 | -0.512 | 0.926 | -0.439 | 0.00618 | 0.366 | 0.114 | 0.438 |
| Q22600 | -1.77 | -0.558 | -0.557 | -1.45 | 0.00763 | 0.341 | 0.342 | 0.0233 |

| | | | | | | | | |
|---|---|---|---|---|---|---|---|---|
| G5EEJ7 | -1.78 | -0.659 | 0.921 | -0.984 | 0.00742 | 0.266 | 0.128 | 0.106 |
| Q9U2I0 | -1.79 | -1.02 | -0.209 | -0.365 | 0.0395 | 0.218 | 0.795 | 0.65 |
| O76357 | -1.8 | -0.542 | 1.71 | -1.2 | 0.00547 | 0.338 | 0.0074 | 0.0457 |
| Q20310 | -1.8 | -1.47 | 0.344 | -2.09 | 0.000239 | 0.00138 | 0.365 | 5.87E-05 |
| Q19877 | -1.82 | 0.277 | -0.655 | 1.01 | 0.000261 | 0.471 | 0.102 | 0.0173 |
| G5ECL3 | -1.84 | -1.28 | -0.31 | -1.19 | 0.0154 | 0.0763 | 0.649 | 0.0973 |
| A0A9J5D<br>XX3 | -1.85 | -3.38 | 0.245 | -1.46 | 0.0842 | 0.00438 | 0.809 | 0.164 |
| P48166 | -1.86 | -0.141 | -1.12 | 0.396 | 0.00906 | 0.822 | 0.0889 | 0.529 |
| P91353 | -1.86 | -0.304 | -1.44 | 0.463 | 0.0163 | 0.661 | 0.0523 | 0.506 |
| O44739 | -1.87 | -1.06 | -0.727 | -1.23 | 0.00192 | 0.0493 | 0.16 | 0.0252 |
| Q17935 | -1.87 | -1.58 | 0.256 | -0.444 | 0.00308 | 0.00895 | 0.631 | 0.409 |
| P91207 | -1.88 | -3.98 | 0.0132 | -2.33 | 0.000464 | 1.73E-07 | 0.975 | 6.44E-05 |
| Q20011 | -1.89 | -0.97 | -0.5 | -1.03 | 0.00533 | 0.113 | 0.398 | 0.0943 |
| Q20720 | -1.9 | -0.351 | 0.733 | -0.348 | 0.0467 | 0.693 | 0.414 | 0.695 |
| O01504 | -1.91 | -0.515 | 0.203 | -0.636 | 2.46E-05 | 0.116 | 0.521 | 0.0578 |
| Q93289 | -1.92 | -1.16 | -0.00579 | -1.05 | 0.0743 | 0.263 | 0.995 | 0.31 |
| P91306 | -1.93 | 2.97 | 1.43 | 2.55 | 0.0901 | 0.0144 | 0.199 | 0.031 |
| G5EC91 | -1.94 | -0.817 | -1.89 | -1.02 | 0.0225 | 0.299 | 0.0256 | 0.2 |
| Q21065 | -1.95 | -3.22 | -1.4 | -2.9 | 0.0305 | 0.00144 | 0.106 | 0.0031 |
| A7DTF5 | -1.95 | -1.54 | 0.849 | -1.49 | 0.000412 | 0.00262 | 0.064 | 0.00335 |
| Q20753 | -1.96 | -0.462 | -1.14 | -1.49 | 0.0158 | 0.527 | 0.133 | 0.0549 |
| P34383 | -1.97 | 1.06 | -1.2 | 0.896 | 0.00439 | 0.0895 | 0.0583 | 0.144 |
| A0A486W<br>Y88 | -1.97 | -0.532 | 0.154 | -1.46 | 0.00536 | 0.388 | 0.8 | 0.0282 |
| Q21693 | -1.98 | -0.961 | 1.67 | -1.14 | 0.0424 | 0.297 | 0.0806 | 0.219 |
| Q9BPN9 | -1.99 | -0.754 | -0.183 | -0.98 | 0.00831 | 0.264 | 0.782 | 0.153 |
| Q9N3G0 | -2 | -2.47 | -0.401 | -2.07 | 0.025 | 0.00799 | 0.623 | 0.0212 |
| Q21930 | -2.01 | -0.149 | -0.129 | -0.0554 | 0.000344 | 0.731 | 0.766 | 0.898 |
| P05690 | -2.03 | 0.589 | -0.271 | 1.14 | 2.95E-05 | 0.0992 | 0.43 | 0.00422 |
| H2KZ22 | -2.03 | -1.61 | -0.242 | -1.22 | 0.00717 | 0.0259 | 0.713 | 0.0796 |
| G5ECM6 | -2.04 | 0.34 | -0.621 | -1.22 | 0.0253 | 0.683 | 0.458 | 0.156 |
| Q9U3B7 | -2.04 | -2.69 | -0.788 | -2.27 | 0.0719 | 0.0228 | 0.465 | 0.0489 |
| Q22285 | -2.06 | -1.41 | 0.175 | -1.27 | 9.58E-05 | 0.00247 | 0.653 | 0.00504 |
| G5EF97 | -2.07 | -0.502 | 0.351 | -0.953 | 0.00173 | 0.362 | 0.52 | 0.0958 |
| Q9XVR8 | -2.08 | -2.21 | 0.461 | -2.58 | 0.00125 | 0.000765 | 0.385 | 0.000194 |
| Q23258 | -2.08 | -3.65 | 0.962 | -2.95 | 0.00206 | 1.23E-05 | 0.103 | 0.000108 |
| O45495 | -2.09 | -0.4 | 0.0973 | -1 | 0.014 | 0.599 | 0.898 | 0.198 |
| Q8MXR6 | -2.1 | -1.91 | 0.495 | -1.87 | 2.79E-05 | 7.33E-05 | 0.171 | 9.18E-05 |
| Q95XR1 | -2.1 | -1.22 | -0.156 | -1.82 | 0.00138 | 0.0358 | 0.771 | 0.00384 |
| O17687 | -2.11 | -0.0101 | 0.844 | -0.342 | 0.00382 | 0.987 | 0.187 | 0.583 |
| Q86NE0 | -2.14 | -1.22 | -0.127 | -0.328 | 0.00205 | 0.0483 | 0.825 | 0.57 |
| Q9N3F7 | -2.14 | -3.44 | -2.1 | -3.04 | 0.0305 | 0.00176 | 0.0335 | 0.00424 |
| G5EBY6 | -2.16 | -0.55 | 1.04 | -1.43 | 0.0051 | 0.411 | 0.13 | 0.0442 |
| G5ECC1 | -2.16 | -1.45 | 0.876 | -1.76 | 0.0551 | 0.181 | 0.41 | 0.111 |
| O02642 | -2.16 | -1.74 | -0.351 | -1.65 | 0.00021 | 0.00134 | 0.432 | 0.00199 |
| Q9NAF4 | -2.16 | -1.29 | 0.279 | -2.6 | 0.00537 | 0.069 | 0.676 | 0.00142 |

| | | | | | | | | |
|---|---|---|---|---|---|---|---|---|
| A0A3P6N624 | -2.19 | -1.47 | 2.5 | -1.74 | 0.221 | 0.406 | 0.166 | 0.326 |
| O61708 | -2.23 | -2.2 | 0.416 | -1.1 | 0.0272 | 0.029 | 0.652 | 0.243 |
| P41847 | -2.23 | -2.26 | 0.74 | -2.46 | 0.00091 | 0.000834 | 0.185 | 0.000412 |
| Q19694 | -2.23 | -2.02 | 0.889 | -1.87 | 0.00253 | 0.00505 | 0.165 | 0.0081 |
| Q18012 | -2.24 | -2.73 | 0.129 | -2.91 | 0.0175 | 0.00543 | 0.879 | 0.00349 |
| A0A1Q2U2F4 | -2.24 | -0.939 | -0.185 | -1.31 | 0.0138 | 0.256 | 0.819 | 0.122 |
| P52009 | -2.27 | -2.64 | -0.534 | -3.68 | 0.00379 | 0.00124 | 0.427 | 6.42E-05 |
| Q95XT5 | -2.27 | 0.62 | -2.59 | -0.386 | 0.0177 | 0.475 | 0.0084 | 0.654 |
| P34500 | -2.28 | -4.3 | 0.343 | -1.3 | 0.00247 | 7.31E-06 | 0.587 | 0.0544 |
| Q23543 | -2.29 | -3.22 | -0.292 | -3.16 | 0.000967 | 4.29E-05 | 0.602 | 5.26E-05 |
| P34714 | -2.3 | -1.93 | -0.55 | -1.84 | 0.0139 | 0.0335 | 0.512 | 0.0415 |
| Q9XTY3 | -2.3 | -1.03 | -1.87 | -2.1 | 0.00365 | 0.139 | 0.013 | 0.00654 |
| P49197 | -2.33 | -1.99 | 0.351 | -1.54 | 1.81E-05 | 8.87E-05 | 0.352 | 0.00086 |
| Q967F1 | -2.33 | -0.835 | -0.739 | -2.63 | 0.0121 | 0.319 | 0.376 | 0.00581 |
| Q9N590 | -2.33 | -3.47 | -3.83 | -3.9 | 0.00146 | 4.18E-05 | 1.53E-05 | 1.24E-05 |
| C6KRN4 | -2.34 | -1.34 | 1.25 | -0.422 | 0.00203 | 0.0483 | 0.0624 | 0.506 |
| Q20748 | -2.36 | -0.835 | -0.0906 | -1.87 | 0.0173 | 0.355 | 0.919 | 0.0499 |
| P91913 | -2.37 | -1.24 | 0.418 | -1.15 | 1.45E-05 | 0.00429 | 0.269 | 0.00673 |
| Q18421 | -2.37 | -1.62 | -0.612 | -1.96 | 0.101 | 0.251 | 0.658 | 0.169 |
| G5ED89 | -2.38 | -1.45 | -0.468 | -2.4 | 0.000788 | 0.0207 | 0.413 | 0.000739 |
| P43508 | -2.42 | -0.984 | 0.827 | 0.692 | 0.000205 | 0.0622 | 0.111 | 0.176 |
| O45713 | -2.42 | -1.25 | -0.11 | -0.503 | 1.63E-06 | 0.00111 | 0.725 | 0.121 |
| O45944 | -2.47 | -1.8 | 0.107 | 1.52 | 0.0329 | 0.105 | 0.919 | 0.165 |
| Q9BIB7 | -2.47 | -3.99 | 0.377 | -3.98 | 0.000104 | 6.02E-07 | 0.426 | 6.26E-07 |
| Q21004 | -2.49 | -3.54 | -0.0732 | -4.11 | 2.34E-05 | 4.62E-07 | 0.857 | 7.62E-08 |
| A5A8P5 | -2.5 | -0.938 | 0.31 | -1.36 | 0.00102 | 0.142 | 0.616 | 0.041 |
| Q93233 | -2.51 | -2.13 | -0.977 | -2.85 | 0.0791 | 0.13 | 0.473 | 0.0498 |
| Q95Y04 | -2.52 | -1.53 | 0.257 | -1.72 | 1.13E-05 | 0.00122 | 0.508 | 0.000458 |
| G5EEV5 | -2.53 | 1.28 | -0.819 | 1.12 | 0.000874 | 0.0504 | 0.193 | 0.0825 |
| P92186 | -2.59 | -1.27 | 0.316 | -1.48 | 0.000217 | 0.0293 | 0.554 | 0.013 |
| P48162 | -2.6 | -3.73 | 0.0785 | -2.96 | 8.00E-05 | 1.74E-06 | 0.87 | 2.18E-05 |
| Q20950 | -2.62 | 0.503 | -0.0308 | 0.474 | 0.000164 | 0.342 | 0.953 | 0.37 |
| Q8MPX7 | -2.62 | -2.43 | -0.696 | 0.392 | 0.0249 | 0.0356 | 0.515 | 0.712 |
| O45181 | -2.68 | -2.42 | 0.532 | -0.302 | 0.0184 | 0.0304 | 0.604 | 0.768 |
| Q9XVT0 | -2.69 | -1.42 | 0.746 | -2.39 | 3.21E-05 | 0.00667 | 0.117 | 0.000105 |
| B3GWA1 | -2.69 | -2.57 | 0.116 | -2.88 | 0.00247 | 0.00348 | 0.876 | 0.00151 |
| B1Q273 | -2.74 | -2.35 | 1.55 | -3.29 | 0.00256 | 0.00718 | 0.057 | 0.000622 |
| Q22336 | -2.76 | -1.74 | 0.512 | -0.897 | 0.000126 | 0.00512 | 0.346 | 0.11 |
| O62388 | -2.8 | -1.95 | 0.604 | -1.31 | 2.55E-05 | 0.000756 | 0.204 | 0.0123 |
| Q09365 | -2.82 | -1.15 | 0.639 | -0.73 | 0.000547 | 0.089 | 0.328 | 0.267 |
| Q27488 | -2.83 | -2.36 | 0.276 | -2.33 | 9.60E-06 | 6.30E-05 | 0.519 | 7.12E-05 |
| Q22836 | -2.84 | -2.82 | -0.217 | -3.44 | 4.81E-05 | 5.15E-05 | 0.663 | 6.41E-06 |
| O45712 | -2.86 | -2.31 | -0.058 | 0.886 | 0.00893 | 0.0283 | 0.952 | 0.363 |
| Q19162 | -2.87 | -2.82 | 0.0116 | -2.87 | 0.0612 | 0.0659 | 0.994 | 0.0613 |
| Q9XWH0 | -2.89 | -1.33 | -1.54 | -2.29 | 0.0104 | 0.194 | 0.137 | 0.0344 |

| | | | | | | | | |
|---|---|---|---|---|---|---|---|---|
| Q9TYK1 | -2.9 | -3.97 | 0.645 | -4.3 | 9.39E-05 | 3.45E-06 | 0.247 | 1.39E-06 |
| P92199 | -2.94 | -2.33 | -1.32 | -1.92 | 0.0131 | 0.041 | 0.221 | 0.0845 |
| Q94051 | -2.94 | -3.29 | 0.118 | -2.43 | 0.000454 | 0.000162 | 0.857 | 0.00205 |
| Q9U256 | -2.95 | -2.56 | -0.329 | -2.9 | 0.000409 | 0.0013 | 0.614 | 0.000477 |
| Q22666 | -2.95 | -2.01 | -2.66 | -2.57 | 0.00584 | 0.0439 | 0.0109 | 0.0134 |
| Q966I7 | -2.99 | -3.09 | 1.31 | -3.26 | 0.00256 | 0.00197 | 0.129 | 0.00131 |
| Q9U3I0 | -3.05 | -0.849 | 0.283 | 0.532 | 0.000813 | 0.256 | 0.699 | 0.47 |
| Q9XU56 | -3.09 | -1.97 | -1.38 | -2.24 | 0.00059 | 0.0134 | 0.0681 | 0.00623 |
| Q95YC7 | -3.1 | -1.56 | 1.84 | -2.04 | 1.93E-05 | 0.00652 | 0.00217 | 0.000961 |
| O62337 | -3.13 | -2.54 | 1.42 | -2.92 | 0.0183 | 0.0483 | 0.245 | 0.0259 |
| O16291 | -3.14 | -1.12 | -1.47 | -1.7 | 0.01 | 0.308 | 0.185 | 0.129 |
| P91156 | -3.17 | -1.92 | -0.103 | -2.34 | 0.024 | 0.146 | 0.936 | 0.0825 |
| Q27535 | -3.2 | -1.38 | -0.0672 | -0.949 | 3.18E-06 | 0.00618 | 0.877 | 0.0432 |
| Q9TVW5 | -3.2 | 0.311 | 0.841 | 0.838 | 2.15E-05 | 0.552 | 0.121 | 0.122 |
| G5EBY3 | -3.22 | -2.34 | 1.12 | -2.75 | 0.0633 | 0.164 | 0.492 | 0.106 |
| O17570 | -3.25 | -2.87 | 0.775 | -2.76 | 1.95E-05 | 6.83E-05 | 0.152 | 0.000101 |
| P91250 | -3.26 | -2.89 | 1.03 | -2.58 | 2.40E-05 | 7.94E-05 | 0.0693 | 0.000236 |
| P45895 | -3.29 | 0.809 | 0.0636 | 0.9 | 2.92E-05 | 0.156 | 0.908 | 0.117 |
| Q17684 | -3.37 | -2.53 | 0.208 | -2.2 | 5.85E-07 | 1.46E-05 | 0.6 | 6.27E-05 |
| Q86FL8 | -3.39 | -3.58 | 0.232 | -0.265 | 0.00917 | 0.00663 | 0.839 | 0.816 |
| Q93805 | -3.43 | -2.97 | 0.455 | -3.43 | 1.25E-05 | 5.60E-05 | 0.395 | 1.23E-05 |
| Q09250 | -3.53 | -1.63 | 0.0387 | -2.82 | 2.54E-07 | 0.000774 | 0.92 | 3.47E-06 |
| D5MCQ2 | -3.64 | -1.86 | 0.82 | -3.15 | 7.47E-07 | 0.000685 | 0.0765 | 3.97E-06 |
| G5ECR7 | -3.86 | -2.36 | 0.329 | -2.7 | 4.44E-05 | 0.00308 | 0.626 | 0.00112 |
| P37165 | -3.96 | -3.44 | 1.9 | -3.55 | 1.92E-05 | 8.05E-05 | 0.00874 | 5.86E-05 |
| Q09958 | -4.07 | -2.5 | 0.969 | -2.91 | 5.96E-07 | 0.000112 | 0.0582 | 2.45E-05 |
| Q20311 | -4.27 | -3.38 | 4.69 | -3.87 | 0.00668 | 0.0245 | 0.00359 | 0.012 |
| O62477 | -4.38 | -3.4 | -0.775 | -3.75 | 0.0194 | 0.0594 | 0.647 | 0.0399 |
| Q94053 | -4.41 | -0.894 | 0.479 | -0.233 | 1.77E-06 | 0.132 | 0.405 | 0.683 |
| Q19969 | -4.46 | -4.03 | -1.34 | -4.25 | 1.41E-05 | 4.02E-05 | 0.0694 | 2.31E-05 |
| Q09254 | -4.49 | -3.88 | -1.66 | -3.2 | 3.00E-04 | 0.00103 | 0.0978 | 0.00422 |
| Q10033 | -4.51 | -4.48 | 0.226 | -3.87 | 2.66E-08 | 2.91E-08 | 0.584 | 1.75E-07 |
| Q9U332 | -4.54 | 0.794 | -2.66 | -1.22 | 0.0414 | 0.7 | 0.208 | 0.555 |
| Q18886 | -4.81 | -3.72 | -0.4 | -4.11 | 8.96E-07 | 1.56E-05 | 0.497 | 5.28E-06 |
| P34460 | -4.89 | -2.65 | -0.0229 | -3.82 | 3.29E-05 | 0.0057 | 0.978 | 0.000347 |
| A0A0K3AYJ1 | -5.38 | 0.462 | 0.428 | 0.212 | 1.23E-07 | 0.41 | 0.444 | 0.702 |
| G5EGS3 | -5.83 | -5.11 | -0.838 | -5.62 | 4.45E-07 | 2.04E-06 | 0.222 | 6.80E-07 |
| O62053 | -8.81 | -3.81 | -0.375 | -2.5 | 1.49E-11 | 6.42E-07 | 0.41 | 6.29E-05 |
